# ZNF16 is a nucleolar-associated protein that regulates expression of the rDNA and cancer-associated genes

**DOI:** 10.1101/2025.10.16.682606

**Authors:** Chelsea L. George, Laura A. Espinoza Quevedo, Jason Paratore, Matthew J. Alcaraz, Arlene P. Levario, Yasir Rahmatallah, Galina V. Glazko, Nathan S. Reyna, Laura A. Diaz-Martinez

**Affiliations:** Department of Biological Sciences, The University of Texas at El Paso, El Paso TX 79968; Department of Biology, Gonzaga University, Spokane WA, 99258; Department of Biomedical Informatics, University of Arkansas for Medical Sciences, Little Rock AR, 72205; Department of Biology, Ouachita Baptist University, Arkadelphia AR 71998

**Keywords:** zinc-finger proteins, nucleolar function, rDNA transcription, rRNA expression

## Abstract

ZNF16 (also known as HZF1 and KOX9) is a multi-C2H2 zinc finger protein first identified via its expression in human T-cells and shown to have a role in blood cell differentiation. ZNF16 was later shown to be ubiquitously expressed in a variety of fetal and adult tissues, suggesting a broader function. In this study, we confirm the ubiquitous expression of ZNF16 in a variety of cancer and non-cancer cell lines and show that ZNF16 depletion reduces cell viability in all cell lines tested. Furthermore, we show that ZNF16 localizes to the nucleolus in a transcription-dependent manner, interacts with the intergenic spacer region of the rDNA and promotes rDNA transcription. Additionally, RNA-seq experiments after ZNF16 depletion revealed that ZNF16 also has roles in a variety of pathways including ECM-receptor interaction, focal adhesions, cytokine-cytokine receptor interactions, human papillomavirus (HPV) infection and cancer pathways. These findings are consistent with broader roles for ZNF16, including the regulation of nucleolar function, a process that is essential for all cells, and provide evidence at the cellular/molecular level of its role in the regulation of cancer-associated genes (e.g., NRAS, BIRC3, EGFR).

## Introduction

Zinc finger proteins represent 5-10% of the human proteome (Andreini et al., 2006; Vilas et al., 2018), with ∼700 of these proteins belonging to the C2H2 zinc finger family (Schmitges et al., 2016), which comprises almost 50% of all putative transcription factors in vertebrates. These proteins typically consist of tandem arrays of zinc finger motifs that provide DNA-binding specificity and effector motifs (e.g., KRAB, SCAN) that mediate interaction with transcriptional activators or inhibitors (Schmitges et al., 2016). Although the most studied role of zinc finger motifs is in DNA binding (Klug and Rhodes, 1987), zinc finger motifs can also interact with RNA (Laity et al., 2001), proteins (Gamsjaeger et al., 2007) and lipids (Laity et al., 2001), indicating a wide diversity of functions for these types of proteins.

ZNF16 (also called HZF1) is a C2H2 zinc finger protein that has been associated with cancer progression (Ahn et al., 2020; Lee et al., 2021; Zhang et al., 2020). ZNF16 expression was positively associated with histological grade, shorter survival, and increased risk of relapse in gallbladder carcinoma patients (Ahn et al., 2020). Genetic changes in ZNF16 have also been associated with tongue squamous cell carcinoma (TSCC), with 35% of TSCC samples having ZNF16 amplification and 5% missense mutations (Zhang et al., 2020). Consistent with a role for ZNF16 in TSCC, ZNF16 depletion reduces TSCC cell viability and xenograft tumor size (Zhang et al., 2020).

ZNF16 has fifteen C2H2 zinc finger motifs in the C-terminus (Deng et al., 2010; Schmitges et al., 2016) and is localized in a zinc finger gene cluster on human chromosome 8q24.3 that contains seven zinc finger proteins (Lorenz et al., 2010). Interestingly, ZNF16 is the only protein in the cluster that lacks a Krüppel-associated box (KRAB) domain (Lorenz et al., 2010), a domain commonly associated with transcriptional repressors (Urrutia, 2003). Consistent with a role in transcription, ZNF16 is localized in the nucleus, activates transcription of a GAL1-lacZ reporter in yeast, and contains a transactivation domain in its unstructured N-terminal region that can activate transcription in yeast when fused to the GAL4-DNA binding domain(Deng et al., 2010).

ZNF16 was first identified via its expression in human T-cells (Thiesen, 1990). It was later shown to have a role in in vitro erythroid and megakaryocytic differentiation in vitro (Peng et al., 2006), and its overexpression increases cell proliferation and moderately reduces apoptosis induced by sodium arsenate (Li et al., 2011). However, the molecular mechanisms behind these functions are unclear. ZNF16 interacts with the CDK1-inhibitor INCA (Li et al., 2011) and binds upstream of the c-KIT promoter (Chen et al., 2014) in K562 cells, but ZNF16 overexpression has only minimal effects on cell cycle progression (Li et al., 2011).

In addition to its expression in blood cells, ZNF16 has also been shown to be expressed in a variety of human tissues including fetal brain, testis, cerebellum, and kidney (Lorenz et al., 2010), as well as adult brain, heart, skeletal muscle, liver, and bone marrow (Peng et al., 2006). ZNF16 interacts with proteins associated with a variety of functions including linker histones (Zhang et al., 2016), transcriptional regulators (e.g.,HMGA1, DCAF7, TRIM28, HDAC1), cell signaling (e.g., MAPK1, PPP2A,GRB2), ubiquitination and protein degradation (e.g., TRIM27, USP11, PSMC3), DNA replication and repair (e.g., LIG3, MCM6), proteins found in nuclear bodies (e.g., Coilin, PML), and proteins found at the nucleolus and/or involved in rRNA processing (e.g., UBTF, TCOF1, KNOP1, NOL9) (Schmitges et al., 2016).

Nucleoli are membraneless and highly dynamic organelles that contain over 800 proteins (Jarboui et al., 2011). They are organized around nucleolar organizing regions (NORs) that consist of multiple tandem repeats of the rDNA gene, separated by intergenic spacers (IGS) (Potapova and Gerton, 2019; Trinkle-Mulcahy, 2018). Nucleoli are the site of rDNA transcription, rRNA processing, and ribosome biogenesis, as well as a cellular stress response center (González-Arzola, 2024; Hua et al., 2022; Núñez Villacís et al., 2018). Nucleoli number, size and activity are increased in hyperproliferative cells, serving as a prognostic marker for tumor malignancy (Trinkle-Mulcahy, 2018).

Given the expression of ZNF16 in multiple human tissues (Lorenz et al., 2010; Peng et al., 2006) and its association with a variety of proteins, including nucleolar proteins (Schmitges et al., 2016), we hypothesized that ZNF16 has a more universal role that is relevant for cells in all these tissues in addition to its specific role in blood cell differentiation (Peng et al., 2006). Here, we describe a novel role for ZNF16 at the nucleolus and its association with changes in expression of a variety of genes involved in cancer pathways.

## Materials and Methods

### Cell culture & drug treatments

U2OS (ATCC), HeLa Tet-On, DLD1, RPE1-hTERT, and Hct116 cells (kind gift from H. Yu) were grown in a humidified incubator at 37°C with 5% CO_2_ in Dulbecco’s modified Eagle’s medium with 4.5 g/L glucose, L-glutamine, and pyruvate (DMEM; Corning), supplemented with 10% fetal bovine serum (FBS; HyClone). Drug treatments: Actinomycin-D (40 μM), cisplatin (20 μM), Camptothecin (10 μM), and DMSO.

### Plasmids and siRNA transfections

Plasmid pIRESpuro-EGFP-ZNF16 (EGFP-ZNF16) was generated by PCRing the open reading frame (ORF) and 3’UTR of ZNF16 from a pBeloBAC11containing the full ZNF16 gene (CTD-2012A17; Invitrogen), adding the FseI and AscI sites for subcloning into pIRESpuro-EGFP (EGFP; kind gift from H. Yu). The pIRESpuro-3xEGFP-ZNF16 plasmid (3xEGFP-ZNF16) was generated by subcloning of the ZNF16 gene from the pIRESpuro-EGFP-ZNF16 plasmid using FseI/AscI. Plasmids pHrD-IRES-Luc (human rRNA promoter-luciferase reporter) and pIRES-Luciferase were kind gifts from K. Ghoshal (Ghoshal et al., 2004) and Z. Karamysheva, respectively. Plasmids were transfected at a final concentration of 0.4 ng/ul using Lipofectamine 2000 (Life Technologies). The stably transfected 3xEGFP-ZNF16 cell line was produced by transfection of the pIRESpuro-3xEGFP-ZNF16 plasmid in HeLa Tet-On cells, selection with 0.5 ug/mL puromycin, clone picking and confirmation of gene expression via microscopy.

The sequences of the siRNAs are as follows: siControl (AccuTarget Negative Control siRNA (BioRP SN-1002, VWR 95030-562), siZNF16-E 5’-AAACUAUGCUGGUGAUGUU-3’ (Dharmacon), and siZNF16-3UTR 5’-UGACGUUUGGUUUGAGAUA-3’ (ON-TARGET Plus, Horizon Discovery). siRNAs were transfected at a final concentration of 10nM using Lipofectamine RNAiMAX (Life Technologies) according to manufacturer recommendations.

### Cell Viability Assays

Cells were transfected with the indicated siRNAs for 72h, then incubated with OZBlue (OZ Biosciences) for at least 30 min and fluorescence was measured using a FLx800 plate reader (BioTek). Background fluorescence from blank wells was subtracted and fluorescence was normalized to untreated cells.

### Western Blot

Cells were washed in PBS and lysed in lysis buffer (250mM SDS, 82.5mM Tris Base, 30% glycerol, 1.5mM bromophenol blue, and 43.7mM DTT), sonicated, and boiled for 10 minutes. Samples were separated by SDS-PAGE and transferred to a nitrocellulose membrane (GE Healthcare Life Sciences) using a semi-dry transfer apparatus (BioRad). The membranes were incubated with primary antibodies against ZNF16 (Rabbit anti-ZNF16 at 1:50; Sigma-Aldrich) and tubulin (Mouse anti-tubulin ascites at 1:100; kindly provided by S. Roychowdhury) dissolved in 5% non-fat dry milk in TBS-T (1X TBS + 0.01% Tween) overnight at 4°C. Membrane was then incubated with secondary antibodies anti-Rabbit IRDye-680RD (LICOR, 1:2000) and goat Anti-Mouse IgG (H&L) Horseradish peroxidase (HRP) conjugated antibody (ImmunoReagents, 1:10,000) at room temperature for 30 minutes, followed by incubation with Clarity Western Luminol/Enhancer Reagent (BioRad) for visualization of the HRP-conjugated antibody. The membranes were imaged using a LI-COR Odyssey imager.

### Immunostaining

Cells were cultured in 8-well chamber slides (Nunc Lab-Tek II or Falcon) and treated as described. Cells were fixed with 4% paraformaldehyde (PFA) for 20 minutes at room temperature. Primary antibodies were diluted in blocking buffer (0.2% Triton X-100 in PBS with 3% BSA) and incubated at 4°C overnight. Secondary antibodies were diluted in blocking buffer and incubated at room temperature, in the dark for 30 minutes. Cells were washed with 0.2% Triton X-100 in PBS, counterstained with DAPI, and mounted using Vectashield anti-fade mounting media (VectorLabs). Primary antibodies used: Rabbit anti-ZNF16 (1:50; Sigma-Aldrich), rabbit anti-GFP (1:500; Novus), mouse anti-Ki67 (1:250; BD Biosciences), mouse anti-γH2AX (1:200; BD Biosciences), rabbit anti-UBF (1:400, Novus), mouse anti-FBL (1:400, Novus), and mouse anti-NPM1 (1:800, Protein Tech). Secondary antibodies used: Alexa Fluor 488 donkey anti-rabbit IgG, Alexa Fluor 568 donkey anti-mouse IgG, and donkey anti-mouse Alexa Fluor 647 IgG (1:500, Life Technologies).

### 5-EU incorporation assay

Cells were transfected with siRNAs for 72h, then incubated with 0.5 mM 5’-Ethynyl uridine (5’-EU) for an hour. Next, cells were fixed with either 4% PFA or cold methanol and stained via click reaction with a fluorescent azide using the RNA Synthesis Assay Kit (ab228561, Abcam) or the Click-IT RNA AlexaFluor 594 RNA synthesis imaging kit (Invitrogen), according to manufacturer’s instructions. Cells were then washed overnight with 0.2% Triton X-100 in PBS, counterstained with DAPI, and mounted using Vectashield anti-fade mounting media (VectorLabs).

### Confocal Microscopy & Image Analysis

Samples were imaged with an LSM 700 confocal microscope (Zeiss, New York, New York, USA), equipped with an EC Plan-NEOFLUAR 63x/1.25 N.A. oil immersion objective and ZEN 2009 software (Zeiss), or a TCS SPE-II confocal microscope (Leica) equipped with the following ACS APO oil immersion objectives: 40X/1.15 N.A. Oil CS; 63X/1.30 N.A. and LASX software. At least 12 z-stacks were acquired per field. Images were then semi-automatically processed in ImageJ using a macro for z-stack projection (kind gift from B. Bell). Nuclei and nucleoli were segmented and quantified using CellProfiler (Stirling et al., 2021). The data was analyzed using excel, R-studio and/or JMP.

### Chromatin Immunoprecipitation (ChIP)

HeLa Tet-On and EGFP-ZNF16 HeLa Tet-On cells were crosslinked with 1% formaldehyde, collected, washed, and lysed, then the nuclei were isolated and the chromatin immunoprecipitated with 5 ug of antibodies against UBF, GFP, or control IgG using a Magnetic ChIP Kit (ThermoScientific Pierce) according to the manufacturer protocol. After DNA elution from the beads, the immunoprecipitated rDNA was quantified by qPCR using the Power up SYBR green mastermix (Applied Biosystems) and previously described primers spanning the rDNA repeat (Zhu et al., 2010).

### qPCR

Total RNA was extracted from cells using TRIzol with the Direct-zol RNA Miniprep Kit (Zymo Research) according to the manufacturer’s protocol. RNA extracts were reverse transcribed using the High-Capacity cDNA Kit (Applied Biosystems). cDNA samples were subjected to qPCR in an Applied Biosystems 7900HT qPCR, using either TaqMan probes and the TaqMan Gene Expression Master Mix (ThermoFisher Scientific), or DNA primers and the Luna Universal probe qPCR mastermix (M3004, New England Biosciences).

### RNA-seq

Cells were treated with siControl and siZNF16-E siRNAs for 72h, then lysed in TRIzol, frozen and shipped to the University of Arkansas for Medical Sciences (UAMS) Genomics Core Facility. Triplicate samples per siRNA were analyzed. Quality control for the 76 base pairs single-end raw reads was ensured using *Trimmomatic* (Bolger et al., 2014) to perform the following steps: 1) remove Illumina adapter and PCR primer sequences, 2) remove leading and trailing bases with low quality, 3) scan reads with a 4-base wide sliding window and cut when the average quality score per base drops below 15, and 4) drop reads shorter than 36 bases long. Reads surviving the quality control criteria were aligned to the human genome model hg19 using *Tophat* (Trapnell et al., 2012), allowing two mismatches. Alignments were quantified per gene using *featureCounts* from package *Subread* (Liao et al., 2014). Genes with zero counts in 5 or more out of 6 samples were deemed unexpressed and discarded, leaving 30,245 expressed genes. Initial differential expression (DE) analysis with paired-sample design was performed using Wald test from Bioconductor package DESeq2 (Love et al., 2014). The RNA-seq read summary and the full results from DESeq2 are provided in the Supplemental Table. RNA-seq results where further analyzed using iPathwayGuide (AdvaitaBio) as follows: Differentially expressed genes were identified as those with an adjusted p-value <0.05 and log_2_ fold change of >0.6 and the data were analyzed for enrichment of metabolic pathways and diseases using the Kyoto Encyclopedia of Genes and Genomes (KEGG) database (Release 96.0+/11-21, Nov20), gene ontologies from the Gene Ontology Consortium database (2020-Oct14), miRNAs from miRbase (MIRBASE Version 22.1, 10/18) and the TARGETSCAN database (human version 7.2), network of regulatory relations from BioGRID (v4.0.189, Aug 25^th^ 2020), chemical/drugs/toxicants from the Comparative Toxiccogenomics Database (July 2020). The full iPathwayGuide report, containing the full methodology for pathway analysis and results is provided in the supplemental materials.

### DNA damage assays

HeLa or U2OS cells were transfected with the indicated plasmids as described in the plasmid and siRNA transfections section. Seventy-two hours after transfection, the cells were treated with 20 μM cisplatin,10 μM Camptothecin, or DMSO (control) for four hours, then fixed and stained with yH2AX antibody as described above. Irradiation of cells with X-rays was carried out with a dose of 5Gy by use of a X-RAD 160 (Precision X-Ray Inc. N. Brandford, CT USA).

## Results

### ZNF16 is expressed and promotes cell viability in a variety of cell lines

ZNF16 has been reported to be expressed in a variety of human fetal and adult tissues (Lorenz et al., 2010; Peng et al., 2006). However, all studies on ZNF16 function at the cellular/molecular level have been performed in K652 leukemia cells (Chen et al., 2014; Li et al., 2011; Peng et al., 2006) or in cells expressing exogenous ZNF16 (Deng et al., 2010). To begin our study of ZNF16 functions, we first tested a panel of five non-blood cell lines for ZNF16 expression by qPCR using probes against the ZNF16 mRNA and GAPDH as internal control. ZNF16 was expressed in all five cell lines tested (Fig. 1A). Relative expression was normalized to HeLa Tet-On, which was the lowest expressing cell line. Remarkably, there was a wide range of ZNF16 expression with the osteosarcoma cell line U2OS having a 30-fold higher expression than HeLa. The two colon cancer cell lines (Hct116 and DLD1) and the non-cancer RPE1-hTERT cell line (RPE1) had intermediate levels of expression. These results indicate that ZNF16 is ubiquitously expressed in both cancer and non-cancer cell lines, suggesting that it has other potential functions, in addition to blood cell differentiation.

**Figure 1.**
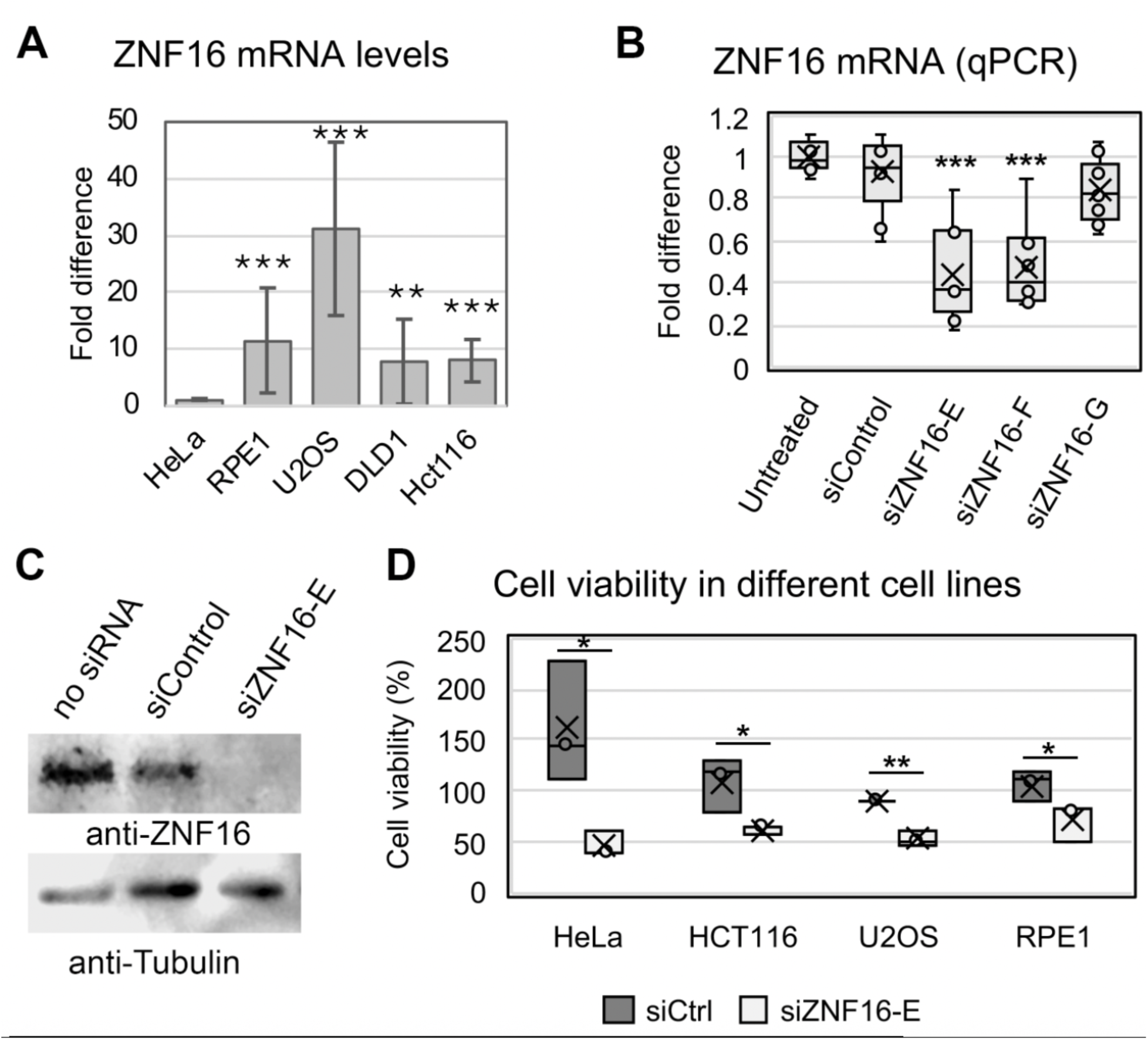
ZNF16 is expressed and promotes cell viability in different cell lines. (A) Quantification of ZNF16 mRNA by qPCR in the indicated cell lines. (B) Quantification of ZNF16 mRNA by qPCR in U2OS treated with the indicated siRNAs for 72h and (C) corresponding western blot. (D) Cell viability of indicated cell lines 72h after transfection with the indicated siRNAs. Results from each treatment were normalized to the corresponding untreated sample (100% viability, not shown). Bars represent the average of three biological repeats. Error bars represent standard deviations. Asterisks denote p-values as follows: *p<0.05, **p<0.01, ***p<0.001.

To begin addressing the role of ZNF16, three independent siRNAs were transfected into the highly expressing U2OS cell line, and their knockdown efficiency was quantified by qPCR. siZNF16-E most consistently reduced ZNF16 siRNAs compared to the other two siRNAs (Fig. 1B), and the effectiveness of the knockdown was confirmed by western blot (Fig. 1C). Thus, siZNF16-E was used for all subsequent experiments.

To test whether ZNF16 depletion affected cell viability, a sub-panel of cell lines was transfected with a non-targeting siRNA (siControl or siCtrl) or siZNF16-E siRNAs. Hct116 was included while DLD1 was omitted for simplicity, given that both are colon cancer cell lines and express similar levels of ZNF16 (Fig. 1A). Cell viability was markedly reduced in all cell lines treated with siZNF16-E compared to siControl. Note that the samples are normalized to an untreated sample, which is why the siControl samples are not all at 100% viability.

### ZNF16 is enriched at the nucleoli

To begin studying ZNF16 function, we first asked where ZNF16 is located in cells. First, we visualized ZNF16 localization by transient transfection of plasmids containing EGFP (control), EGFP-ZNF16 or 3xEGFP-ZNF16 in HeLa and U2OS cells, followed by immunostaining with antibodies against the nucleolar protein Ki67 (Fig. 2A-B). Although all exogenous forms of ZNF16 localize to the nucleus, the exact localization within subnuclear structures varies depending on the tag: 3xEGFP-ZNF16 is localized to the nucleoli in both cell lines, while the localization of EGFP-ZNF16 varies by cell line. EGFP-ZNF16 localizes to the nucleoplasm in both cell lines and is enriched in the nucleoli in U2OS cells but not in HeLa cells. These results indicate that the size of the N-terminal tag might affect ZNF16 folding and localization. Furthermore, transient transfection of 3xEGFP-ZNF16 also resulted in two main patterns of localization: A majority of the cells (71.5% on average from four independent experiments, Fig. 2C) showed 3xEGFP-ZNF16 enrichment in the nucleoli with little localization to the nucleoplasm, while the remaining cells showed both nucleolar localization and localization to other subnuclear structures (freckled localization in Fig. 2C). These results indicate that exogenous ZNF16 localization is influenced by both the tag size and the level of expression.

**Figure 2.**
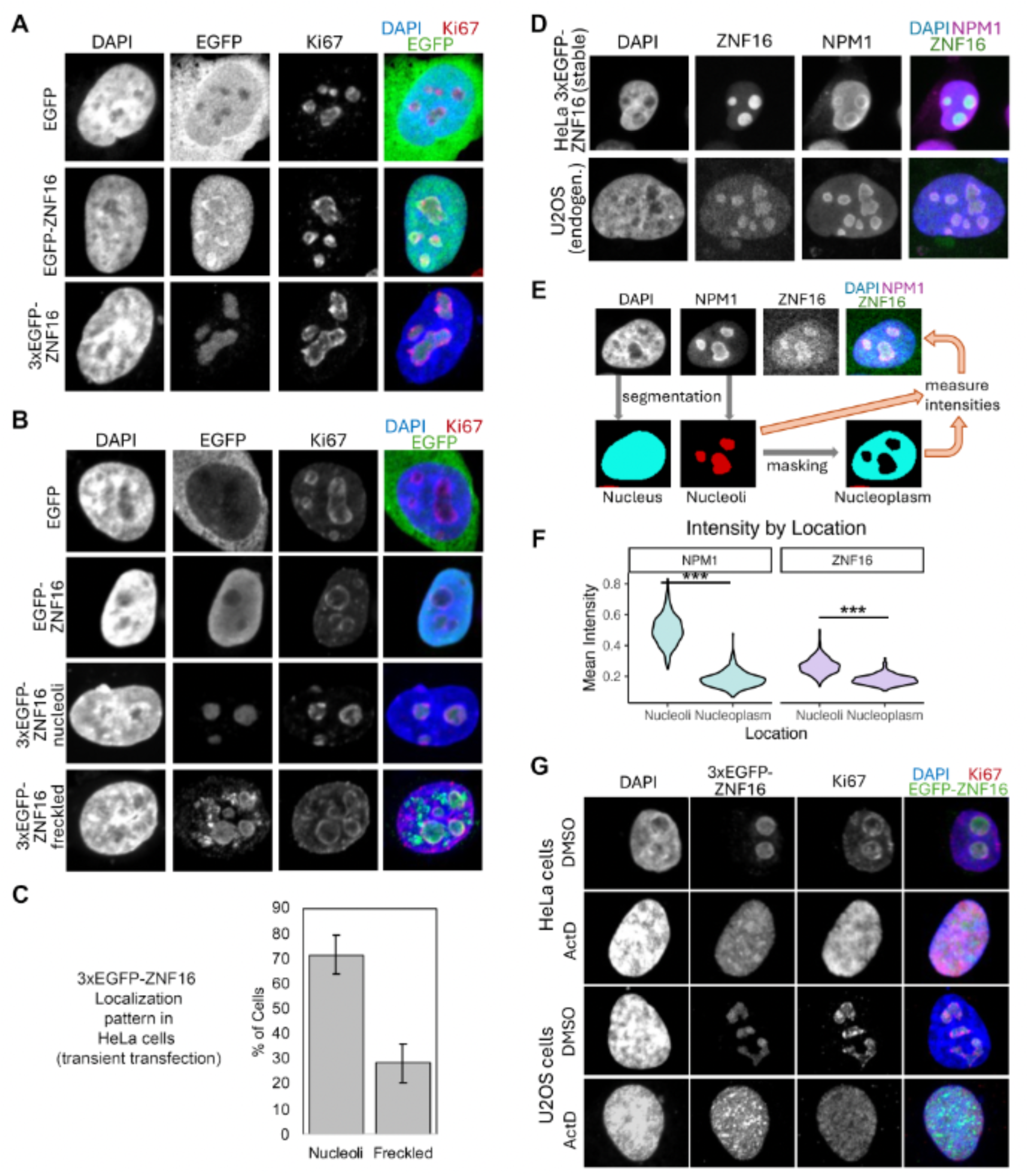
ZNF16 is enriched at the nucleoli. (A-B) Representative micrographs of U2OS (A) and HeLa (B) cells transiently transfected with the plasmids containing EGFP, EGPF-ZNF16, or 3xEGFP-ZNF16, and immunostained with the indicated antibodies. (C) Quantification of the localization patterns after transfection of 3xEGFP-ZNF16 in HeLa cells. (D) Representative micrographs of HeLa cells stably transfected with 3xEGFP-ZNF16 (top row) and U2OS cells (bottom row) immunostained with the indicated antibodies. (E) Schematic showing the image analysis performed in CellProfiler to measure the mean intensity of ZNF16 and NPM1 signal in the nucleoli and nucleoplasm at the single cell level. Cells were segmented in the DAPI and NPM1 channels to obtain masks for the nucleus and nucleoli, respectively. The nucleoli masks were then subtracted from the nuclei masks to obtain a nucleoplasm mask. Mean intensities were then measured using the nucleoli and nucleoplasm masks per cell. (F) Violin plot showing the mean intensity signal of NPM1 and ZNF16 in the nucleoli vs nucleoplasm (n = 507 cells). Asterisks denote p-values as follows: *p<0.05, **p<0.01, ***p<0.001. (G) Representative micrographs of cells transfected with 3xEGFP-ZNF16 plasmid, incubated for 24 h, then incubated with 40 μM actinomycin D for 4h before fixation and immunostaining with the indicated antibodies.

In order to avoid potential artifacts due to protein overexpression, we obtained a HeLa cell line stably transfected with the 3xEGFP-ZNF16 plasmid. This cell line expresses 3xEGFP-ZNF16 at lower levels than transient transfection (data not shown). Immunostaining of this stable cell line with antibodies against the nucleolar protein nucleophosmin 1 (NPM1, Fig. 2D top row) shows enrichment of 3xEGFP-ZNF16 at the nucleoli with little to no “freckles” in the nucleoplasm. Lastly, immunostaining of endogenous ZNF16 in U2OS cells with anti-ZNF16 antibodies shows a similar pattern of nucleolar enrichment (Fig. 2D bottom row). To quantify the intensity of ZNF16 in the nucleoli vs the nucleoplasm, U2OS cells immunostained with anti-ZNF16 and anti-NPM1 antibodies were segmented and analyzed using CellProfiler (Stirling et al., 2021). First, the nucleus was segmented in the DAPI channel, followed by segmentation of the nucleoli in the NPM1 channel. Then, the nucleoli regions were subtracted from the nucleus by generating a nucleoplasm mask. The mean intensity of NPM1 and ZNF16 was measured using the nucleoli and the nucleoplasm masks at the single cell level (see schematic in Fig. 2E). This analysis revealed that the mean ZNF16 signal is significantly higher in the nucleoli than the nucleoplasm (Fig. 2F), following a pattern similar to NPM1, a protein that is known to be localized to the nucleoli (Schmidt-Zachmann et al., 1987).

Lastly, HeLa and U2OS cells transiently expressing 3xGFP-ZNF16 were incubated with the RNApol-I inhibitor actinomycin-D to test the effect of nucleolar transcription inhibition on ZNF16 localization. After treatment with actinomycin-D, ZNF16 and the nucleolar protein Ki67 relocalize from the nucleolus to the nucleoplasm. Taken together, these results indicate that ZNF16 and exogenously expressed 3xEGFP-ZNF16 are enriched in the nucleolus, and their nucleolar localization is disrupted after inhibition of RNApol-I transcription.

### ZNF16 regulates rDNA expression

Given that both endogenous ZNF16 and 3xEGFP-ZNF16 localize to the nucleus and are enriched in the nucleoli compared to the nucleoplasm (∼1.5-fold for endogenous ZNF16), we next asked whether ZNF16 regulates nucleolar function by evaluating its role in rDNA expression. First, we tested whether transient expression of exogenous ZNF16 impacts transcription of an rDNA luciferase reporter. HeLa and U2OS cells were transiently co-transfected with plasmids containing EGFP-ZNF16 or 3xEGFP-ZNF16 and the pHrD-IRES-Luc plasmid (Ghoshal et al., 2004), which contains the human rDNA promoter fused to a luciferase reporter. Expression of 3xEGFP-ZNF16, which was shown to be enriched at the nucleolus (Fig. 2A-B), significantly increased luciferase activity in both U2OS and HeLa cells (Fig. 3A-B). In contrast, EGFP-ZNF16 expression did not increase the activity of the rDNA promoter in U2OS (Fig. 3B) and increased it only slightly in HeLa cells (Fig. 3A), suggesting that the ability of 3xEGFP-ZNF16 to localize to the nucleolus might be important for its ability to activate the rDNA promoter. Furthermore, this effect is specific to the rDNA promoter since a similar experiment using a luciferase reporter driven by a cytomegalovirus promoter (CMV) resulted in a decrease in luciferase activity after expression of 3xEGFP-ZNF16 in both cell lines (Fig. 3A-B).

**Figure 3.**
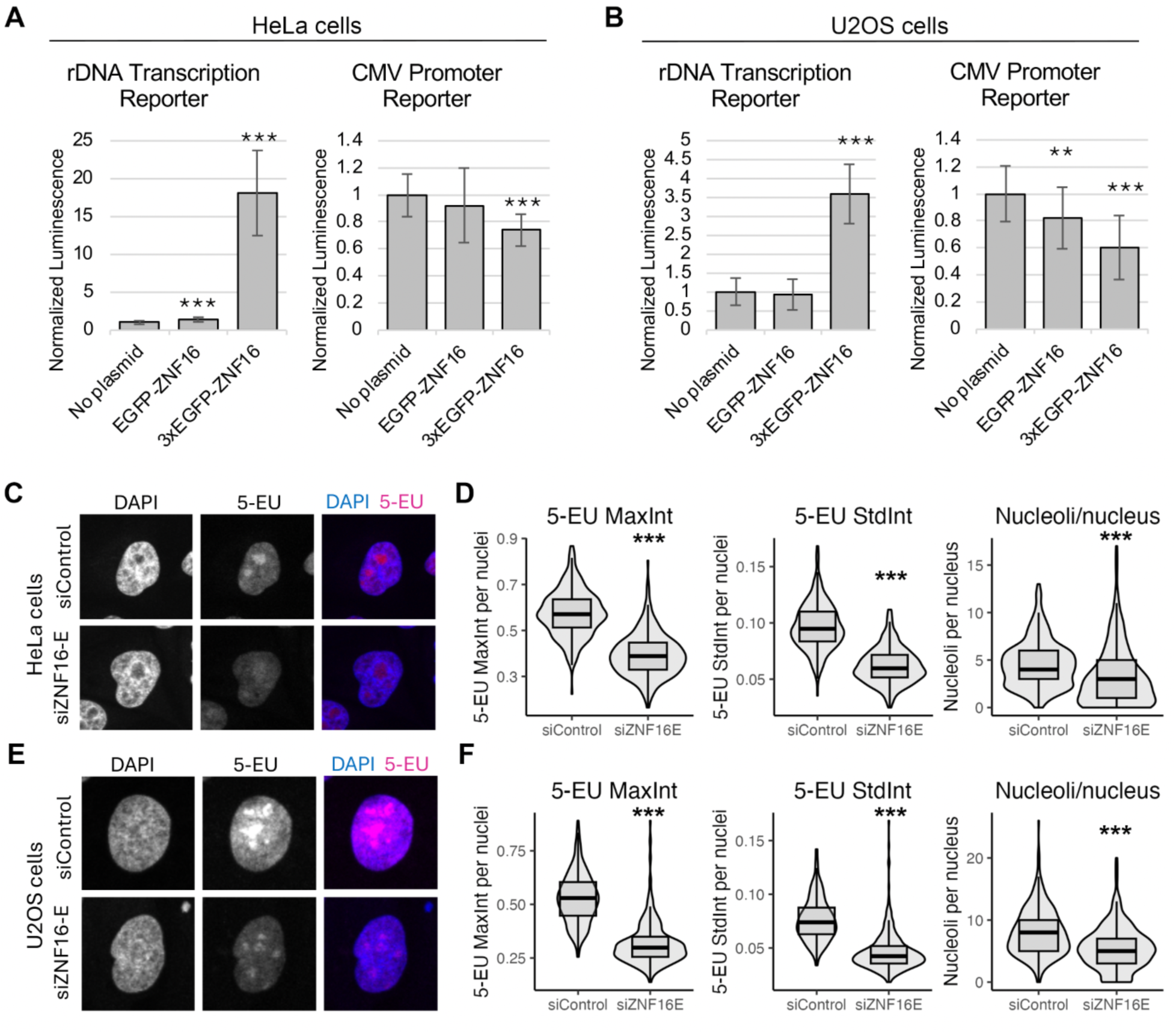
ZNF16 regulates rDNA transcription. (A-B) Luciferase assays after co-transfection of HeLa (A) or U2OS (B) cells with plasmids containing EGFP, EGFP-ZNF16 or 3xEGFP-ZNF16 in combination with either an rDNA transcription luciferase reporter or a CMV promoter luciferase reporter. Bar graphs represent the average of triplicate experiments and error bars are the standard deviation. Asterisks denote p-values as follows: *p<0.05, **p<0.01, ***p<0.001. (C-F) 5-EU incorporation assay for HeLa (C-D) or U2OS cells (E-F). (C, E) Representative images of HeLa (C) or U2OS cells (E) visualized with Alexa594 conjugated 5-EU and counterstained with DAPI. (D, F) Quantification of the number of nucleoli per nucleus, maximum intensity of 5-EU per nucleus (5-EU MaxInt) and standard deviation of the 5-EU intensity per nucleus (5-EU StdInt) for HeLa cells (D) or U2OS cells (F).

The role of endogenous ZNF16 in rDNA transcription was tested via a 5-EU incorporation assay. U2OS or HeLa cells were incubated with siControl or siZNF16-E siRNAs for 72h. Then, the cells were incubated with the nucleoside analog 5-EU, which is incorporated into nascent RNA transcripts and can be visualized by conjugation with Alexa-594 via click chemistry (Jao and Salic, 2008). Since ∼90% of the total RNA in human cells corresponds to rRNA (Palazzo and Lee, 2015), and we observed that the bulk of the 5-EU-containing RNA is localized in the nucleoli of siControl-transfected cells (Fig. 3C,E), quantification of 5-EU intensity was performed in whole nuclei. To quantify 5-EU intensity, confocal images were segmented on the DAPI channel using CellProfiler (Stirling et al., 2021) to detect individual nuclei and several measures of signal intensity (e.g., maximum intensity of 5-EU signal, standard deviation of the 5-EU signal) were obtained per nucleus. Given that the nucleolus is the region of the nucleus that shows the highest level of 5-EU incorporation, we reasoned that comparing the maximum intensity of 5-EU signal per nuclei reflects the levels of maximum rDNA transcription in the cells. Comparisons using this metric showed a significant decrease in maximum 5-EU intensity in cells transfected with siZNF16-E compared to siControl (Fig. 3D,F). Another way to quantify the extent of rDNA transcription is by looking at the difference between the high transcription regions (nucleoli) and the low transcription regions (nucleoplasm) by measuring the standard deviation of the 5-EU intensities in the nucleus. This analysis showed a significant decrease in the standard deviation of the intensity in siZNF16-E vs siControl-transfected cells. A lower standard deviation indicates a more homogenous 5-EU staining, as observed in siZNF16-E treated cells that have lower 5-EU staining in the nucleoli that is closer in value to the staining in the nucleoplasm (Fig. 3C,E). These metrics were chosen because they capture more accurately the changes in 5-EU incorporation in the nucleoli as compared to the measures that quantify 5-EU incorporation in the whole nucleus such as the mean intensity, which also showed a significant decrease after siZNF16-E transfection (data not shown).

In addition, segmentation of the nucleoli was attempted on the 5-EU channel using CellProfiler in order to compare the number, size and intensity of the nucleoli. However, due to the large difference in 5-EU intensity levels between siControl and siZNF16-E cells we were unable to find segmentation settings that worked well to identify all nucleoli in both treatments. Quantification of the number of nucleoli per nucleus from an attempt using settings that successfully identified all nucleoli in the siControl cells is shown in Figure 3D, F (right-most graphs). These results show a significant decrease in the number of nucleoli per nucleus in cells treated with siZNF16-E compared to siControl. However, this is likely due to the failure to identify many nucleoli in the siZNF16-E treated cells due to the low intensity of 5-EU in these cells. Although these results might not be an actual reflection of the number of nucleoli in these cells, they are consistent with a significant decrease in rDNA transcription that results in the inability to segment many nucleoli in siZNF16-E treated cells. Taken together, these results indicate a role for ZNF16 in promoting rDNA transcription.

### ZNF16 binds preferentially in the intergenic spacer region of the rDNA

Since our previous results indicated that ZNF16 regulates rDNA transcription and given that ZNF16 contains multiple zinc-finger motifs, which are commonly associated with DNA binding, we tested whether ZNF16 interacts with the rDNA region by chromatin immunoprecipitation (ChIP). Cells stably expressing 3xEGFP-ZNF16 were crosslinked and incubated with anti-EGFP antibody or control IgG. Binding to different regions of the rDNA was quantified by qPCR using primers targeted to different rDNA regions (Zhu et al., 2010)(Fig. 4A). All regions of the rDNA showed at least 5-fold binding enrichment in anti-GFP ChIPs compared to control IgG (Fig. 4B). Interestingly, primers H27, H36 and H42, which target the second half of the intergenic spacer region (IGS), have greater than 15-fold enrichment compared to IgG (Fig. 4B), indicating that ZNF16 binds preferentially to this region of the IGS. These results indicate that ZNF16 regulates rDNA transcription via binding to the IGS region.

**Figure 4.**
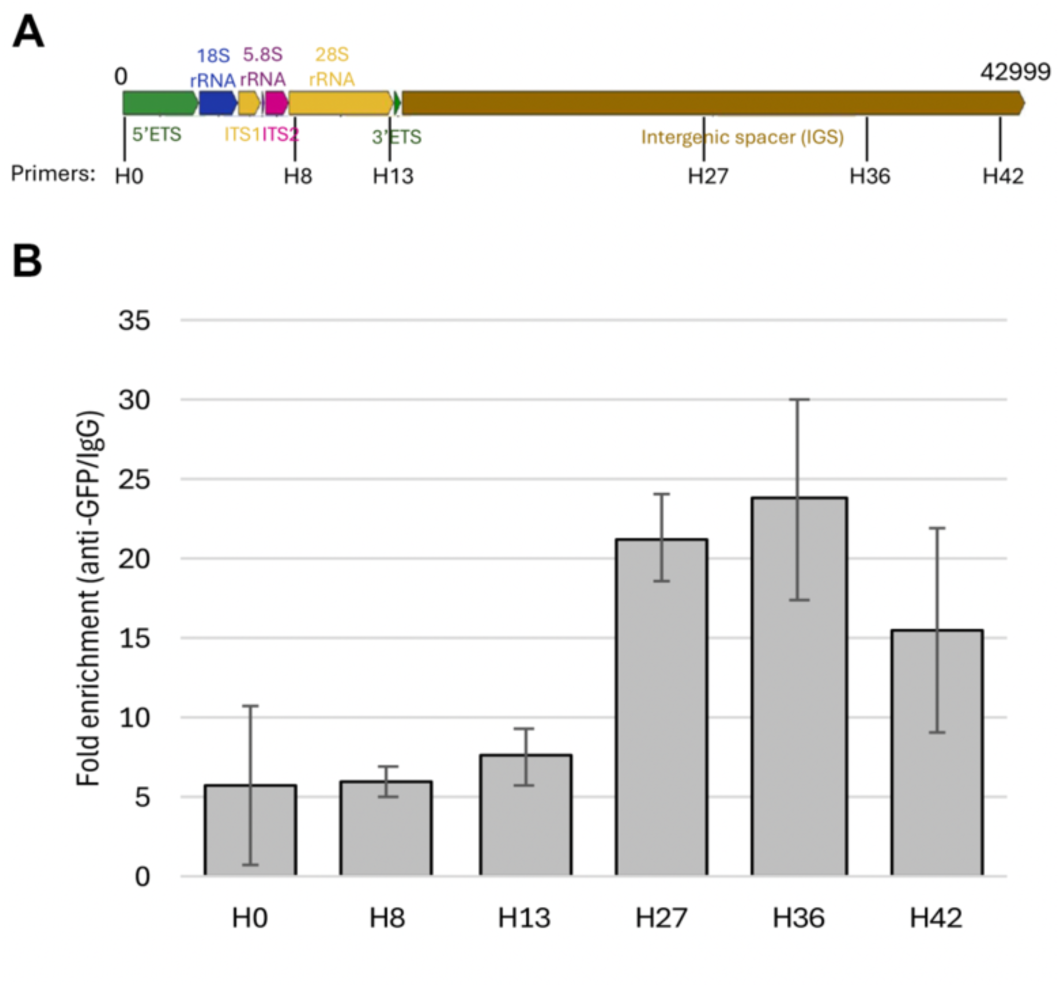
ZNF16 binds to the intergenic spacer (IGS) region of the rDNA. (A) Schematic of the rDNA unit showing the location of the qPCR primers (H0 to H42). (B) Quantification of fold enrichment for different regions in the rDNA unit via chromatin immunoprecipitation (ChIP) with anti-GFP and control IgG antibodies. Mean and standard deviation of three ChIPs.

### ZNF16 regulates expression of genes in a diversity of pathways and biological processes

To further explore the function of ZNF16, changes in gene expression after ZNF16 depletion were tested by RNA sequencing (RNA-seq). The siRNA siZNF16-E was selected for this experiment due to its consistent reduction of ZNF16 mRNA (Fig. 1B) and protein levels (Fig. 1C). Comparison between the non-targeting siControl and siZNF16-E transfected samples identified 2,833 differentially expressed genes out of a total 21,291 genes with measured expression (Fig. 5A).

**Figure 5.**
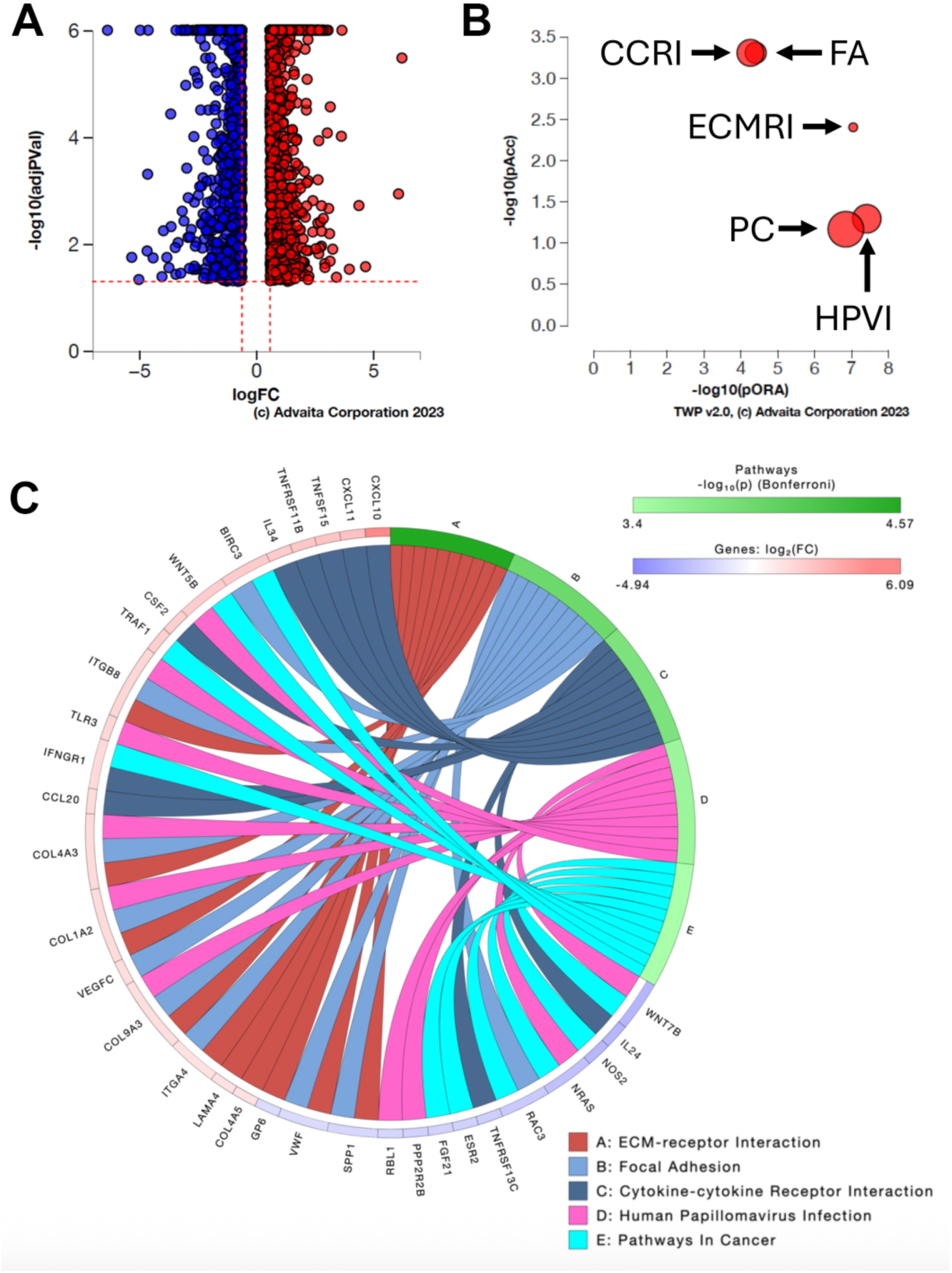
Top five pathways affected by ZNF16 depletion. U2OS cells were transfected with siControl or siZNF16-E siRNAs and gene expression was measured by RNA-seq. (A) Volcano plot showing the 2883 significantly differentially expressed genes represented by their expression change (logFC) vs the significance of the change (- log10(adjPVal)). Upregulated genes are shown in red and downregulated genes in blue. (B) Graph showing the top five pathways associated with ZNF16 depletion based on pathway overrepresentation (pORA) and total pathway accumulation (pAcc). Each pathway is represented by a single bubble and the size of the bubble is proportional to the size of the pathway it represents. Abbreviations: FA = Focal adhesion, CCRI = Cytokine-cytokine receptor interaction, ECM = extracellular matrix-receptor interaction, HPVI = Human papillomavirus infection, PC = pathways in cancer. (C) Chord diagram of the top 10 differentially expressed genes in the five pathways. Each pathway is represented by a different chord color as shown in the legend. Changes in gene expression (log2FC) are depicted in red-blue as shown in the legend.

Pathway analysis using iPathwayGuide identified focal adhesion (FA), cytokine-cytokine receptor interaction (CCRI), extracellular matrix-receptor interaction (ECMRI), human papillomavirus infection (HPVI), and pathways in cancer (PC) as the top five pathways affected by ZNF16 depletion (Fig. 5B and Table 1). Many of the top ten differentially expressed genes in each pathway (e.g., BIRC3, WNT5B, ITGB8, NRAS) are present in more than one pathway as can be seen in the chord diagram (Fig. 5C).

**Table 1.**
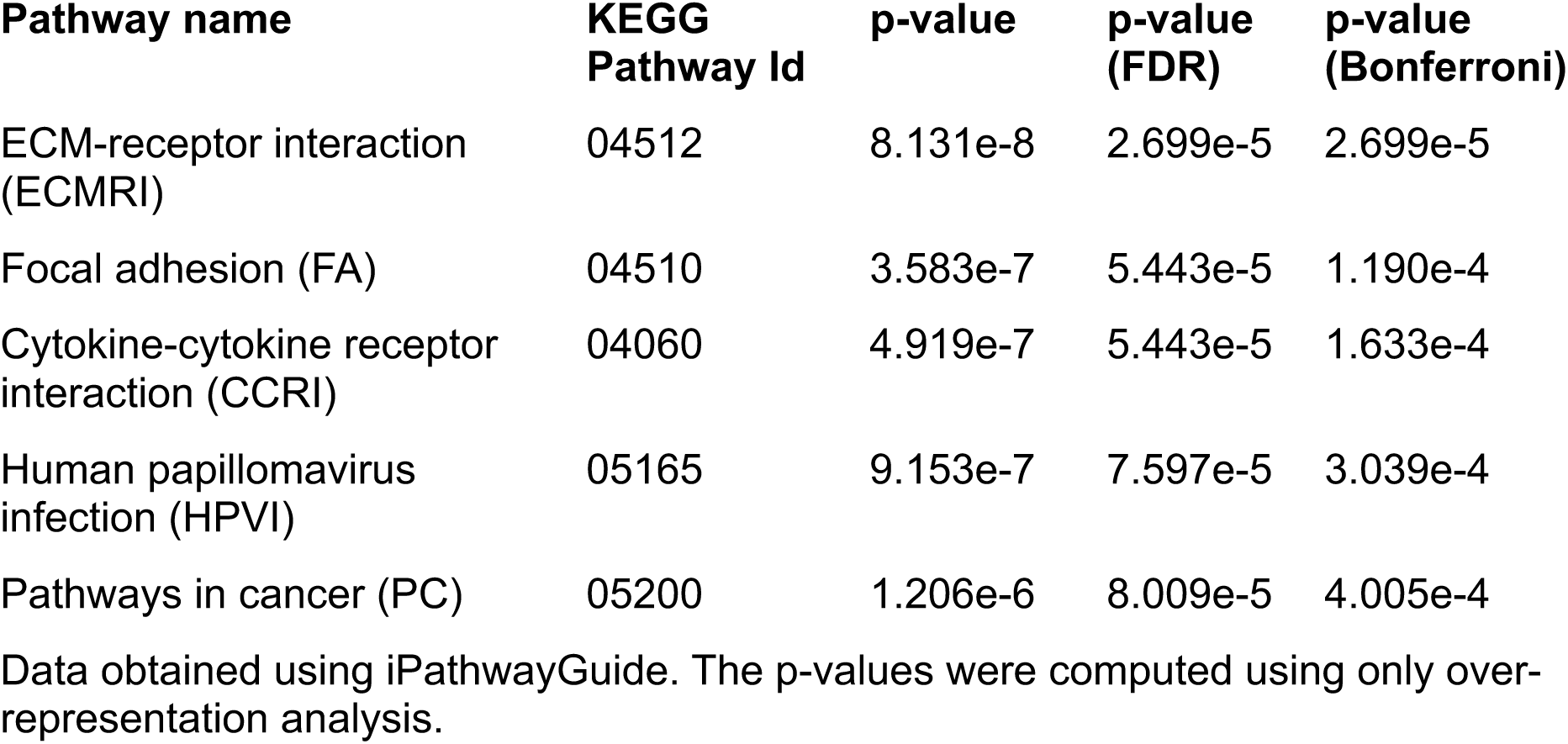
Top five pathways identified by pathway analysis of RNA-seq data after depletion of ZNF16.

Gene ontology (GO) analysis was performed using iPathwayGuide with high-specificity and smallest common denominator pruning to identify biological processes and molecular functions associated with ZNF16. The top five biological processes identified with these two methods are shown in Table 2, and the top five molecular functions identified are shown in Table 3. The top biological process and molecular function are both related to extracellular matrix biology, indicating a potential novel function for ZNF16 that remains to be explored.

**Table 2.**
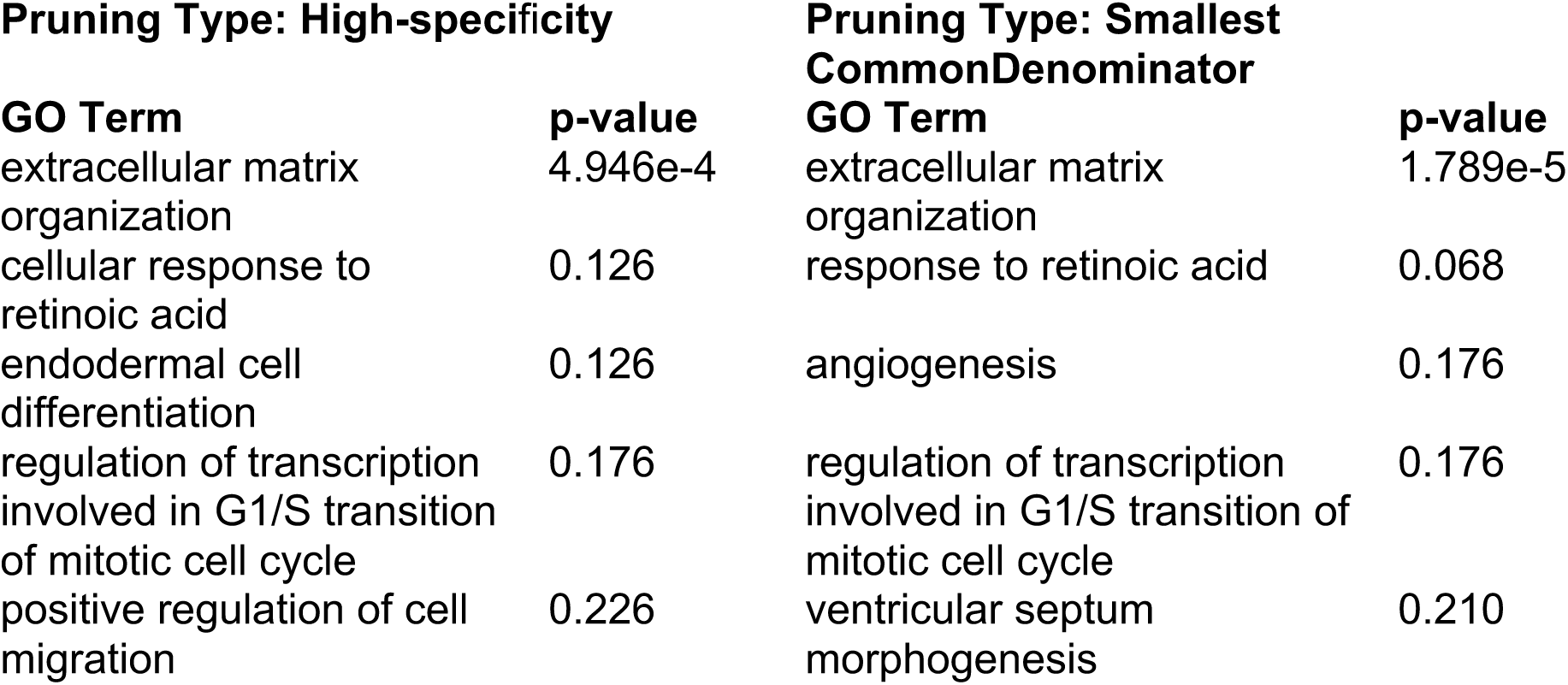
Top five biological processes identified by gene ontology (GO) analysis.

**Table 3.**
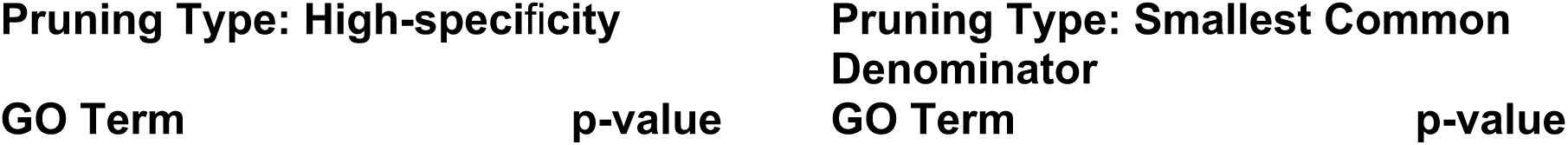
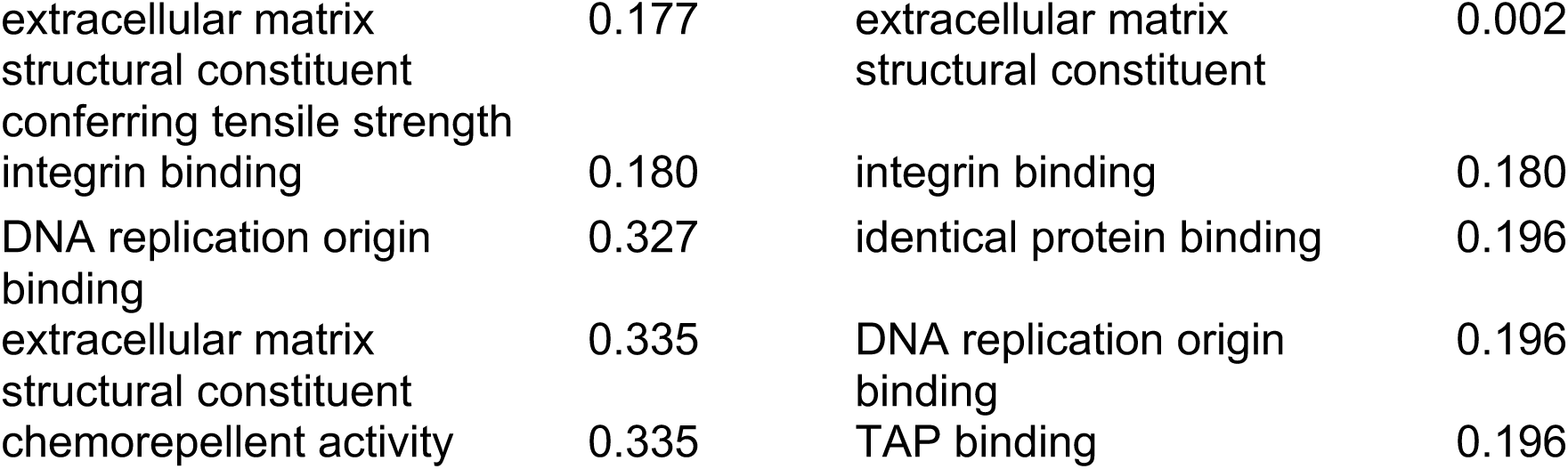
Top five molecular functions identified by gene ontology (GO) analysis.

**Pruning Type: High-specificity**
| GO Term | p-value |
| --- | --- |

| GO Term | p-value |
| --- | --- |

|  |  |  |  |
| --- | --- | --- | --- |
| extracellular matrix<br>structural constituent<br>conferring tensile strength | 0.177 | extracellular matrix<br>structural constituent | 0.002 |
| integrin binding | 0.180 | integrin binding | 0.180 |
| DNA replication origin<br>binding | 0.327 | identical protein binding | 0.196 |
| extracellular matrix<br>structural constituent | 0.335 | DNA replication origin<br>binding | 0.196 |
| chemorepellent activity | 0.335 | TAP binding | 0.196 |

### ZNF16 depletion affects expression of genes involved in cancer-related pathways

Given that ZNF16 expression and mutations have been associated with gallbladder carcinoma (Ahn et al., 2020) and tongue squamous cell carcinoma (Zhang et al., 2020), we next explored differentially expressed genes that are related to cancer processes. The top twenty differentially expressed genes in pathways in cancer (PC) include the well-known tumor suppressor NRAS (Hobbs et al., 2016), the inhibitor of apoptosis BIRC3/cIAP2 (Frazzi, 2021), and WNT7B, a component of the WNT/μ-catenin pathway (Arensman et al., 2014) (Fig. 6A). The results from the RNA-seq experiment for these three genes were confirmed by qPCR (Fig. 6B), showing statistically significant upregulation of BIRC3 and downregulation of NRAS and WNT7B in U2OS cells transfected with siZNF16-E. Another cancer-associated gene that was differentially expressed in the RNA-seq experiment but was not listed in the pathways in cancer list is EGFR. Upregulation of EGFR after ZNF16 depletion was similarly confirmed by qPCR (Fig. 6C).

**Figure 6.**
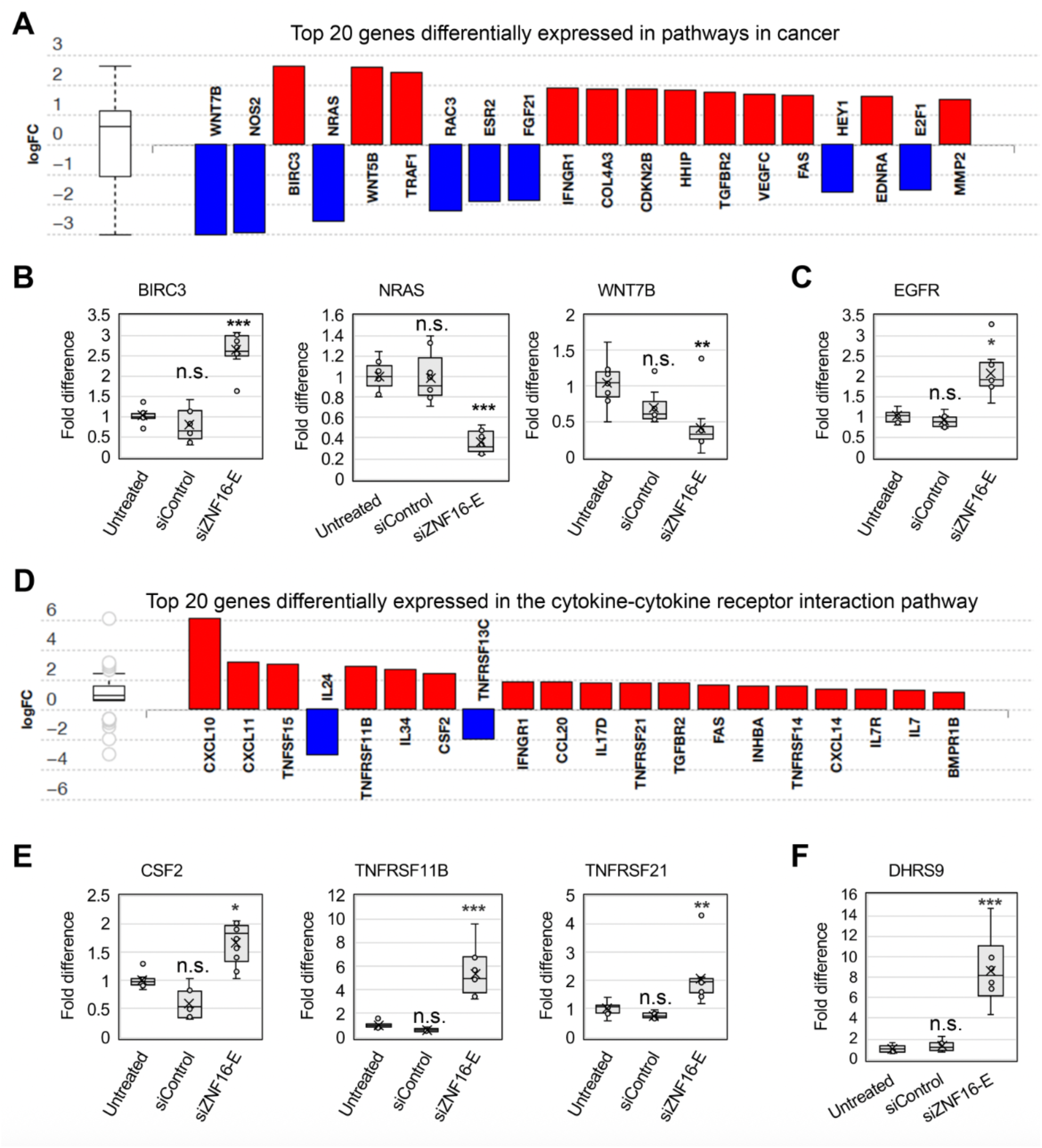
ZNF16 depletion affects expression of genes in cancer-associated pathways. (A) Top 20 genes differentially expressed in pathways in cancer (KEGG: 05200). Upregulated genes shown in red, downregulated genes in blue. (B) Independent confirmation of gene expression changes by qPCR for three genes in the pathway shown in (A). Box plots represent the results from three experiments. Fold difference normalized to the average of the untreated sample. (C) Fold difference in the expression of EGFR. Box plot represents the average of three experiments. Fold difference normalized to the average of the untreated sample. (D) Top 20 genes differentially expressed in the cytokine-cytokine receptor interaction (CCRI) pathway (KEGG: 04060). Upregulated genes shown in red, downregulated genes in blue. (E) Independent confirmation of gene expression changes by qPCR for three genes in the CCRI pathway shown in (D). Box plots represent the results from three experiments. Fold difference normalized to the average of the untreated sample. (F) Fold difference in the expression of DHRS9. Box plot represents the average of three experiments. Fold difference normalized to the average of the untreated sample. *p<0.05, **p<0.01, ***p<0.001

Next, we explored the differentially expressed genes included in the cytokine-cytokine receptor interaction (CCRI) pathway (Fig. 6D). Cytokines are central regulators of immunity and inflammation and play a key role in anti-tumor immunity (Kureshi and Dougan, 2025). The results from the RNA-seq experiment for three genes from the CCRI pathway were confirmed by qPCR (Fig. 6E), showing statistically significant upregulation of CSF2, TNFRSF11B, and TNFRSF21 in U2OS cells transfected with siZNF16-E. Another gene involved in immune regulation and associated with poor prognosis in tongue squamous cell carcinoma is DHRS9 (Shimomura et al., 2018). DHRS9 was significantly increased in the RNA-seq experiment but was not listed in the CCRI pathway. Upregulation of DHRS9 after ZNF16 depletion was similarly confirmed by qPCR (Fig. 6F).

### Overexpression of nucleolar ZNF16 reduces DNA damage by camptothecin

Given its association with cancer pathways, we next tested whether ZNF16 overexpression affected the cellular response to DNA-damaging agents commonly used as anti-cancer therapies. U2OS and HeLa cells transfected with the 3xEGFP-ZNF16 plasmid and incubated with 10 μM camptothecin for four hours had decreased levels of DNA damage, as measured by yH2AX staining (Fig. 7A-B). However, treatment with cisplatin and X-rays resulted in contradictory results in HeLa vs U2OS. HeLa cells showed increased DNA damage in the presence of cisplatin when transfected with 3xEGFP-ZNF16 while U2OS showed decreased DNA damage (Fig. 7C). Conversely, HeLa cells showed decreased DNA damage in response to X-rays when transfected with 3xEGFP-ZNF16 while U2OS showed increased damage (Fig. 7C). Thus, while ZNF16 overexpression changes DNA damage levels, the specific role of ZNF16 in the response to DNA damage agents depends on the type of agent, the localization of ZNF16 and the type of cell line being tested.

**Figure 7.**
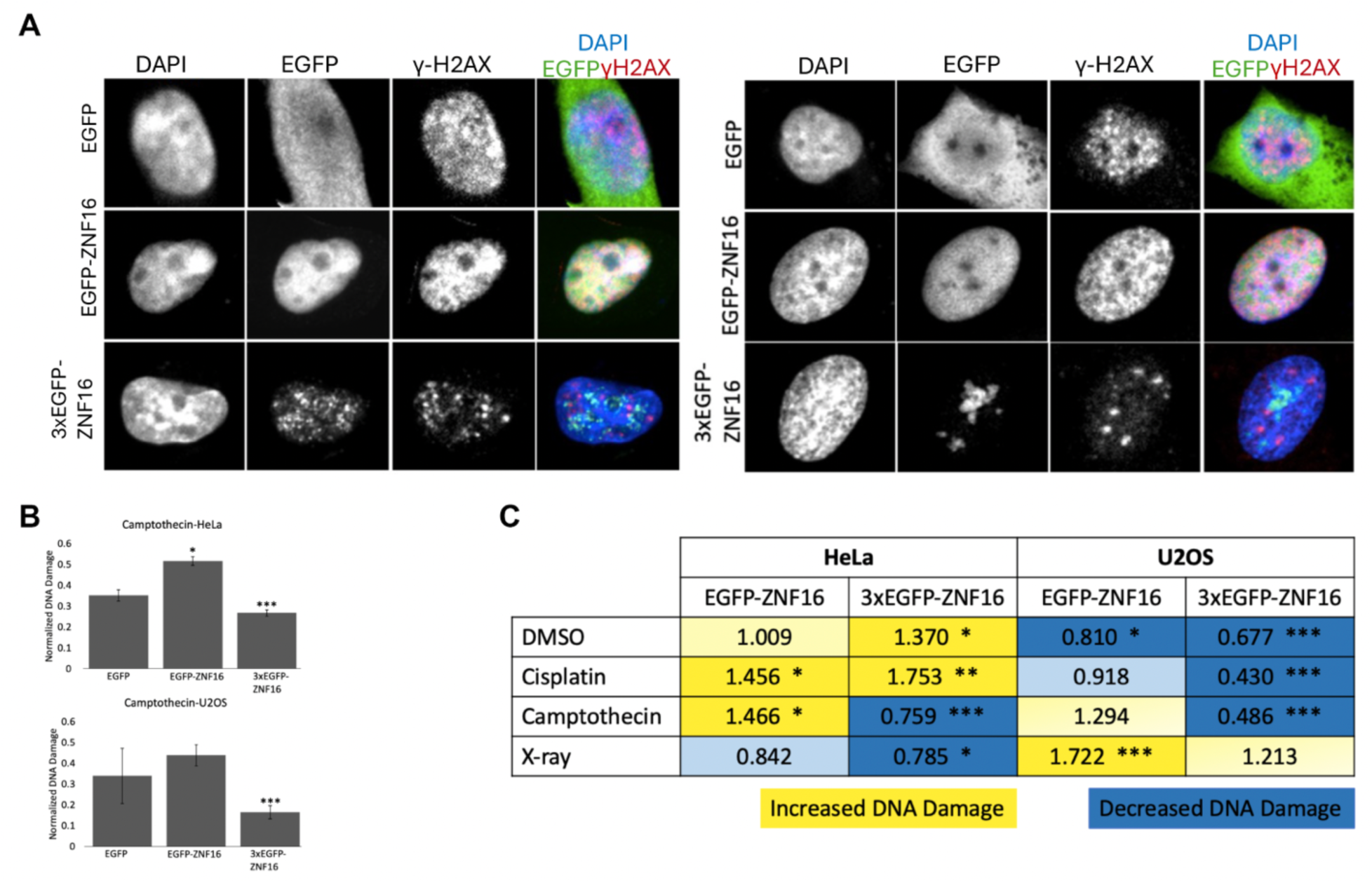
ZNF16 overexpression reduces DNA damage by camptothecin. (A) Representative micrographs of HeLa (left) and U2OS (right) cells transiently transfected with the plasmids containing EGFP, EGPF-ZNF16, or 3xEGFP-ZNF16, incubated with 10 μM camptothecin for 4 hours, and immunostained with the indicated antibodies. (B) Quantification of the yH2AX signal for the experiment in (A). (C) Summary of fold yH2AX intensity for different DNA damage treatments after expression of EGFP-ZNF16 and 3xEGFP-ZNF16. The fold yH2AX intensity is normalized to the intensity in the EGFP-transfected cells with the same treatment.

Together, our results expand our understanding of ZNF16 function by characterizing ZNF16 as a nucleolar protein that regulates rDNA expression via interaction with the intergenic spacer region of the rDNA unit and providing evidence for other potential roles for ZNF16 in regulating pathways involved in cancer, immune regulation, and extracellular matrix function.

## Discussion

ZNF16 was previously studied in the context of blood cell differentiation but it is a ubiquitously expressed protein (Lorenz et al., 2010; Peng et al., 2006). Our research has confirmed that ZNF16 is expressed in different cancer and non-cancer cell lines and identified a novel role for ZNF16 at the nucleolus as a regulator of rDNA expression. These results are consistent with evidence from co-IPs that showed interaction between ZNF16 and several nucleolar proteins including TCOF1, UBTF, NOL9, and PELP1 (Schmitges et al., 2016). Our results from the 3xEGFP-ZNF16 ChIP indicate that ZNF16 likely regulates rDNA transcription via binding to the IGS region. Importantly, the ribosomal DNA consists of multiple repeats of the rDNA, one after the other. Given that ZNF16 preferentially binds to the 3’ half of the IGS (Fig. 4B), this region is physically located upstream of the next rDNA sequence. However, the end of the IGS is still ∼4000 bp upstream of the 18S rRNA sequence, indicating that most likely ZNF16 regulates rDNA transcription via long-distance chromatin interactions rather than by directly binding to the rDNA promoter.

Several zinc finger proteins are reported to localize at the nucleolus, including LYAR (Izumikawa et al., 2019), PARP-1 (Meder et al., 2005), PHF12 (also known as PF1) (Graveline et al., 2017), ZNF274 (Yano et al., 2000), ZNF692 (Lafita-Navarro et al., 2023) and ZPR1 (Galcheva-Gargova et al., 1998). PARP-1, PHF12, ZPR1, and ZNF692 are involved in rRNA processing and/or ribosomal biogenesis (Galcheva-Gargova et al., 1998; Graveline et al., 2017; Izumikawa et al., 2019; Lafita-Navarro et al., 2023), while LYAR and ZNF274 are transcription regulators (Begnis et al., 2024; Izumikawa et al., 2019). These two rDNA transcriptional regulators have been shown to act in different ways: ZNF274 is a transcriptional repressor that sequesters gene clusters to transcriptionally inactive perinucleolar regions (Begnis et al., 2024), while LYAR directly binds near the rDNA promoter and promotes histone acetylation via recruitment of the BRD2-KAT7 complex (Izumikawa et al., 2019). Our results indicate that ZNF16 is another rDNA transcriptional regulator that likely works via interaction with the IGS. However, whether the interaction is direct or indirect remains to be determined.

In addition to its role at the nucleolus, RNA-seq analysis after depletion of ZNF16 indicates that changes in ZNF16 levels impact gene expression of pathways involved in immune regulation and cancer. ZNF16 depletion results in changes in expression of well-characterized tumor suppressors and oncogenes like NRAS and EGFR. Further exploration of this role of ZNF16 will shed light on its association with tumor progression (Ahn et al., 2020; Lee et al., 2021). Lastly, the GO term analysis also shows potential involvement of ZNF16 with extracellular membrane functions, a potential function that remains to be explored.

## Acknowledgements

Thanks to Dr. Sid Das and all members of the Das and Diaz-Martinez labs for support and productive discussions, R. Warrington for assistance with cloning, A. Levario for generation of stable cell line, B. Bell for the ImageJ macro, M. Cannata for assistance with the 5-EU incorporation assay, M. Blankenfeld and L. LeBlanc for assistance with qPCR. This work was supported by the M.J. Murdock Charitable Trust Grant # 202016503 to LADM. Sequencing at the UAMS Genomics Core facility and bioinformatics support were respectively provided through the Research Technology Core and Bioinformatics Research Support Core of the Arkansas INBRE program, by a grant from the National Institute of General Medical Sciences (NIGMS), P20 GM103429 from the National Institutes of Health. Undergraduate student research was partially supported by Gonzaga Science Research Program (GSRP) and the Cell Biology Education Consortium (CBEC) via NSF awards #18270660 and #2316122 to N. Reyna. The contents of this manuscript are solely the responsibility of the authors and do not necessarily represent the official views of NIH or NSF.

## Declaration of Interest Statement

The authors report there are no competing interests to declare.

## Author contributions according to CRediT taxonomy

Chelsea L. George: formal analysis, investigation, validation, visualization & writing (original draft, reviewing & editing).

Laura A. Espinoza Quevedo: formal analysis, investigation, validation, visualization & writing (original draft, reviewing & editing).

Jason Paratore: formal analysis, investigation & validation.

Matthew J. Alcaraz: formal analysis, investigation, validation & writing (review & editing).

Arlene P. Levario: investigation, resources.

Yasir Rahmatallah: formal analysis, methodology, resources & writing (review & editing).

Galina V. Glazko: formal analysis, investigation, methodology, resources.

Nathan S. Reyna: formal analysis, funding acquisition, resources.

Laura A. Diaz-Martinez: conceptualization, data curation, formal analysis, funding acquisition, investigation, project administration, supervision, validation, visualization & writing (original draft, reviewing & editing).

## Notes

### Competing Interest Statement

The authors have declared no competing interest.

## References

Ahn, S. W., Ahn, A. R., Ha, S. H., Hussein, U. K., Do Yang, J., Kim, K. M., Park, H. S., Park, S. H., Yu, H. C. and Jang, K. Y. (2020). Expression of FAM83H and ZNF16 are associated with shorter survival of patients with gallbladder carcinoma. Diagn Pathol 15, 1–14.

Andreini, C., Banci, L., Bertini, I. and Rosato, A. (2006). Counting the zinc-proteins encoded in the human genome. J Proteome Res 5, 196–201.

Arensman, M. D., Kovochich, A. N., Kulikauskas, R. M., Lay, A. R., Yang, P. T., Li, X., Donahue, T., Major, M. B., Moon, R. T., Chien, A. J., et al. (2014). WNT7B mediates autocrine Wnt/β-catenin signaling and anchorage-independent growth in pancreatic adenocarcinoma. Oncogene 33, 899–908.

Begnis, M., Duc, J., Offner, S., Grun, D., Sheppard, S., Rosspopoff, O. and Trono, D. (2024). Clusters of lineage-specific genes are anchored by ZNF274 in repressive perinucleolar compartments. Sci. Adv 10, eado1662.

Bolger, A. M., Lohse, M. and Usadel, B. (2014). Trimmomatic: A flexible trimmer for Illumina sequence data. Bioinformatics 30, 2114–2120.

Chen, J., Li, X.-B., Su, R., Song, L., Wang, F. and Zhang, J.-W. (2014). ZNF16 (HZF1) promotes erythropoiesis and megakaryocytopoiesis via regulation of the c-KIT gene. Biochemical Journal 458, 171–183.

Deng, M.-J., Li, X.-B., Peng, H. and Zhang, J.-W. (2010). Identification of the Trans-Activation Domain and the Nuclear Location Signals of Human Zinc Finger Protein HZF1 (ZNF16). Mol Biotechnol 44, 83–89.

Frazzi, R. (2021). BIRC3 and BIRC5: multi-faceted inhibitors in cancer. Cell Biosci 11, 8.

Galcheva-Gargova, Z., Gangwani, L., Konstantinov, K. N., Mikrut, M., Theroux, S. J., Enoch, T. and Davis, R. J. (1998). The cytoplasmic zinc finger protein ZPR1 accumulates in the nucleolus of proliferating cells. Mol Biol Cell 9, 2963–71.

Gamsjaeger, R., Liew, C. K., Loughlin, F. E., Crossley, M. and Mackay, J. P. (2007). Sticky fingers: zinc-fingers as protein-recognition motifs. Trends Biochem Sci 32,.

Ghoshal, K., Majumder, S., Datta, J., Motiwala, T., Bai, S., Sharma, S. M., Frankel, W. and Jacob, S. T. (2004). Role of Human Ribosomal RNA (rRNA) Promoter Methylation and of Methyl-CpG-binding Protein MBD2 in the Suppression of rRNA Gene Expression. Journal of Biological Chemistry 279, 6783–6793.

González-Arzola, K. (2024). The nucleolus: Coordinating stress response and genomic stability. Biochim Biophys Acta Gene Regul Mech 1867, 195029.

Graveline, R., Marcinkiewicz, K., Choi, S., Paquet, M., Wurst, W., Floss, T. and David, G. (2017). The Chromatin-Associated Phf12 Protein Maintains Nucleolar Integrity and Prevents Premature Cellular Senescence. Mol Cell Biol 37, e00522–16.

Hobbs, G. A., Der, C. J. and Rossman, K. L. (2016). RAS isoforms and mutations in cancer at a glance. J Cell Sci 129, 1287–1292.

Hua, L., Yan, D., Wan, C. and Hu, B. (2022). Nucleolus and Nucleolar Stress: From Cell Fate Decision to Disease Development. Cells 11, 3017.

Izumikawa, K., Ishikawa, H., Yoshikawa, H., Fujiyama, S., Watanabe, A., Aburatani, H., Tachikawa, H., Hayano, T., Miura, Y., Isobe, T., et al. (2019). LYAR potentiates rRNA synthesis by recruiting BRD2/4 and the MYST-type acetyltransferase KAT7 to rDNA. Nucleic Acids Res 47, 10357–10372.

Jao, C. Y. and Salic, A. (2008). Exploring RNA transcription and turnover in vivo by using click chemistry. Proc Natl Acad Sci U S A 105, 15779–15784.

Jarboui, M. A., Wynne, K., Elia, G., Hall, W. W. and Gautier, V. W. (2011). Proteomic profiling of the human T-cell nucleolus. Mol Immunol 46, 441–452.

Klug, A. and Rhodes, D. (1987). “Zinc fingers”: a novel protein motif for nucleic acid recognition. Trends Biochem Sci 12, 464–469.

Kureshi, C. T. and Dougan, S. K. (2025). Cytokines in cancer. Cancer Cell 43, 15–35.

Lafita-Navarro, M. C., Hao, Y. H., Jiang, C., Jang, S., Chang, T. C., Brown, I. N., Venkateswaran, N., Maurais, E., Stachera, W., Zhang, Y., et al. (2023). ZNF692 organizes a hub specialized in 40S ribosomal subunit maturation enhancing translation in rapidly proliferating cells. Cell Rep 42, 113280.

Laity, J. H., Lee, B. M. and Wright, P. E. (2001). Zinc finger proteins: new insights into structural and functional diversity. Curr Opin Struct Biol 11, 39–46.

Lee, M., Yang, J. Do, Ahn, S. W. and Yu, H. C. (2021). A study for the expression of FAM83H, ZNF16, and FAM83H-related proteins in gallbladder cancer. Ann Hepatobiliary Pancreat Surg 25 Suppl 1, S337.

Li, X.-B., Chen, J., Deng, M.-J., Wang, F., Du, Z.-W. and Zhang, J.-W. (2011). Zinc finger protein HZF1 promotes K562 cell proliferation by interacting with and inhibiting INCA1. Mol Med Rep 4, 1131–1137.

Liao, Y., Smyth, G. K. and Shi, W. (2014). FeatureCounts: An efficient general purpose program for assigning sequence reads to genomic features. Bioinformatics 30, 923– 930.

Lorenz, P., Dietmann, S., Wilhelm, T., Koczan, D., Autran, S., Gad, S., Wen, G., Ding, G., Li, Y., Rousseau-Merck, M.-F., et al. (2010). The ancient mammalian KRAB zinc finger gene cluster on human chromosome 8q24.3 illustrates principles of C2H2 zinc finger evolution associated with unique expression profiles in human tissues. BMC Genomics 11, 206.

Love, M. I., Huber, W. and Anders, S. (2014). Moderated estimation of fold change and dispersion for RNA-seq data with DESeq2. Genome Biol 15, 550.

Meder, V. S., Boeglin, M., de Murcia, G. and Schreiber, V. (2005). PARP-1 and PARP-2 interact with nucleophosmin/B23 and accumulate in transcriptionally active nucleoli. J Cell Sci 118, 211–222.

Núñez Villacís, L., Wong, M. S., Ferguson, L. L., Hein, N., George, A. J. and Hannan, K. M. (2018). New Roles for the Nucleolus in Health and Disease. BioEssays 40, e1700233.

Palazzo, A. F. and Lee, E. S. (2015). Non-coding RNA: What is functional and what is junk? Front Genet 6, 2.

Peng, H., Du, Z.-W. and Zhang, J.-W. (2006). Identification and characterization of a novel zinc finger protein (HZF1) gene and its function in erythroid and megakaryocytic differentiation of K562 cells. Leukemia 20, 1109–1116.

Potapova, T. A. and Gerton, J. L. (2019). Ribosomal DNA and the nucleolus in the context of genome organization. Chromosome Research 27, 109–127.

Schmidt-Zachmann, M. S., Hügle-Dörr, B. and Franke, W. W. (1987). A constitutive nucleolar protein identified as a member of the nucleoplasmin family. EMBO J 6, 1881–1890.

Schmitges, F. W., Radovani, E., Najafabadi, H. S., Barazandeh, M., Campitelli, L. F., Yin, Y., Jolma, A., Zhong, G., Guo, H., Kanagalingam, T., et al. (2016). Multiparameter functional diversity of human C2H2 zinc finger proteins. Genome Res 26, 1742–1752.

Shimomura, H., Sasahira, T., Nakashima, C., Shimomura-Kurihara, M. and Kirita, T. (2018). Downregulation of DHRS9 is associated with poor prognosis in oral squamous cell carcinoma. Pathology 50, 642–647.

Stirling, D. R., Swain-Bowden, M. J., Lucas, A. M., Carpenter, A. E., Cimini, B. A. and Goodman, A. (2021). CellProfiler 4: improvements in speed, utility and usability. BMC Bioinformatics 22, 1–11.

Thiesen, H. J. (1990). Multiple genes encoding zinc finger domains are expressed in human T cells. New Biologist 2, 363–374.

Trapnell, C., Roberts, A., Goff, L., Pertea, G., Kim, D., Kelley, D. R., Pimentel, H., Salzberg, S. L., Rinn, J. L. and Pachter, L. (2012). Differential gene and transcript expression analysis of RNA-seq experiments with TopHat and Cufflinks. Nat Protoc 7, 562–578.

Trinkle-Mulcahy, L. (2018). Nucleolus: The Consummate Nuclear Body. Nuclear Architecture and Dynamics 257–282.

Urrutia, R. (2003). KRAB-containing zinc-finger repressor proteins. Genome Biol 4, 231.

Vilas, C. K., Emery, L. E., Denchi, E. L. and Miller, K. M. (2018). Caught with One’s Zinc Fingers in the Genome Integrity Cookie Jar. Trends in Genetics 34, 313.

Yano, K., Ueki, N., Oda, T., Seki, N., Masuho, Y. and Muramatsu, M. (2000). Identification and Characterization of Human ZNF274 cDNA, which Encodes a Novel Kruppel-type Zinc-Finger Protein Having Nucleolar Targeting Ability. Genomics 65, 75–80.

Zhang, P., Branson, O. E., Freitas, M. A. and Parthun, M. R. (2016). Identification of replication-dependent and replication-independent linker histone complexes: Tpr specifically promotes replication-dependent linker histone stability. BMC Biochem 17, 18.

Zhang, H., Song, Y., Du, Z., Li, X., Zhang, J., Chen, S., Chen, F., Li, T. and Zhan, Q. (2020). Exome sequencing identifies new somatic alterations and mutation patterns of tongue squamous cell carcinoma in a Chinese population. Journal of Pathology 251, 353–364.

Zhu, Z., Wang, Y., Li, X., Wang, Y., Xu, L., Wang, X., Sun, T., Dong, X., Chen, L., Mao, H., et al. (2010). PHF8 is a histone H3K9me2 demethylase regulating rRNA synthesis. Cell Res 20, 794–801.

