## Supplemental Materials for "ZNF16 is a nucleolar-associated protein that regulates expression of the rDNA and cancer-associated genes"

iPathwayGuide

| | |
|---|---|
| **Title:** | siControl vs siZNF-16E |
| **Description:** | File: Fixed for them DESeq2_results_siControl_siE.txt |
| **Organism:** | Homo sapiens (9606) |
| **Contrast** | siZNF16-E vs. siControl - mRNA (RNA-seq) |
| **Creation time:** | 10-22-2021 01:35 PM |

### 1. Introduction

In this experiment, **2,883** differentially expressed (DE) genes were identified out of a total of **21,291** genes with measured expression. These were identified using thresholds defined by the user. In this experiment, the user chose a threshold of **0.05** for statistical significance (p-value) and a log fold change of expression with absolute value of at least **0.6**. These data were analyzed in the context of pathways obtained from the Kyoto Encyclopedia of Genes and Genomes (KEGG) database (Release 96.0+/11-21, Nov 20) (Kanehisa *et al*., 2000; Kanehisa *et al*., 2002), gene ontologies from the Gene Ontology

Consortium database (2020-Oct14) (Ashburner *et al*., 2000; Gene Ontology Consortium, 2001), miRNAs from the miRBase (MIRBASE Version:Version22.1,10/18) and TARGETSCAN (Targetscan version: Mouse:7.2, Human:7.2) databases (Agarwal *et al*., 2015; Nam *et al*., 2014; Griffiths-Jones et al., 2008; Kozomara and Griffiths-Jones, 2014; Friedman *et al*., 2009; Grimson *et al*., 2007), network of regulatory relations from BioGRID: Biological General Repository for Interaction Datasets v4.0.189. Aug. 25th, 2020 (Szklarczyk *et al*., 2017), chemicals/drugs/toxicants from the Comparative Toxicogenomics Database July 2020 (Davis *et al*., 2019), and diseases from the KEGG database (Release 96.0+/11-21, Nov 20) (Kanehisa *et al*., 2000; Kanehisa *et al*., 2002). In summary, **88** pathways were found to be significantly impacted. In addition, **1,615** Gene Ontology (GO) terms, **2** miRNAs , **259** gene upstream regulators, **255** chemical upstream regulators and **51** diseases were found to be significantly enriched before the correction for multiple comparisons.

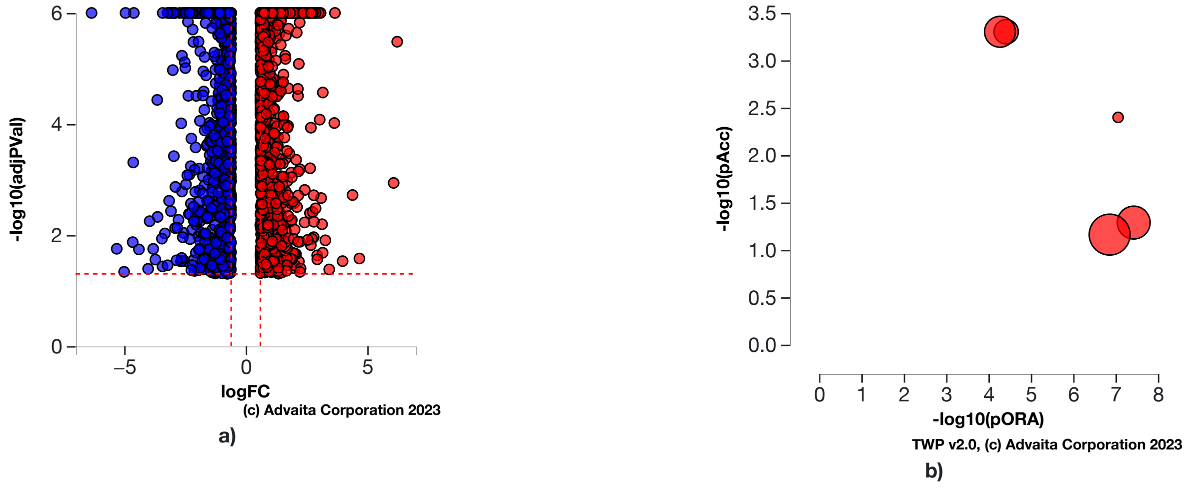


**Fig. 1.1: a) Volcano plot:** *All **2883** significantly differentially expressed (DE) genes are represented in terms of their measured expression change (x-axis) and the significance of the change (y-axis). The significance is represented in terms of the negative log (base 10) of the p-value, so that more significant genes are plotted higher on the y-axis. The dotted lines represent the thresholds used to select the DE genes: **0.6** for expression change and **0.05** for significance. The up-regulated genes (positive log fold change) are shown in red, while the down-regulated genes are blue.* **b) Pathways perturbation vs over-representation:** *The top 5 pathways are plotted in terms of the two types of evidence computed by iPathwayGuide: over-representation on the x-axis (pORA) and the total pathway accumulation on the y-axis (pAcc). Each pathway is represented by a single dot, with significant pathways shown in red, non-significant in black, and the size of each dot is proportional to the size of the pathway it represents. Both p-values are shown in terms of their negative log (base 10) values.*

### 2. Pathway Analysis

#### 2.1. Methods

iPathwayGuide scores pathways using the Impact Analysis method (Draghici *et al*., 2007; Tarca *et al*., 2009, Khatri *et al*., 2007). Impact analysis uses two types of evidence: i) the over-representation of differentially expressed (DE) genes in a given pathway and ii) the perturbation of that pathway computed by propagating the measured expression changes across the pathway topology. These aspects are captured by two independent probability values, pORA and pAcc, that are then combined in a unique pathway-specific p-value. The underlying pathway topologies, comprised of genes and their directional interactions, are obtained from the KEGG database (Kanehisa *et al*., 2000; Kanehisa *et al*., 2010; Kanehisa *et al*., 2012; Kanehisa *et al*., 2014).

The first probability, pORA, expresses the probability of observing the number of DE genes in a given pathway that is greater than or equal to the number observed, by random chance (Draghici *et al*., 2003; Draghici 2011). Let us consider there are *N* genes measured in the experiment, with *M* of these on the given pathway. Based on the user-defined a priori selection of DE genes, *K* out of *M* genes were found to be differentially expressed. The probability of observing exactly *x* differentially expressed genes on the given pathway is computed based on the hypergeometric distribution:

$$(1) \qquad P(X=x|N,M,K) = \frac{\binom{M}{x}\binom{N-M}{K-x}}{\binom{N}{K}}$$

Because the hypergeometric distribution is discrete, the probability of observing fewer than *x* genes on the given pathway just by chance can be calculated by summing the probabilities of randomly observing *0*, *1*, *2*, ..., up to *x-1* DE genes on the pathway:

$$(2) \qquad p_u(x-1) = P(X=1)+P(X=2)+\ldots+P(X=x-1) = \sum_{i=0}^{x-1} \frac{\binom{M}{i}\binom{N-M}{K-i}}{\binom{N}{K}}$$

iPathwayGuide calculates the probability of randomly observing a number of DE genes on the given pathway that is greater than or equal to the number of DE genes obtained from data, by computing the over-representation p-value: $pORA = p_o(x) = 1 - p_u(x-1)$:

$$(3) \qquad p_o(x) = 1 - \sum_{i=0}^{x-1} \frac{\binom{M}{i}\binom{N-M}{K-i}}{\binom{N}{K}}$$

The second probability, pAcc, is calculated based on the amount of total accumulation measured in each pathway. A perturbation factor is computed for each gene on the pathway using:

$$(4) \qquad PF(g) = \alpha(g) \cdot \triangle E(g) + \sum_{u \epsilon US_g} \beta_{ug} \frac{PF(u)}{N_{ds}(u)}$$

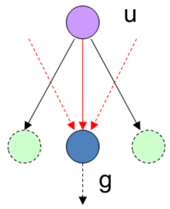


In Equation 4, *PF(g)* is the perturbation factor for gene *g*, the term *ΔE(g)* represents the signed normalized measured expression change of gene *g*, and *α(g)* is a priori weight based on the type of the gene. The last term is the sum of the perturbation factors of all genes *u*, directly upstream of the target gene *g*, normalized by the number of downstream genes of each such gene $N_{ds}(u)$. The value of $\beta_{ug}$ quantifies the strength of the interaction between genes *g* and *u*. The sign of *β* represents the type of interaction: positive for activation-like signals, and negative for inhibition-like signals. Subsequently, iPathwayGuide calculates the accumulation at the level of each gene, *Acc(g)*, as the difference between the perturbation factor *PF(g)* and the observed log fold-change:

$$(5) \qquad Acc(g_i) = PF(g_i) - \triangle E(g_i)$$

All perturbation accumulations are computed at the same time by solving the system of linear equations resulting from combining Equation 4 for all genes on a given pathway. Once all gene perturbation accumulations are computed, iPathwayGuide computes the total accumulation of the pathway as the sum of all absolute accumulations of the genes in a given pathway. The significance of obtaining a total accumulation (pAcc) at least as large as observed, just by chance, is assessed through bootstrap analysis.

The two types of evidence, pORA and pAcc, are combined into an overall pathway score by calculating a p-value using Fisher's method. This p-value is then corrected for multiple comparisons using false dicovery rate (FDR) and Bonferroni corrections. Bonferroni is simpler and more conservative of the two (Bonferroni, 1935; Bonferroni, 1936). It reduces the false discovery rate by imposing a stringent threshold on each comparison adjusted for the total number of comparisons. The FDR correction has more power, but only controls the family-wise false positives rate (Benjamini and Hochberg, 1995; Benjamini and Yekutieli, 2001).

### 2.2. Results

**Table 2.2.1: Top pathways and their associated p-values**

| Pathway name | Pathway Id | p-value | p-value (FDR) | p-value (Bonferroni) |
|---|---|---|---|---|
| ECM-receptor interaction | 04512 | **8.131e-8** | **2.699e-5** | **2.699e-5** |
| Focal adhesion | 04510 | **3.583e-7** | **5.443e-5** | **1.190e-4** |
| Cytokine-cytokine receptor interaction | 04060 | **4.919e-7** | **5.443e-5** | **1.633e-4** |
| Human papillomavirus infection | 05165 | **9.153e-7** | **7.597e-5** | **3.039e-4** |
| Pathways in cancer | 05200 | **1.206e-6** | **8.009e-5** | **4.005e-4** |

* the p-value corresponding to the pathway was computed using only over-representation analysis.

#### ECM-receptor interaction (KEGG: 04512)

The extracellular matrix (ECM) consists of a complex mixture of structural and functional macromolecules and serves an important role in tissue and organ morphogenesis and in the maintenance of cell and tissue structure and function. Specific interactions between cells and the ECM are mediated by transmembrane molecules, mainly integrins and perhaps also proteoglycans, CD36, or other cell-surface-associated components. These interactions lead to a direct or indirect control of cellular activities such as adhesion, migration, differentiation, proliferation, and apoptosis. In addition, integrins function as mechanoreceptors and provide a force-transmitting physical link between the ECM and the cytoskeleton. Integrins are a family of glycosylated, heterodimeric transmembrane adhesion receptors that consist of noncovalently bound alpha- and beta-subunits.

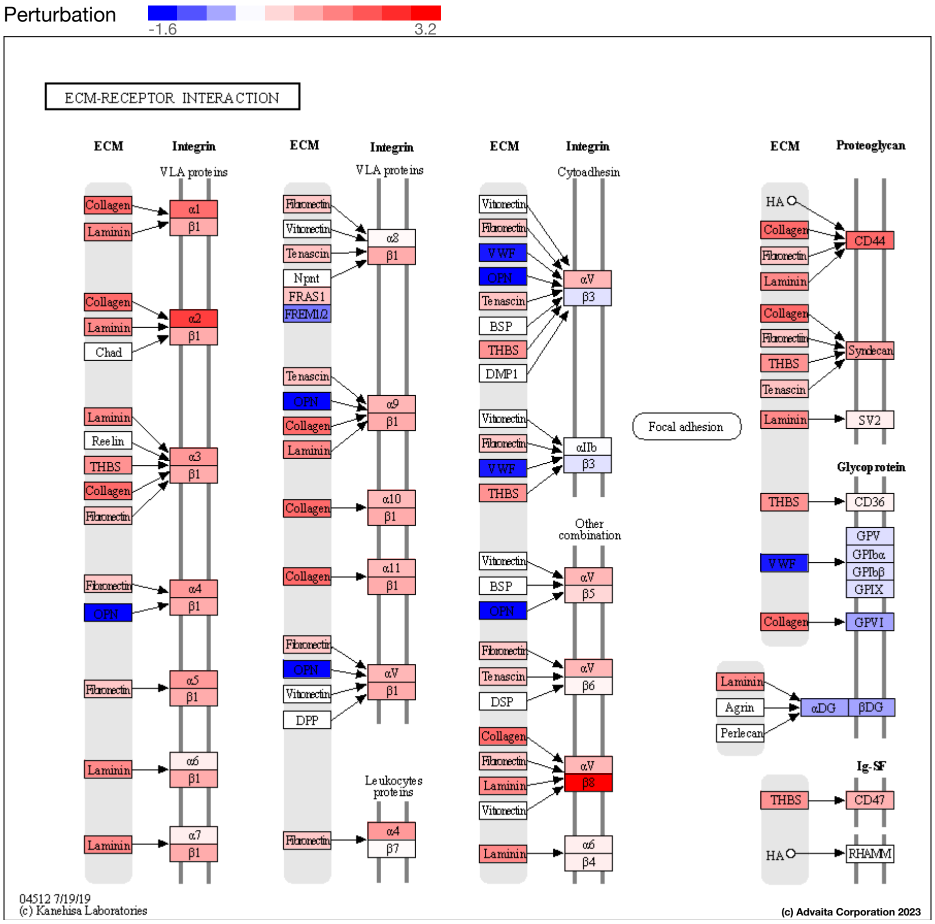


**Fig. 2.2.1: ECM-receptor interaction (KEGG: 04512):** *The pathway diagram is overlayed with the computed perturbation of each gene. The perturbation accounts both for the gene's measured fold change and for the accumulated perturbation propagated from any upstream genes (accumulation). The highest negative perturbation is shown in dark blue, while the highest positive perturbation in dark red. The legend describes the values on the gradient. Note: For legibility, one gene may be represented in multiple places in the diagram and one box may represent multiple genes in the same gene family. A gene is highlighted in all locations it occurs in the diagram. For each gene family, the color corresponding to the gene with the highest absolute perturbation is displayed.*

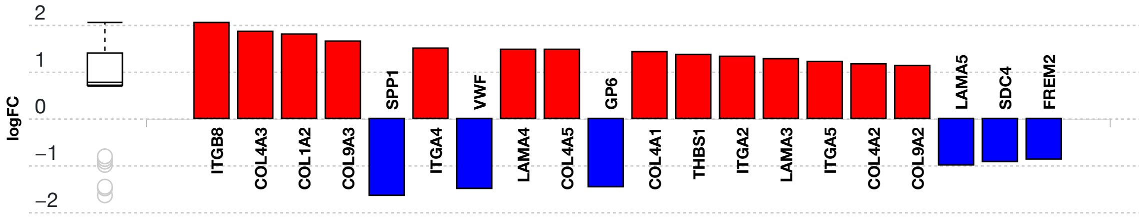


(c) Advaita Corporation 2023

**Fig. 2.2.2: Gene measured expression bar plot:** *All the differentially expressed genes in ECM-receptor interaction (KEGG: 04512) are ranked based on their absolute value of log fold change. The plot is limited to the top 20 genes out of a total of 31 differentially expressed genes. Upregulated genes are shown in red, downregulated genes are shown in blue. The box and whisker plot on the left summarizes the distribution of all the differentially expressed genes in this pathway. The box represents the 1st quartile, the median and the 3rd quartile, while the outliers are represented by circles.*

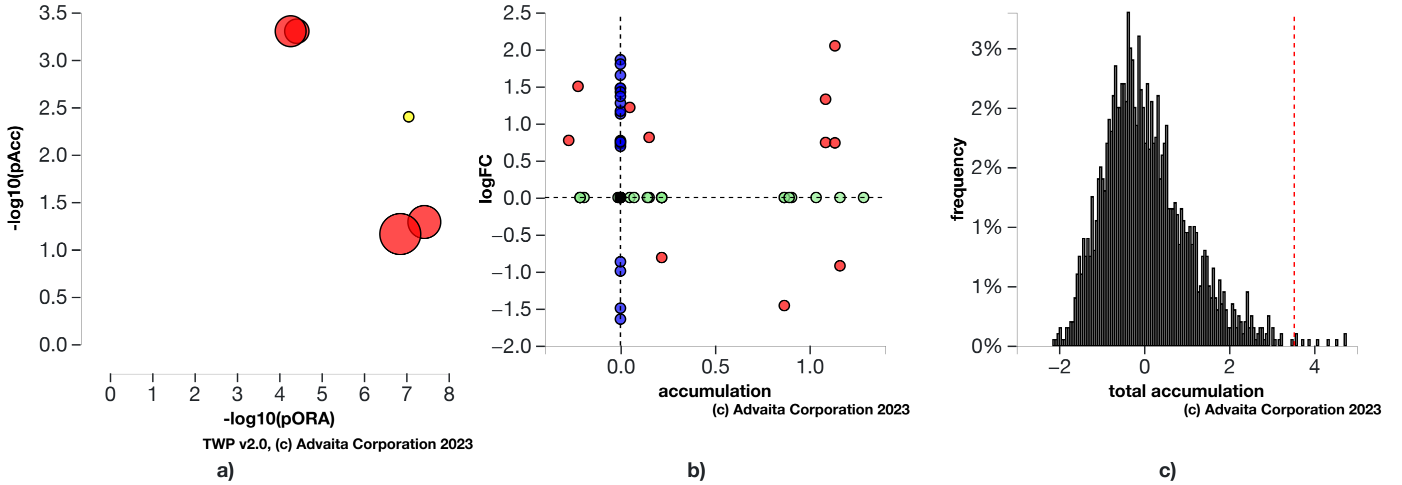


**Fig. 2.2.3: a) Perturbation vs over-representation:** *ECM-receptor interaction (KEGG: 04512) (yellow) is shown, using negative log of the accumulation and over-representation p-values, along with the other most significant pathways. Pathways in red are significant based on the combined uncorrected p-values, whereas the ones in black are non-significant (where applicable).* **b) Gene measured expression vs accumulation:** *All the genes from this pathway are represented in terms of their measured fold change (y-axis) and accumulation (x-axis). Accumulation is the perturbation received by the gene from any upstream genes. Genes displayed in red had both accumulation and measured fold change.Genes in blue had only measured fold change.Genes in green had only accumulation.The remaining genes that were not measured and had no accumulation are shown in black.* **c) Bootstrap diagram:** *The perturbation p-value is computed using bootstrap analysis. Bootstrapping assesses the probability of observing a sum of all absolute gene accumulation total accumulation at least as extreme as the computed one just by chance. A null distribution (gray bars) is computed through an iterative process that is repeated 2000 times. At each iteration, a number of genes equal to the number of differentially expressed genes in this pathway is randomly assigned anywhere in the pathway and the total accumulation is recomputed. The red line indicates the observed total accumulation of genes in the given pathway in relation to the distribution of expected values. The perturbation p-value is more significant the further away from the mean it is.*

#### Focal adhesion (KEGG: 04510)

Cell-matrix adhesions play essential roles in important biological processes including cell motility, cell proliferation, cell differentiation, regulation of gene expression and cell survival. At the cell-extracellular matrix contact points, specialized structures are formed and termed focal adhesions, where bundles of actin filaments are anchored to transmembrane receptors of the integrin family through a multi-molecular complex of junctional plaque proteins. Some of the constituents of focal adhesions participate in the structural link between membrane receptors and the actin cytoskeleton, while others are signalling molecules, including different protein kinases and phosphatases, their substrates, and various adapter proteins. Integrin signaling is dependent upon the non-receptor tyrosine kinase activities of the FAK and src proteins as well as the adaptor protein functions of FAK, src and Shc to initiate downstream signaling events. These signalling events culminate in reorganization of the actin cytoskeleton; a prerequisite for changes in cell shape and motility, and gene expression. Similar morphological alterations and modulation of gene expression are initiated by the binding of growth factors to their respective receptors, emphasizing the considerable crosstalk between adhesion- and growth factor-mediated signalling.

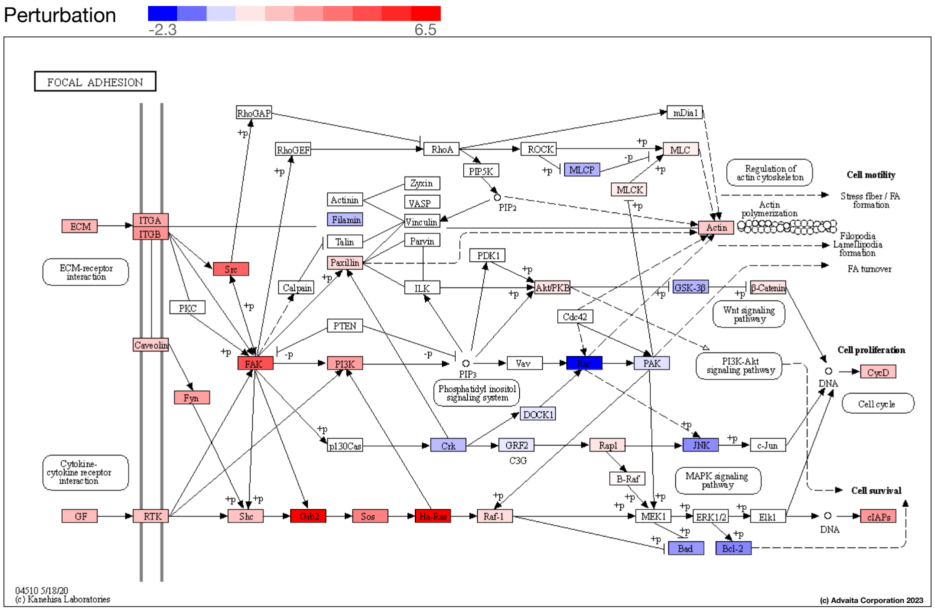


**Fig. 2.2.4: Focal adhesion (KEGG: 04510):** *The pathway diagram is overlayed with the computed perturbation of each gene. The perturbation accounts both for the gene's measured fold change and for the accumulated perturbation propagated from any upstream genes (accumulation). The highest negative perturbation is shown in dark blue, while the highest positive perturbation in dark red. The legend describes the values on the gradient. Note: For legibility, one gene may be represented in multiple places in the diagram and one box may represent multiple genes in the same gene family. A gene is highlighted in all locations it occurs in the diagram. For each gene family, the color corresponding to the gene with the highest absolute perturbation is displayed.*

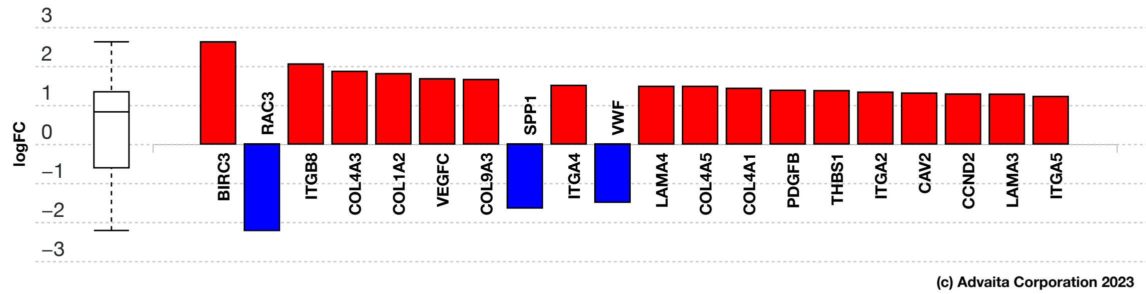


**Fig. 2.2.5: Gene measured expression bar plot:** *All the differentially expressed genes in Focal adhesion (KEGG: 04510) are ranked based on their absolute value of log fold change. The plot is limited to the top 20 genes out of a total of 48 differentially expressed genes. Upregulated genes are shown in red, downregulated genes are shown in blue. The box and whisker plot on the left summarizes the distribution of all the differentially expressed genes in this pathway. The box represents the 1st quartile, the median and the 3rd quartile, while the outliers are represented by circles.*

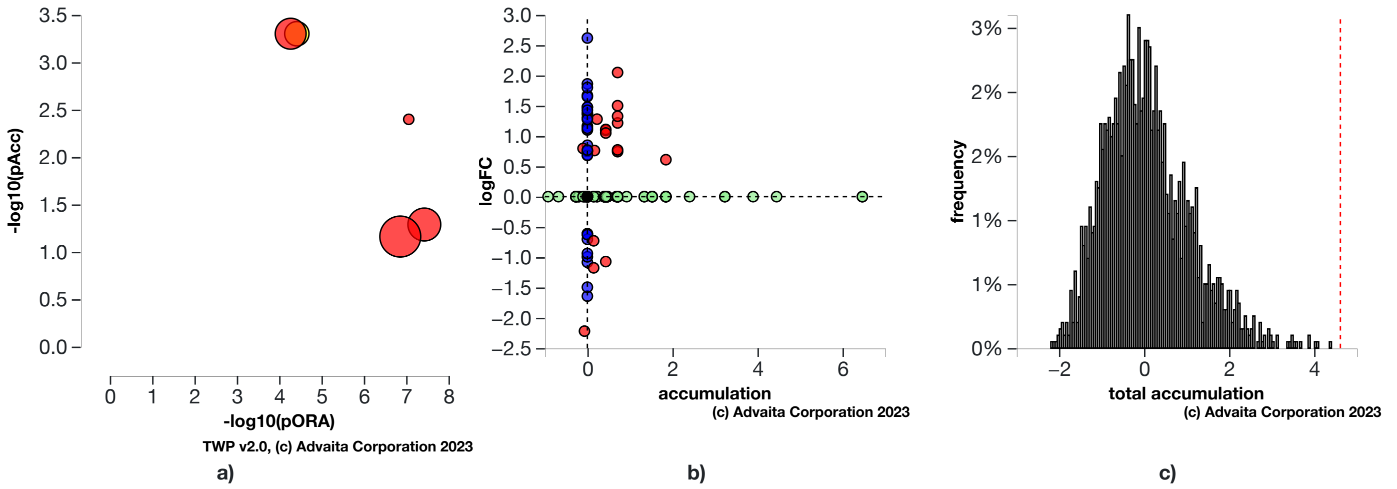


**Fig. 2.2.6: a) Perturbation vs over-representation:** *Focal adhesion (KEGG: 04510) (yellow) is shown, using negative log of the accumulation and over-representation p-values, along with the other most significant pathways. Pathways in red are significant based on the combined uncorrected p-values, whereas the ones in black are non-significant (where applicable).* **b) Gene measured expression vs accumulation:** *All the genes from this pathway are represented in terms of their measured fold change (y-axis) and accumulation (x-axis). Accumulation is the perturbation received by the gene from any upstream genes. Genes displayed in red had both accumulation and measured fold change.Genes in blue had only measured fold change.Genes in green had only accumulation.The remaining genes that were not measured and had no accumulation are shown in black.* **c) Bootstrap diagram:** *The perturbation p-value is computed using bootstrap analysis. Bootstrapping assesses the probability of observing a sum of all absolute gene accumulation total accumulation at least as extreme as the computed one just by chance. A null distribution (gray bars) is computed through an iterative process that is repeated 2000 times. At each iteration, a number of genes equal to the number of differentially expressed genes in this pathway is randomly assigned anywhere in the pathway and the total accumulation is recomputed. The red line indicates the observed total accumulation of genes in the given pathway in relation to the distribution of expected values. The perturbation p-value is more significant the further away from the mean it is.*

#### Cytokine-cytokine receptor interaction (KEGG: 04060)

Cytokines are soluble extracellular proteins or glycoproteins that are crucial intercellular regulators and mobilizers of cells engaged in innate as well as adaptive inflammatory host defenses, cell growth, differentiation, cell death, angiogenesis, and development and repair processes aimed at the restoration of homeostasis. Cytokines are released by various cells in the body, usually in response to an activating stimulus, and they induce responses through binding to specific receptors on the cell surface of target cells. Cytokines can be grouped by structure into different families and their receptors can likewise be grouped.

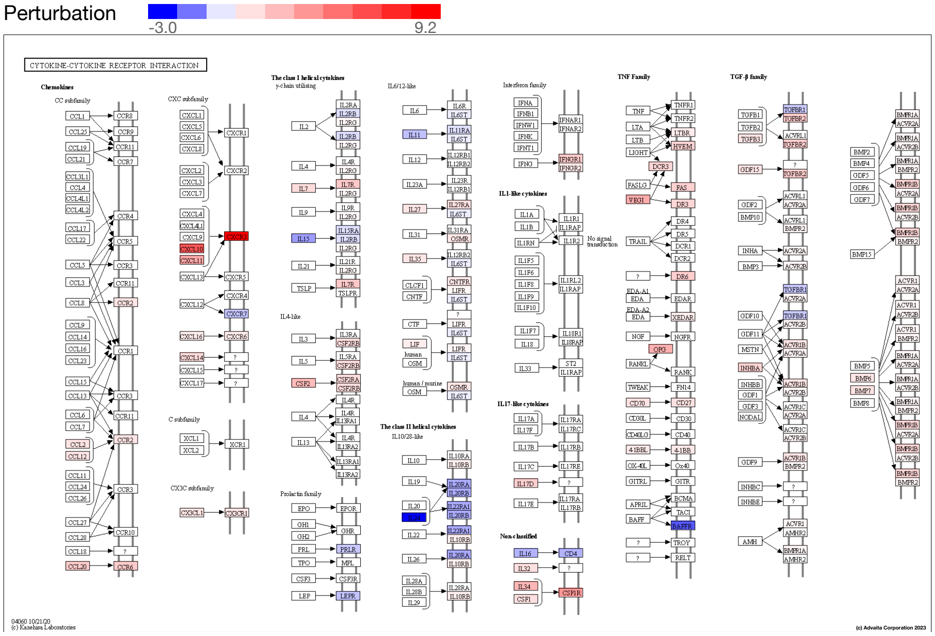


**Fig. 2.2.7: Cytokine-cytokine receptor interaction (KEGG: 04060):** *The pathway diagram is overlayed with the computed perturbation of each gene. The perturbation accounts both for the gene's measured fold change and for the accumulated perturbation propagated from any upstream genes (accumulation). The highest negative perturbation is shown in dark blue, while the highest positive perturbation in dark red. The legend describes the values on the gradient. Note: For legibility, one gene may be represented in multiple places in the diagram and one box may represent multiple genes in the same gene family. A gene is highlighted in all locations it occurs in the diagram. For each gene family, the color corresponding to the gene with the highest absolute perturbation is displayed.*

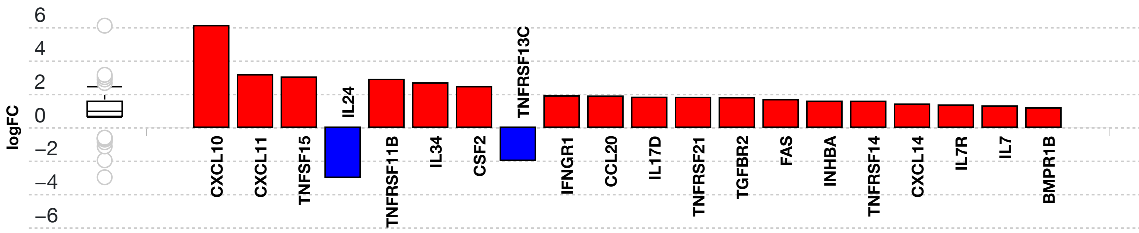


**(c) Advaita Corporation 2023**

**Fig. 2.2.8: Gene measured expression bar plot:** *All the differentially expressed genes in Cytokine-cytokine receptor interaction (KEGG: 04060) are ranked based on their absolute value of log fold change. The plot is limited to the top 20 genes out of a total of 49 differentially expressed genes. Upregulated genes are shown in red, downregulated genes are shown in blue. The box and whisker plot on the left summarizes the distribution of all the differentially expressed genes in this pathway. The box represents the 1st quartile, the median and the 3rd quartile, while the outliers are represented by circles.*

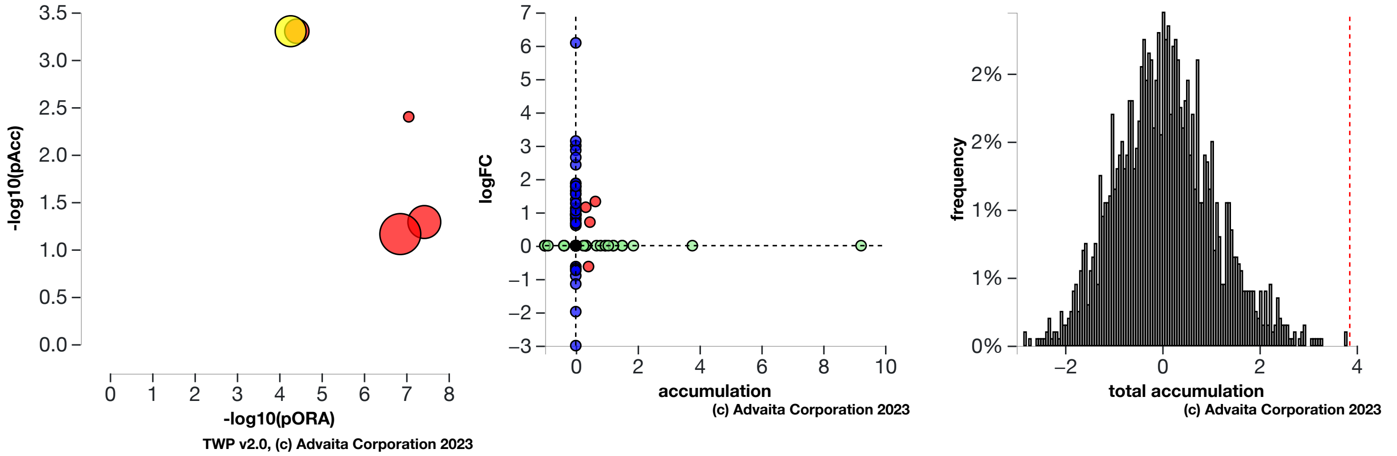


a) b) c)

**Fig. 2.2.9: a) Perturbation vs over-representation:** *Cytokine-cytokine receptor interaction (KEGG: 04060) (yellow) is shown, using negative log of the accumulation and over-representation p-values, along with the other most significant pathways. Pathways in red are significant based on the combined uncorrected p-values, whereas the ones in black are non-significant (where applicable).* **b) Gene measured expression vs accumulation:** *All the genes from this pathway are represented in terms of their measured fold change (y-axis) and accumulation (x-axis). Accumulation is the perturbation received by the gene from any upstream genes. Genes displayed in red had both accumulation and measured fold change.Genes in blue had only measured fold change.Genes in green had only accumulation.The remaining genes that were not measured and had no accumulation are shown in black.* **c) Bootstrap diagram:** *The perturbation p-value is computed using bootstrap analysis. Bootstrapping assesses the probability of observing a sum of all absolute gene accumulation total accumulation at least as extreme as the computed one just by chance. A null distribution (gray bars) is computed through an iterative process that is repeated 2000 times. At each iteration, a number of genes equal to the number of differentially expressed genes in this pathway is randomly assigned anywhere in the pathway and the total accumulation is recomputed. The red line indicates the observed total accumulation of genes in the given pathway in relation to the distribution of expected values. The perturbation p-value is more significant the further away from the mean it is.*

#### Human papillomavirus infection (KEGG: 05165)

Human papillomavirus (HPV) is a non-enveloped, double-stranded DNA virus. HPV infect mucoal and cutaneous epithelium resulting in several types of pathologies, most notably, cervical cancer. All types of HPV share a common genomic structure and encode eight proteins: E1, E2, E4, E5, E6, and E7 (early) and L1 and L2 (late). It has been demonstrated that E1 and E2 are involved in viral transcription and replication. The functions of the E4 protein is not yet fully understood. E5, E6, and E7 act as oncoproteins. E5 inhibits the V-ATPase, prolonging EGFR signaling and thereby promoting cell proliferation. The expression of E6 and E7 not only inhibits the tumor suppressors p53 and Rb, but also alters additional signalling pathways. Among these pathways, PI3K/Akt signalling cascade plays a very important role in HPV-induced carcinogenesis. The L1 and L2 proteins form icosahedral capsids for progeny virion generation.

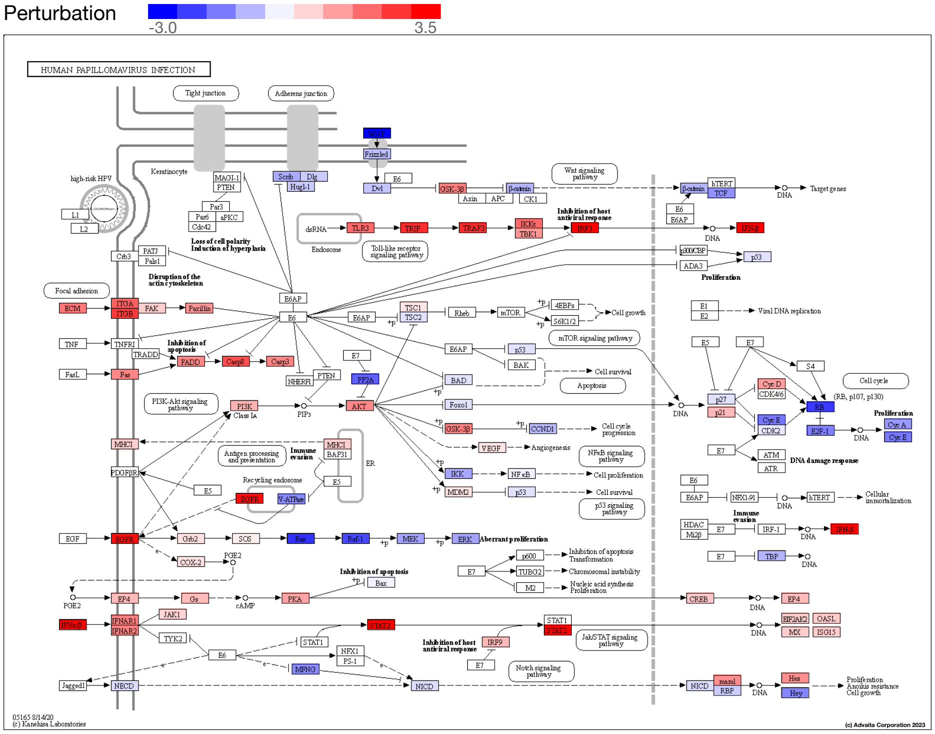


**Fig. 2.2.10: Human papillomavirus infection (KEGG: 05165):** *The pathway diagram is overlayed with the computed perturbation of each gene. The perturbation accounts both for the gene's measured fold change and for the accumulated perturbation propagated from any upstream genes (accumulation). The highest negative perturbation is shown in dark blue, while the highest positive perturbation in dark red. The legend describes the values on the gradient. Note: For legibility, one gene may be represented in multiple places in the diagram and one box may represent multiple genes in the same gene family. A gene is highlighted in all locations it occurs in the diagram. For each gene family, the color corresponding to the gene with the highest absolute perturbation is displayed.*

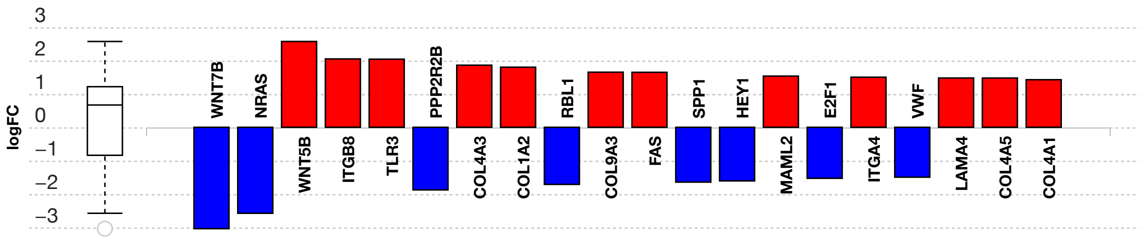


**(c) Advaita Corporation 2023**

**Fig. 2.2.11: Gene measured expression bar plot:** *All the differentially expressed genes in Human papillomavirus infection (KEGG: 05165) are ranked based on their absolute value of log fold change. The plot is limited to the top 20 genes out of a total of 77 differentially expressed genes. Upregulated genes are shown in red, downregulated genes are shown in blue. The box and whisker plot on the left summarizes the distribution of all the differentially expressed genes in this pathway. The box represents the 1st quartile, the median and the 3rd quartile, while the outliers are represented by circles.*

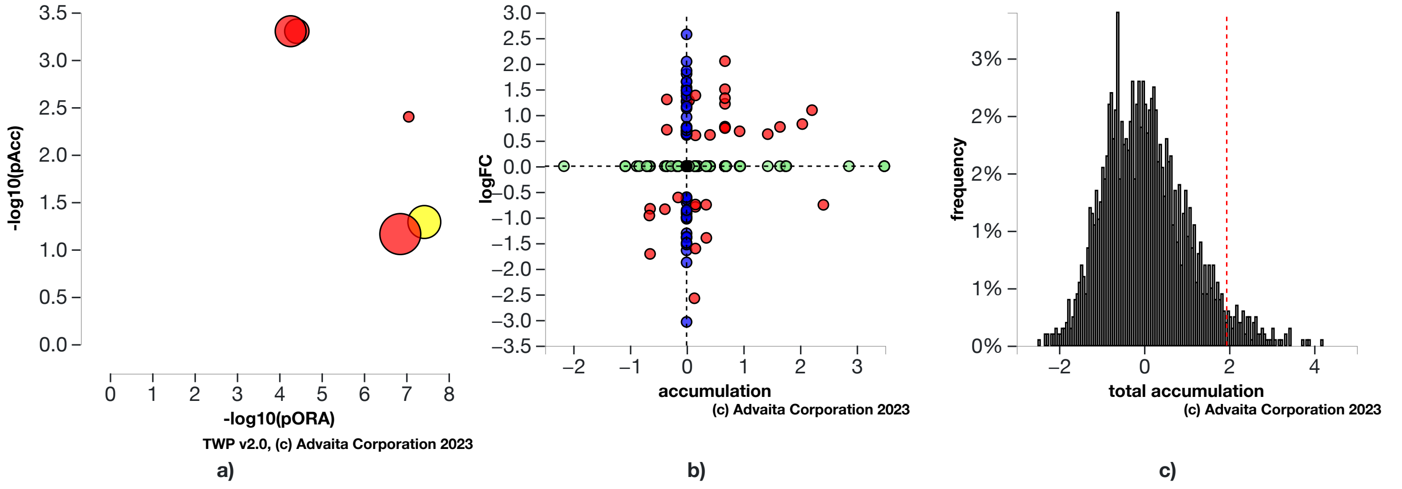


**Fig. 2.2.12: a) Perturbation vs over-representation:** *Human papillomavirus infection (KEGG: 05165) (yellow) is shown, using negative log of the accumulation and over-representation p-values, along with the other most significant pathways. Pathways in red are significant based on the combined uncorrected p-values, whereas the ones in black are non-significant (where applicable).* **b) Gene measured expression vs accumulation:** *All the genes from this pathway are represented in terms of their measured fold change (y-axis) and accumulation (x-axis). Accumulation is the perturbation received by the gene from any upstream genes. Genes displayed in red had both accumulation and measured fold change.Genes in blue had only measured fold change.Genes in green had only accumulation.The remaining genes that were not measured and had no accumulation are shown in black.* **c) Bootstrap diagram:** *The perturbation p-value is computed using bootstrap analysis. Bootstrapping assesses the probability of observing a sum of all absolute gene accumulation total accumulation at least as extreme as the computed one just by chance. A null distribution (gray bars) is computed through an iterative process that is repeated 2000 times. At each iteration, a number of genes equal to the number of differentially expressed genes in this pathway is randomly assigned anywhere in the pathway and the total accumulation is recomputed. The red line indicates the observed total accumulation of genes in the given pathway in relation to the distribution of expected values. The perturbation p-value is more significant the further away from the mean it is.*

#### Pathways in cancer (KEGG: 05200)

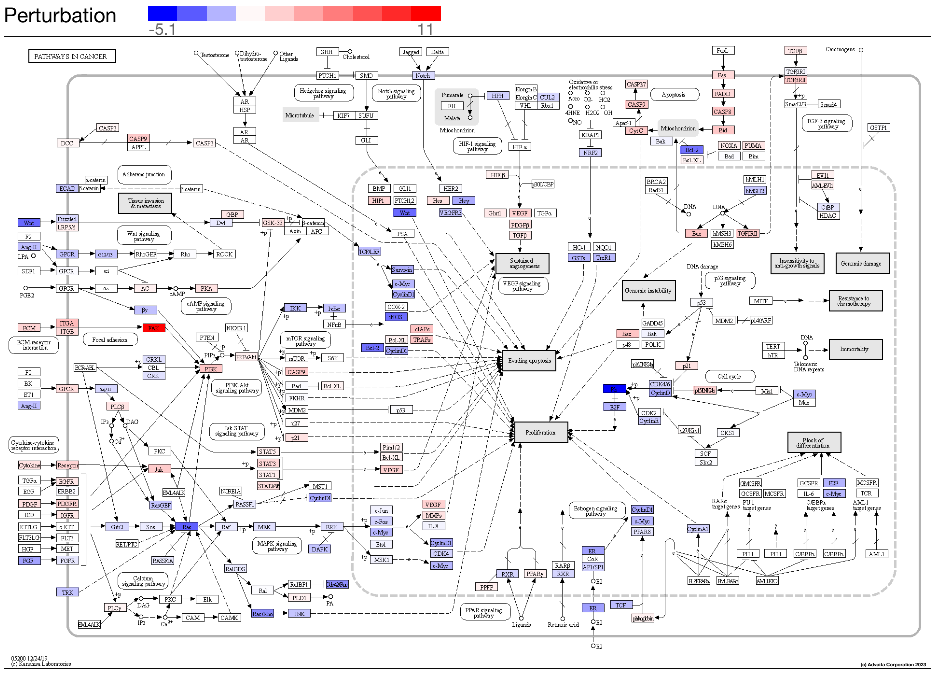


**Fig. 2.2.13: Pathways in cancer (KEGG: 05200):** *The pathway diagram is overlayed with the computed perturbation of each gene. The perturbation accounts both for the gene's measured fold change and for the accumulated perturbation propagated from any upstream genes (accumulation). The highest negative perturbation is shown in dark blue, while the highest positive perturbation in dark red. The legend describes the values on the gradient. Note: For legibility, one gene may be represented in multiple places in the diagram and one box may represent multiple genes in the same gene family. A gene is highlighted in all locations it occurs in the diagram. For each gene family, the color corresponding to the gene with the highest absolute perturbation is displayed.*

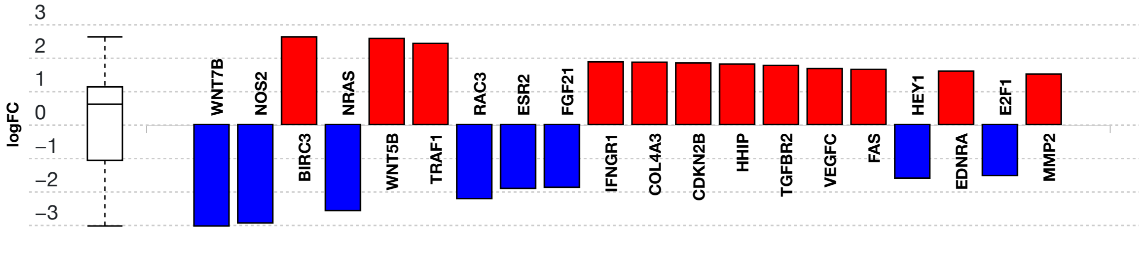


**Fig. 2.2.14: Gene measured expression bar plot:** *All the differentially expressed genes in Pathways in cancer (KEGG: 05200) are ranked based on their absolute value of log fold change. The plot is limited to the top 20 genes out of a total of 108 differentially expressed genes. Upregulated genes are shown in red, downregulated genes are shown in blue. The box and whisker plot on the left summarizes the distribution of all the differentially expressed genes in this pathway. The box represents the 1st quartile, the median and the 3rd quartile, while the outliers are represented by circles.*

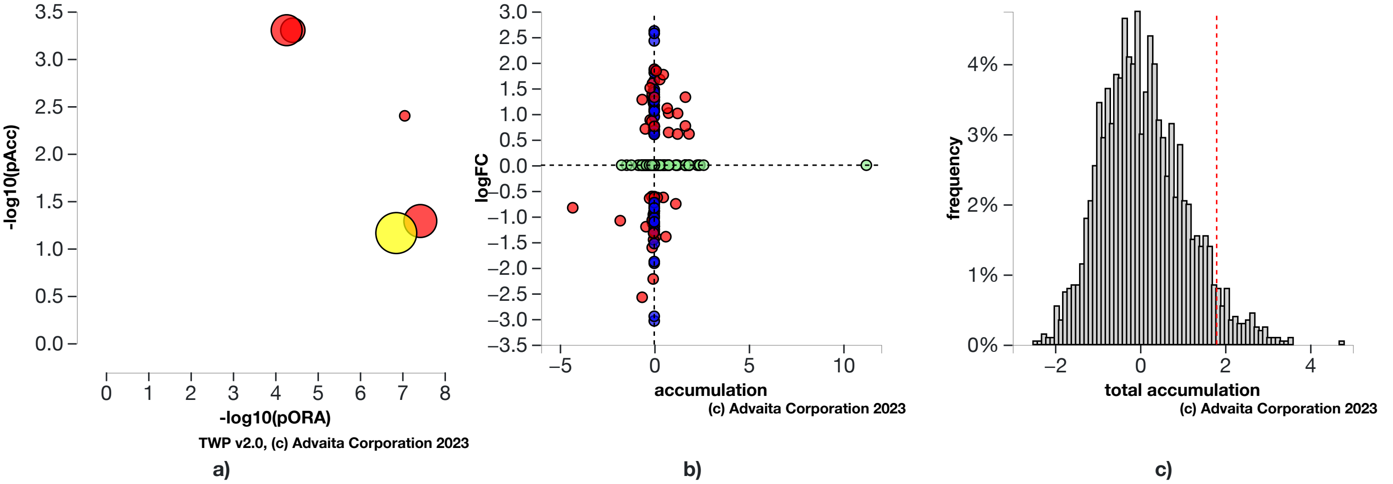


**Fig. 2.2.15: a) Perturbation vs over-representation:** *Pathways in cancer (KEGG: 05200) (yellow) is shown, using negative log of the accumulation and over-representation p-values, along with the other most significant pathways. Pathways in red are significant based on the combined uncorrected p-values, whereas the ones in black are non-significant (where applicable).* **b) Gene measured expression vs accumulation:** *All the genes from this pathway are represented in terms of their measured fold change (y-axis) and accumulation (x-axis). Accumulation is the perturbation received by the gene from any upstream genes. Genes displayed in red had both accumulation and measured fold change.Genes in blue had only measured fold change.Genes in green had only accumulation.The remaining genes that were not measured and had no accumulation are shown in black.* **c) Bootstrap diagram:** *The perturbation p-value is computed using bootstrap analysis. Bootstrapping assesses the probability of observing a sum of all absolute gene accumulation total accumulation at least as extreme as the computed one just by chance. A null distribution (gray bars) is computed through an iterative process that is repeated 2000 times. At each iteration, a number of genes equal to the number of differentially expressed genes in this pathway is randomly assigned anywhere in the pathway and the total accumulation is recomputed. The red line indicates the observed total accumulation of genes in the given pathway in relation to the distribution of expected values. The perturbation p-value is more significant the further away from the mean it is.*

### 3. Gene Ontology Analysis

#### 3.1. Methods

For each Gene Ontology (GO) term (Ashburner *et al*., 2002; Gene Ontology Consortium, 2004), the number of differentially expressed (DE) genes annotated to the term is compared to the number of DE genes expected just by chance. iPathwayGuide uses an over-representation approach to compute the statistical significance of observing at least the given number of DE genes. The p-value is computed using the hypergeometric distribution as described for pORA in the Pathway Analysis section. This p-value is corrected for multiple comparisons using FDR and Bonferroni.

The classical enrichment method used above considers all GO terms to be independent. By definition, all genes annotated to a GO term are also annotated to its ancestors. Because of this, the enrichment approach counts each gene multiple times by propagating it through the GO hierarchy from the most specific term the gene is associated with, all the way to the root of the ontology. This introduces redundancy in the analysis and reports many general and non-informative terms as significant. To overcome this limitation, iPathwayGuide allows users to use two more sophisticated pruning methods: *high-specificity pruning* and *smallest common denominator pruning*. The **high-specificity** pruning method *identifies the most specific GO terms* that are significantly associated with the set of DE genes. Let us consider, BP1 = “induction of apoptosis by intracellular signals” and BP2 = “induction of apoptosis by extracellular signals,” which are two of the children of BP3 = “induction of apoptosis.” If enough DE genes are associated with BP1 and BP2, the high-specificity pruning will report them as significant. The **smallest common denominator** pruning method *identifies the GO terms that best encapsulate the set of DE genes*, at times consolidating significance of two or more specific terms into their common parent. In the example above, this pruning method might report BP3 as significant because it is the most specific biological term that would include all DE genes that make both BP1 and BP2 significant.

#### 3.2. Biological Processes results

**Table 3.2.1: Top identified biological processes. Only the top scoring biological process for each pruning type is described below the table.**

| Pruning Type: None | | | | Pruning Type: High-specificity | | Pruning Type: Smallest Common Denominator | |
|---|---|---|---|---|---|---|---|
| **GO Term** | **p-value** | **p-value (FDR)** | **p-value (Bonferroni)** | **GO Term** | **p-value** | **GO Term** | **p-value** |
| multicellular organism development | **5.800e-11** | **2.605e-7** | **6.104e-7** | extracellular matrix organization | **4.946e-4** | extracellular matrix organization | **1.789e-5** |
| system development | **6.100e-11** | **2.605e-7** | **6.420e-7** | cellular response to retinoic acid | 0.126 | response to retinoic acid | 0.068 |
| extracellular matrix organization | **8.300e-11** | **2.605e-7** | **8.735e-7** | endodermal cell differentiation | 0.126 | angiogenesis | 0.176 |
| extracellular structure organization | **9.900e-11** | **2.605e-7** | **1.042e-6** | regulation of transcription involved in G1/S transition of mitotic cell cycle | 0.176 | regulation of transcription involved in G1/S transition of mitotic cell cycle | 0.176 |
| anatomical structure development | **3.700e-10** | **7.788e-7** | **3.894e-6** | positive regulation of cell migration | 0.226 | ventricular septum morphogenesis | 0.210 |

##### multicellular organism development (GO:0007275)

The biological process whose specific outcome is the progression of a multicellular organism over time from an initial condition (e.g. a zygote or a young adult) to a later condition (e.g. a multicellular animal or an aged adult). In this experiment, the algorithm identified **929** differentially expressed gene(s) out of ALL **4,919** gene(s).

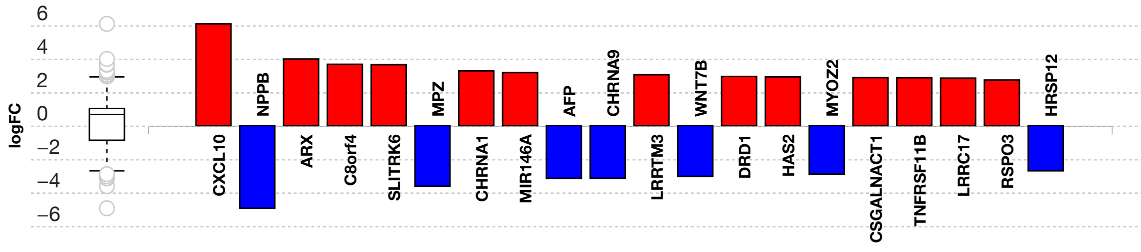


**Fig. 3.2.1: Gene measured expression bar plot:** *All the differentially expressed genes that are annotated to multicellular organism development are ranked based on their absolute value of log fold change. The plot is limited to the top 20 genes out of a total of 929 differentially expressed genes. Upregulated genes are shown in red, downregulated genes are shown in blue. The box and whisker plot on the left summarizes the distribution of all the differentially expressed genes that are annotated to this GO term. The box represents the 1st quartile, the median and the 3rd quartile, while the outliers are represented by circles.*

##### extracellular matrix organization (GO:0030198)

A process that is carried out at the cellular level which results in the assembly, arrangement of constituent parts, or disassembly of an extracellular matrix. In this experiment, the algorithm identified **108** differentially expressed gene(s) out of ALL **369** gene(s).

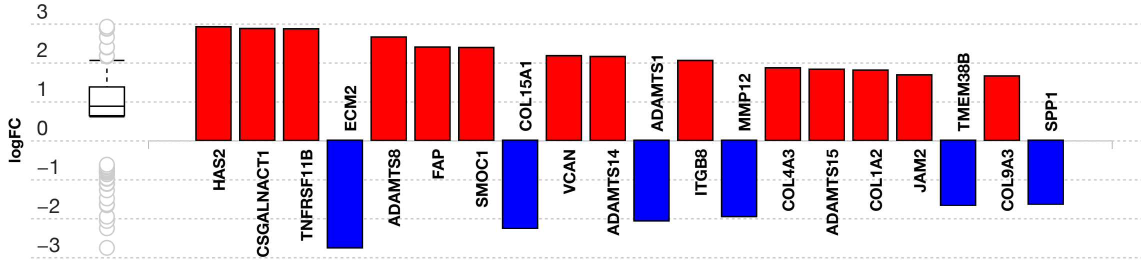


**Fig. 3.2.2: Gene measured expression bar plot:** *All the differentially expressed genes that are annotated to extracellular matrix organization are ranked based on their absolute value of log fold change. The plot is limited to the top 20 genes out of a total of 108 differentially expressed genes. Upregulated genes are shown in red, downregulated genes are*

*shown in blue. The box and whisker plot on the left summarizes the distribution of all the differentially expressed genes that are annotated to this GO term. The box represents the 1st quartile, the median and the 3rd quartile, while the outliers are represented by circles.*

#### 3.3. Molecular Functions results

**Table 3.3.1: Top identified molecular functions. Only the top scoring molecular function for each pruning type is described below the table.**

| Pruning Type: None | | | | Pruning Type: High-specificity | | Pruning Type: Smallest Common Denominator | |
|---|---|---|---|---|---|---|---|
| **GO Term** | **p-value** | **p-value (FDR)** | **p-value (Bonferroni)** | **GO Term** | **p-value** | **GO Term** | **p-value** |
| extracellular matrix structural constituent | **7.700e-7** | **0.002** | **0.002** | extracellular matrix structural constituent conferring tensile strength | **0.177** | extracellular matrix structural constituent | **0.002** |
| extracellular matrix structural constituent conferring tensile strength | **7.400e-5** | 0.089 | 0.177 | integrin binding | 0.180 | integrin binding | 0.180 |
| integrin binding | **1.500e-4** | 0.120 | 0.359 | DNA replication origin binding | 0.327 | identical protein binding | 0.196 |
| protein binding | **2.000e-4** | 0.120 | 0.479 | extracellular matrix structural constituent | 0.335 | DNA replication origin binding | 0.196 |
| signaling receptor binding | **3.100e-4** | 0.123 | 0.742 | chemorepellent activity | 0.335 | TAP binding | 0.196 |

##### extracellular matrix structural constituent (GO:0005201)

The action of a molecule that contributes to the structural integrity of the extracellular matrix. In this experiment, the algorithm identified **49** differentially expressed gene(s) out of ALL **152** gene(s).

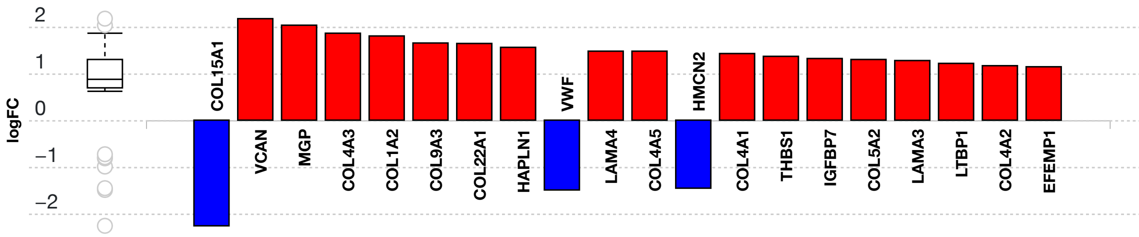


**Fig. 3.3.3: Gene measured expression bar plot:** *All the differentially expressed genes that are annotated to extracellular matrix structural constituent are ranked based on their absolute value of log fold change. The plot is limited to the top 20 genes out of a total of 49 differentially expressed genes. Upregulated genes are shown in red, downregulated genes are shown in blue. The box and whisker plot on the left summarizes the distribution of all the differentially expressed genes that are annotated to this GO term. The box represents the 1st quartile, the median and the 3rd quartile, while the outliers are represented by circles.*

##### extracellular matrix structural constituent conferring tensile strength (GO:0030020)

A constituent of the extracellular matrix that enables the matrix to resist longitudinal stress. In this experiment, the algorithm identified **17** differentially expressed gene(s) out of ALL **40** gene(s).

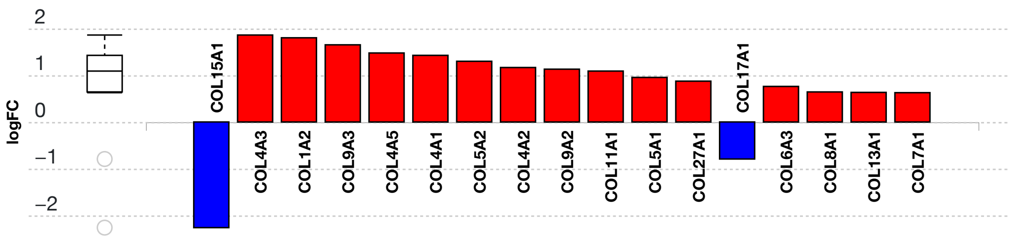


**Fig. 3.3.4: Gene measured expression bar plot:** *All the differentially expressed genes that are annotated to extracellular matrix structural constituent conferring tensile strength are ranked based on their absolute value of log fold change. Upregulated genes are shown in red, downregulated genes are shown in blue. The box and whisker plot on the left summarizes the distribution of all the differentially expressed genes that are annotated to this GO term. The box represents the 1st quartile, the median and the 3rd quartile, while the outliers are represented by circles.*

#### 3.4. Cellular Components results

**Table 3.4.1: Top identified cellular components. Only the top scoring cellular component for each pruning type is described below the table.**

| Pruning Type: None | | | | Pruning Type: High-specificity | | Pruning Type: Smallest Common Denominator | |
|---|---|---|---|---|---|---|---|
| GO Term | p-value | p-value (FDR) | p-value (Bonferroni) | GO Term | p-value | GO Term | p-value |
| endoplasmic reticulum lumen | 4.600e-7 | 4.822e-4 | 5.837e-4 | endoplasmic reticulum lumen | 5.837e-4 | endoplasmic reticulum lumen | 0.001 |
| extracellular matrix | 8.500e-7 | 4.822e-4 | 0.001 | MCM complex | 0.006 | MCM complex | 0.006 |
| cell periphery | 1.200e-6 | 4.822e-4 | 0.002 | glutamatergic synapse | 0.161 | basement membrane | 0.007 |
| plasma membrane | 1.700e-6 | 4.822e-4 | 0.002 | integral component of membrane | 0.209 | extracellular space | 0.028 |
| basement membrane | 1.900e-6 | 4.822e-4 | 0.002 | basement membrane | 0.236 | integral component of postsynaptic membrane | 0.028 |

##### endoplasmic reticulum lumen (GO:0005788)

The volume enclosed by the membranes of the endoplasmic reticulum. In this experiment, the algorithm identified **79** differentially expressed gene(s) out of ALL **285** gene(s).

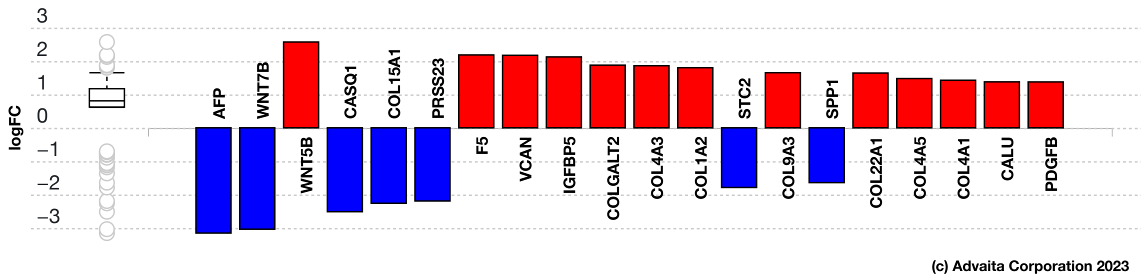


**Fig. 3.4.5: Gene measured expression bar plot:** *All the differentially expressed genes that are annotated to endoplasmic reticulum lumen are ranked based on their absolute value of log fold change. The plot is limited to the top 20 genes out of a total of 79 differentially expressed genes. Upregulated genes are shown in red, downregulated genes are shown in blue. The box and whisker plot on the left summarizes the distribution of all the differentially expressed genes that are annotated to this GO term. The box represents the 1st quartile, the median and the 3rd quartile, while the outliers are represented by circles.*

### 4. Predicted Upstream Regulator Analysis - miRNAs

#### 4.1. Methods

The prediction of active miRNAs (Friedman *et al*., 2009; Lewis *et al*., 2005) is based on enrichment of differentially downregulated target genes of the miRNAs. In general, miRNAs have an inhibitory effect on their targets. Therefore, for any given miRNA the method computes the ratio between the number of differentially downregulated targets and all differentially expressed targets, and compares it to the ratio of all downwardly expressed targets to all targets. Overall, iPathwayGuide calculates the probability of observing at least the number of differentially downregulated target genes for a given miRNA just by chance. This p-value is computed using the hypergeometric distribution as described for pORA in the Pathway Analysis section.

#### 4.2. Results

**Table 4.2.1: Top identified miRNAs**

| miRNA Name | p-value | p-value (FDR) | p-value (Bonferroni) |
|---|---|---|---|
| hsa-miR-335-5p | **0.025** | 1.000 | 1.000 |
| hsa-miR-122-5p | **0.050** | 1.000 | 1.000 |
| hsa-miR-802 | 0.055 | 1.000 | 1.000 |
| hsa-miR-184 | 0.067 | 1.000 | 1.000 |
| hsa-miR-379-5p | 0.100 | 1.000 | 1.000 |

#### hsa-miR-335-5p (MIMAT0000765)

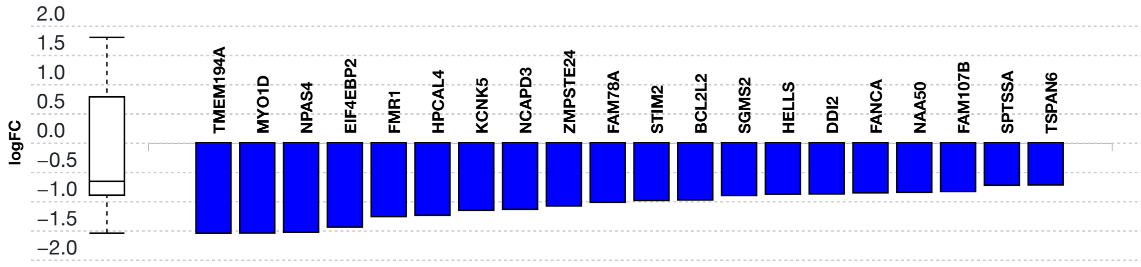


(c) Advaita Corporation 2023

**Fig. 4.2.1: Gene measured expression bar plot:** *All the differentially expressed genes that are targeted by hsa-miR-335-5p are ranked based on their measured expression change (most downregulated to upregulated). The downregulated genes are shown in blue, and the upregulated ones are shown in red (where applicable). The plot is limited to the top 20 genes out of a total of 50 differentially expressed target genes. Out of all the differentially expressed target genes, 30 were found to be downregulated. The box and whisker plot on the left summarizes the distribution of all the differentially expressed genes targeted by this miRNA. The box represents the 1st quartile, the median and the 3rd quartile, while the outliers are represented by circles.*

#### hsa-miR-122-5p (MIMAT0000421)

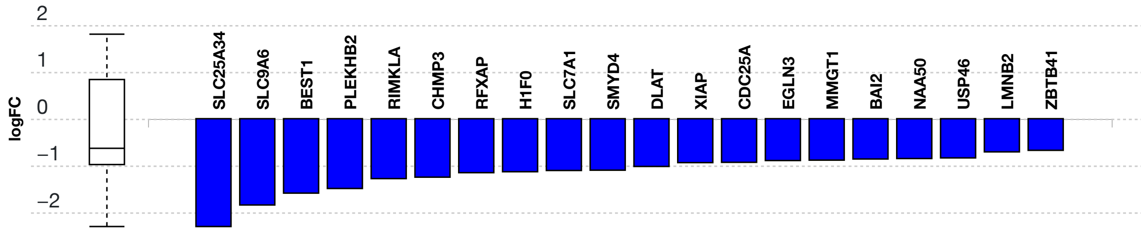


(c) Advaita Corporation 2023

**Fig. 4.2.2: Gene measured expression bar plot:** *All the differentially expressed genes that are targeted by hsa-miR-122-5p are ranked based on their measured expression change (most downregulated to upregulated). The downregulated genes are shown in blue, and the upregulated ones are shown in red (where applicable). The plot is limited to the top 20 genes out of a total of 43 differentially expressed target genes. Out of all the differentially expressed target genes, 23 were found to be downregulated. The box and whisker plot on the left summarizes the distribution of all the differentially expressed genes targeted by this miRNA. The box represents the 1st quartile, the median and the 3rd quartile, while the outliers are represented by circles.*

#### hsa-miR-802 (MIMAT0004185)

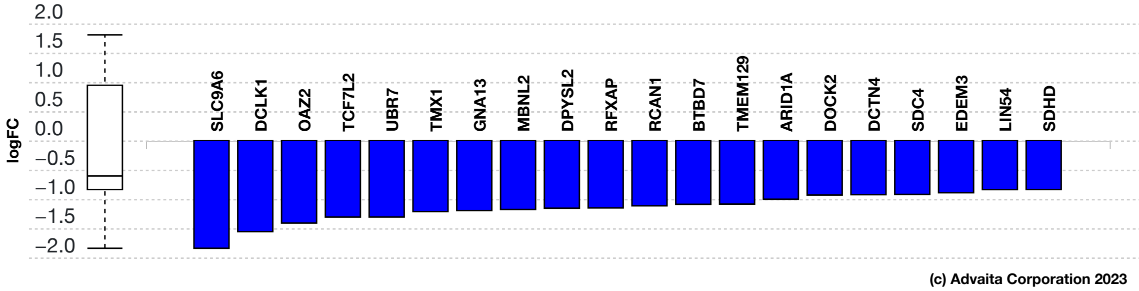


**Fig. 4.2.3: Gene measured expression bar plot:** *All the differentially expressed genes that are targeted by hsa-miR-802 are ranked based on their measured expression change (most downregulated to upregulated). The downregulated genes are shown in blue, and the upregulated ones are shown in red (where applicable). The plot is limited to the top 20 genes out of a total of 77 differentially expressed target genes. Out of all the differentially expressed target genes, 40 were found to be downregulated. The box and whisker plot on the left summarizes the distribution of all the differentially expressed genes targeted by this miRNA. The box represents the 1st quartile, the median and the 3rd quartile, while the outliers are represented by circles.*

#### hsa-miR-184 (MIMAT0000454)

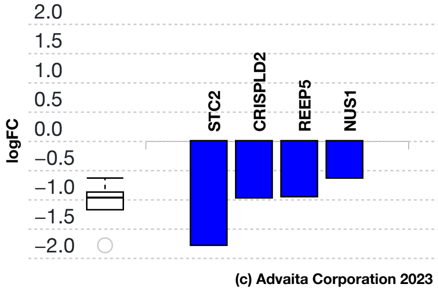


**Fig. 4.2.4: Gene measured expression bar plot:** *All the differentially expressed genes that are targeted by hsa-miR-184 are ranked based on their measured expression change (most downregulated to upregulated). The downregulated genes are shown in blue, and the upregulated ones are shown in red (where applicable). Out of all the differentially expressed target genes, 4 were found to be downregulated. The box and whisker plot on the left summarizes the distribution of all the differentially expressed genes targeted by this miRNA. The box represents the 1st quartile, the median and the 3rd quartile, while the outliers are represented by circles.*

#### hsa-miR-379-5p (MIMAT0000733)

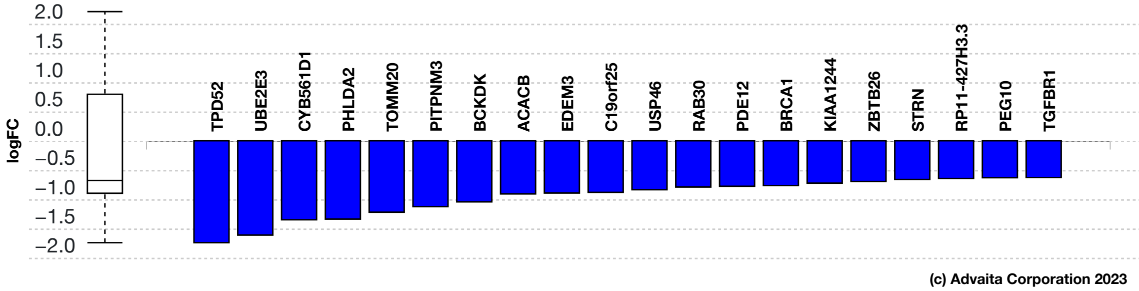


**Fig. 4.2.5: Gene measured expression bar plot:** *All the differentially expressed genes that are targeted by hsa-miR-379-5p are ranked based on their measured expression change (most downregulated to upregulated). The downregulated genes are shown in blue, and the upregulated ones are shown in red (where applicable). The plot is limited to the top 20 genes out of a total of 32 differentially expressed target genes. Out of all the differentially expressed target genes, 20 were found to be downregulated. The box and whisker plot on the left summarizes the distribution of all the differentially expressed genes targeted by this miRNA. The box represents the 1st quartile, the median and the 3rd quartile, while the outliers are represented by circles.*

### 5. Predicted Upstream Regulator Analysis - Genes

#### 5.1. Methods

The prediction of upstream regulators is based on two types of information: i) the enrichment of differentially expressed genes from the experiment and ii) a network of regulatory interactions from our proprietary knowledge base (see the report information for details). The network is a directed graph in which the nodes represent genes, and the edges represent regulatory interactions between two genes. A signed edge in this graph consists of a source gene, a target gene, and a sign to indicate the type of signal: activation (+) or inhibition (-). To create the network, the analysis selects only those edges observed in the literature with at least a medium confidence (evidence score greater than or equal to 400). The analysis considers two hypotheses:

HA. The upstream regulator is **activated** in the condition studied.
HI. The upstream regulator is **inhibited** in the condition studied.

The analysis divides the set of all the genes obtained from NCBI Gene database into several subsets based on the measurements in the experiment and the definitions shown in **Figure 5.1.1** and **Figure 5.1.2**. Let the sign of a measured DE gene be the sign of the log fold change value: (+) for up-regulated genes and (-) for down-regulated genes. A gene is a target gene if it corresponds to a node in the network that has at least one incoming edge. We define a *consistent gene* as a target DE gene such that the sign of the gene is consistent both with the type of the signal **and** with the hypothesis considered. Formally, by definition, a target DE gene *g* is consistent with Hypothesis HA if and only if an incoming edge *e* exists such that *sign(g) = sign(e)*. In other words, this describes the situation when the upstream regulator is predicted as activated, the signal is activation and the target DE gene is up-regulated, or the signal is inhibition and the target DE gene is down-regulated (see panel A in **Figure 5.1.1**). A target DE gene *g* is consistent with Hypothesis HI if and only if an incoming edge *e* exists such that *sign(g) ≠ sign(e)*. This second case captures the situation in which the upstream regulator is inhibited, the signal is inhibition and the target DE gene is up-regulated, or the signal is activation and the target DE gene is down-regulated (see panel B in **Figure 5.1.1**).

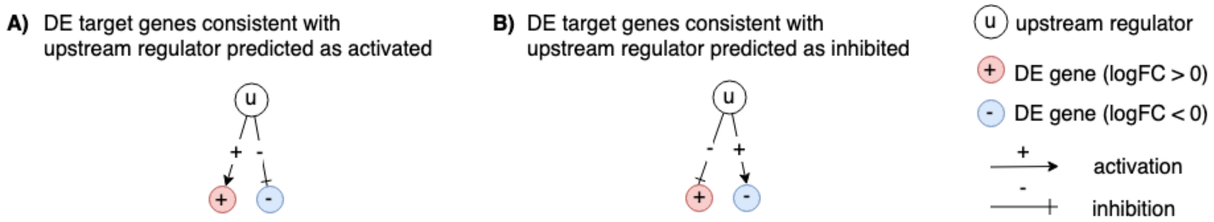


**Fig. 5.1.1: Target genes consistent with the hypothesis considered:** *In panel A, the signs of the DE genes match the signs of their respective incoming edges, increasing the likelihood that the upstream regulator u is activated. In panel B, the signs of the DE genes are opposite to the signs of their edges, increasing the likelihood that the upstream regulator u is inhibited.*

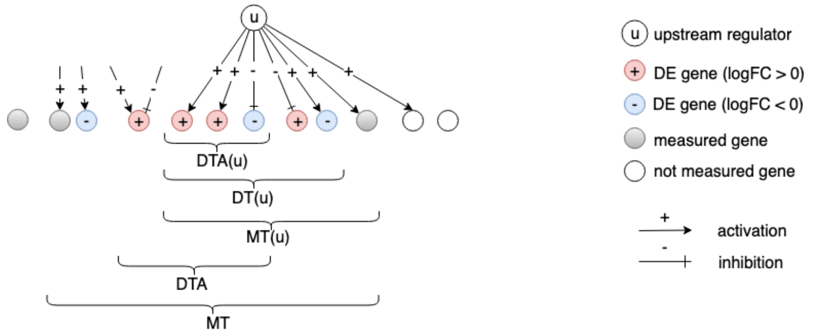


**Fig. 5.1.2:** *The set of all genes includes the set of measured genes that are also targets in the network, or Measured Targets (MT). We define the subset of "DE Targets consistent with the first hypothesis that the upstream regulators are Activated", DTA. For a selected upstream regulator u, we have the set of "Measured Targets of u" MT(u), "Differentially expressed Targets downstream of u" DT(u), and the set of "DE targets consistent with the hypothesis HA that u is Activated" DTA(u). The equivalent graphic for the hypothesis HI associated with DTI and DTI(u) is not shown.*

#### Upstream regulators Z-score

For both research hypotheses, the analysis computes a Z-score for each upstream regulator *z(u)* by iterating over the genes in DT(u) and their incoming edges *in(g)*. We can then compute the p-value corresponding to the z-score $P_z$ as the one-tailed area under the probability density function for a normal distribution, N(0,1).

#### Upstream regulators predicted as activated

Here, the research hypothesis considers the upstream regulator as activated. For each upstream regulator *u*, the number of consistent DE genes downstream of *u*, DTA(u) is compared to the number of measured target genes expected to be both consistent and DE just by chance. iPathwayGuide uses an over-representation approach to compute the statistical significance of observing at least the given number of consistent DE genes. The p-value $P_{act}$ is computed using the hypergeometric distribution (Draghici *et al*., 2003, Draghici 2011).

After computing a p-value for both types of evidence, $P_z$ and $P_{act}$, we need to combine these two probabilities into one global probability value, $P_G$ that is used to rank the upstream regulators and test the research hypothesis that the upstream regulators are predicted as activated in the condition studied. Since only a positive z-score indicates that the upstream regulator is predicted as activated, we only combine p-values for a positive z-score. Moreover, to avoid introducing false positives, only $P_z$ for significant z-scores ( $z \geq 2$ ) are combined. The analysis uses the standard Fisher's method to combine p-values into one test statistic (Fisher 1925).

#### Upstream regulators predicted as inhibited

In parallel with upstream regulators predicted as activated, we use $P_{inh}$ and $P_z$ to predict upstream regulators that are inhibited. Here, the research hypothesis states that the upstream regulators are inhibited in the conditions studied. For each upstream regulator *u*, the number of consistent DE genes downstream of *u*, DTI(u) is compared to the number of measured target genes expected to be both consistent and DE just by chance. Using the Fisher's method as above, the analysis combines $P_{inh}$ and $P_z$, where $P_z$ is considered only for significant negative z-scores ( $z \leq -2$ ).

#### 5.2. Results: upstream regulators predicted as activated

| Upstream Regulator (u) | DTA(u) | DT(u) | p-value | p-value (FDR) | p-value (Bonferroni) |
|---|---|---|---|---|---|
| TP53 | 57 | 70 | 5.683e-9 | 1.308e-5 | 1.308e-5 |
| FAM20C | 24 | 32 | 1.349e-5 | 0.012 | 0.031 |
| FAM20A | 24 | 32 | 1.583e-5 | 0.012 | 0.036 |
| E2F5 | 5 | 5 | 1.967e-4 | 0.113 | 0.453 |
| RBL1 | 4 | 4 | 7.437e-4 | 0.244 | 1.000 |

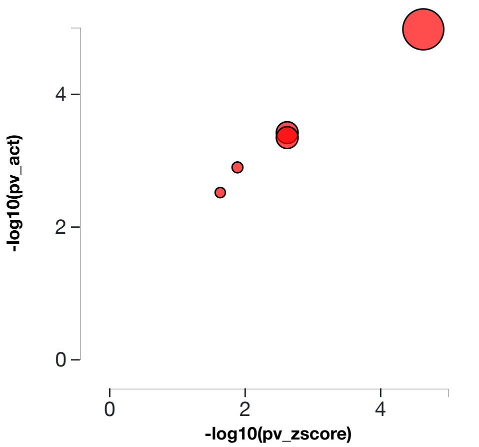


**Table 5.2.1: Top upstream regulators predicted as activated.** *For each upstream regulator u, the table shows the number of DE targets supporting the hypothesis that the regulator is activated DTA(u) the total number of DE genes downstream of u DT(u), the combined raw p-value, and the p-value corrected for multiple comparisons.* **Fig. 5.2.1: A two-way plot showing the top five upstream regulators predicted as activated.** *Dots representing upstream regulators are positioned using $P_{zscore}$ on the horizontal axis, and using $P_{act}$ on the vertical axis. $P_{act}$ is the p-value based on the number of DE targets consistent with the typeof the incoming signal and with the selected hypothesis type. Upstream regulators with a significant combined p-value are shown in red. The size of each dot represents the number of consistent DE genes for that regulator.*

##### TP53 (tumor protein p53)

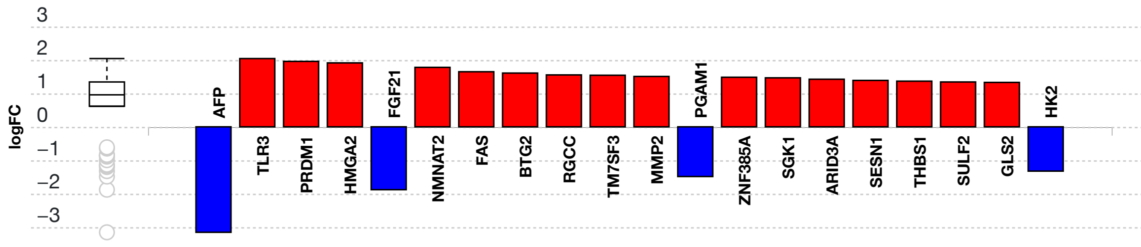


**Fig. 5.2.3: Gene measured expression bar plot:** *All the consistent differentially expressed genes that are targeted by TP53 are ranked based on their absolute value of log fold change. The plot is limited to the top 20 genes out of a total of 57 consistent differentially expressed target genes. Upregulated genes are shown in red, downregulated genes are shown in blue. The box and whisker plot on the left summarizes the distribution of all the consistent differentially expressed genes targeted by this upstream regulator. The box shows the 1st quartile, the median and the 3rd quartile, while the outliers are represented by circles.*

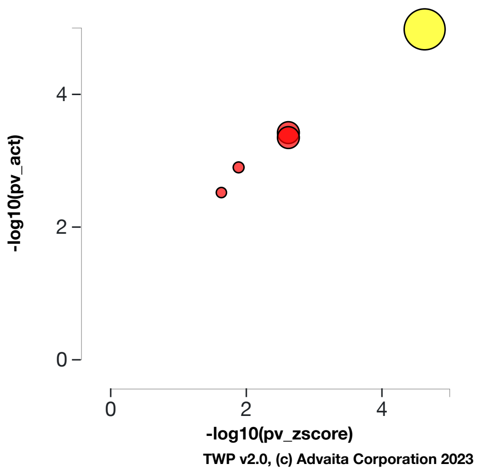


**Fig. 5.2.4: Activation p-value vs zscore p-value:** *TP53, tumor protein p53, (yellow) is shown, using negative log of the activation and zscore p-values, along with the other most significant upstream regulators. The size of the dot represents the relative number of consistent DE genes, which for selected upstream regulator is 57.*

#### FAM20C (FAM20C golgi associated secretory pathway kinase)

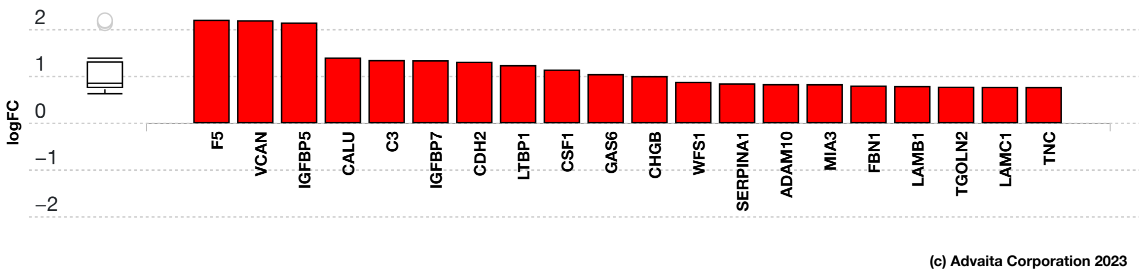


**Fig. 5.2.5: Gene measured expression bar plot:** *All the consistent differentially expressed genes that are targeted by FAM20C are ranked based on their absolute value of log fold change. The plot is limited to the top 20 genes out of a total of 24 consistent differentially expressed target genes. Upregulated genes are shown in red, downregulated genes are shown in blue. The box and whisker plot on the left summarizes the distribution of all the consistent differentially expressed genes targeted by this upstream regulator. The box shows the 1st quartile, the median and the 3rd quartile, while the outliers are represented by circles.*

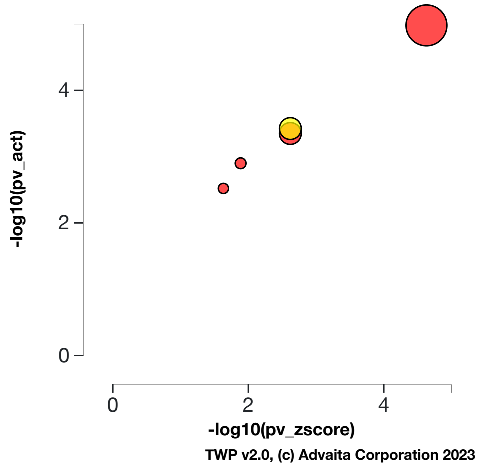


**Fig. 5.2.6: Activation p-value vs zscore p-value:** *FAM20C, FAM20C golgi associated secretory pathway kinase, (yellow) is shown, using negative log of the activation and zscore p-values, along with the other most significant upstream regulators. The size of the dot represents the relative number of consistent DE genes, which for selected upstream regulator is 24.*

#### FAM20A (FAM20A golgi associated secretory pathway pseudokinase)

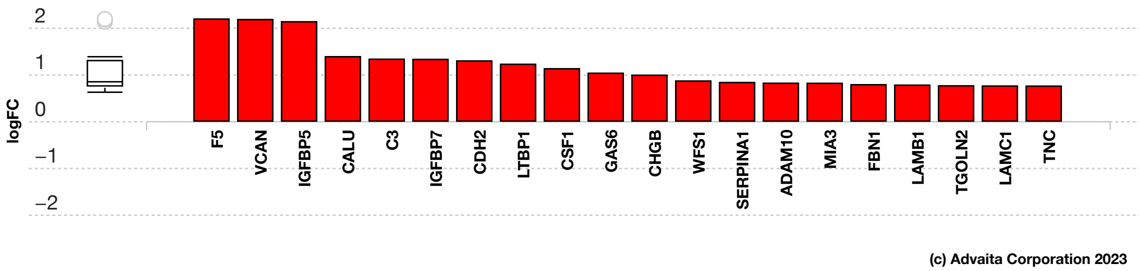


**Fig. 5.2.7: Gene measured expression bar plot:** *All the consistent differentially expressed genes that are targeted by FAM20A are ranked based on their absolute value of log fold change. The plot is limited to the top 20 genes out of a total of 24 consistent differentially expressed target genes. Upregulated genes are shown in red, downregulated genes are shown in blue. The box and whisker plot on the left summarizes the distribution of all the consistent differentially expressed genes targeted by this upstream regulator. The box shows the 1st quartile, the median and the 3rd quartile, while the outliers are represented by circles.*

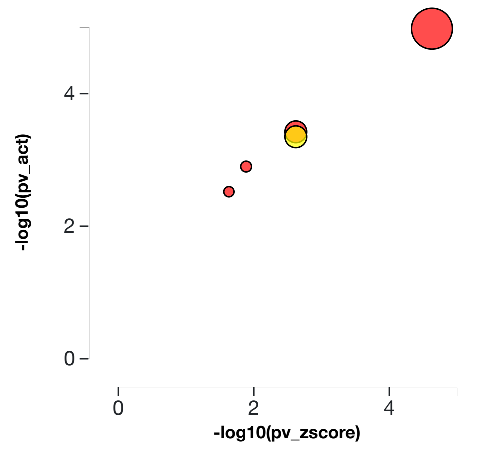


**Fig. 5.2.8: Activation p-value vs zscore p-value:** *FAM20A, FAM20A golgi associated secretory pathway pseudokinase, (yellow) is shown, using negative log of the activation and zscore p-values, along with the other most significant upstream regulators. The size of the dot represents the relative number of consistent DE genes, which for selected upstream regulator is 24.*

#### E2F5 (E2F transcription factor 5)

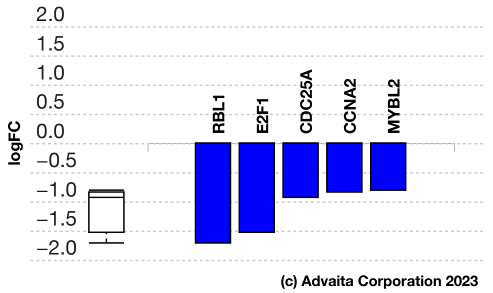


**Fig. 5.2.9: Gene measured expression bar plot:** *All the consistent differentially expressed genes that are targeted by E2F5 are ranked based on their absolute value of log fold change. Upregulated genes are shown in red, downregulated genes are shown in blue. The box and whisker plot on the left summarizes the distribution of all the consistent differentially expressed genes targeted by this upstream regulator. The box shows the 1st quartile, the median and the 3rd quartile, while the outliers are represented by circles.*

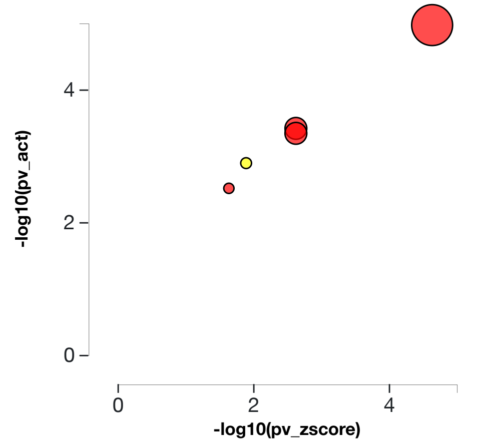


**Fig. 5.2.10: Activation p-value vs zscore p-value:** *E2F5, E2F transcription factor 5, (yellow) is shown, using negative log of the activation and zscore p-values, along with the other most significant upstream regulators. The size of the dot represents the relative number of consistent DE genes, which for selected upstream regulator is 5.*

#### RBL1 (RB transcriptional corepressor like 1)

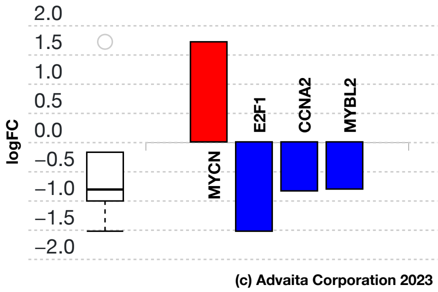


**Fig. 5.2.11: Gene measured expression bar plot:** *All the consistent differentially expressed genes that are targeted by RBL1 are ranked based on their absolute value of log fold change. Upregulated genes are shown in red, downregulated genes are shown in blue. The box and whisker plot on the left summarizes the distribution of all the consistent differentially expressed genes targeted by this upstream regulator. The box shows the 1st quartile, the median and the 3rd quartile, while the outliers are represented by circles.*

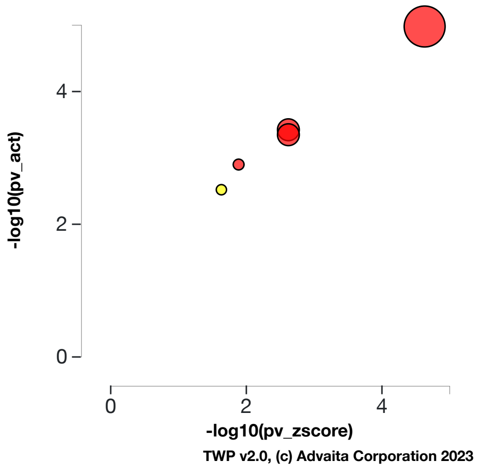


**Fig. 5.2.12: Activation p-value vs zscore p-value:** *RBL1, RB transcriptional corepressor like 1, (yellow) is shown, using negative log of the activation and zscore p-values, along with the other most significant upstream regulators. The size of the dot represents the relative number of consistent DE genes, which for selected upstream regulator is 4.*

### 5.3. Results: upstream regulators predicted as inhibited

| Upstream Regulator (u) | DTI(u) | DT(u) | p-value | p-value (FDR) | p-value (Bonferroni) |
|---|---|---|---|---|---|
| PPP2CB | 23 | 24 | **3.201e-8** | **5.240e-6** | **7.369e-5** |
| RANGAP1 | 23 | 24 | **5.198e-8** | **5.240e-6** | **1.197e-4** |
| BUB1 | 22 | 23 | **9.333e-8** | **5.240e-6** | **2.148e-4** |
| CENPC | 22 | 23 | **9.333e-8** | **5.240e-6** | **2.148e-4** |
| CENPE | 22 | 23 | **9.333e-8** | **5.240e-6** | **2.148e-4** |

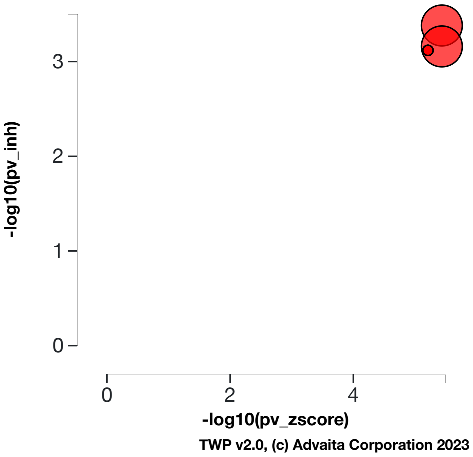


**Table 5.3.1: Top upstream regulators predicted as inhibited.** *For each upstream regulator u, the table shows the number of DE targets supporting the hypothesis that the regulator is inhibited DTI(u) the total number of DE genes downstream of u DT(u), the combined raw p-value, and the p-value corrected for multiple comparisons.* **Fig. 5.3.1: A two-way plot showing the top five upstream regulators predicted as inhibited.** *Dots representing upstream regulators are positioned using $P_{zscore}$ on the horizontal axis, and using $P_{inh}$ on the vertical axis. $P_{inh}$ is the p-value based on the number of DE targets consistent with the typeof the incoming signal and with the selected hypothesis type. Upstream regulators with a significant combined p-value are shown in red. The size of each dot represents the number of consistent DE genes for that regulator.*

#### PPP2CB (protein phosphatase 2 catalytic subunit beta)

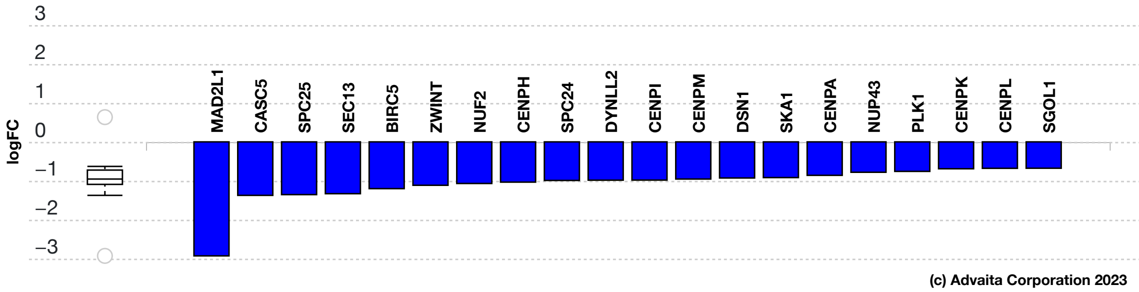


**Fig. 5.3.13: Gene measured expression bar plot:** *All the consistent differentially expressed genes that are targeted by PPP2CB are ranked based on their absolute value of log fold change. The plot is limited to the top 20 genes out of a total of 23 consistent differentially expressed target genes. Upregulated genes are shown in red, downregulated genes are shown in blue. The box and whisker plot on the left summarizes the distribution of all the consistent differentially expressed genes targeted by this upstream regulator. The box shows the 1st quartile, the median and the 3rd quartile, while the outliers are represented by circles.*

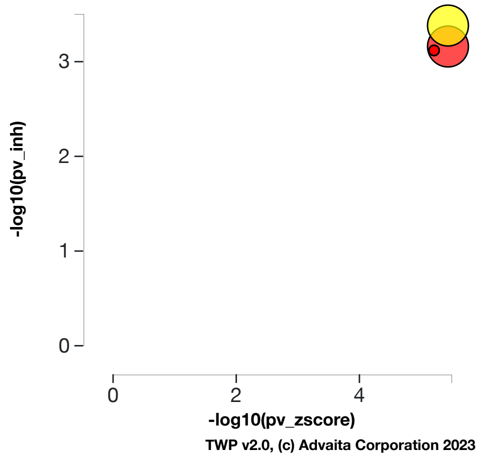


**Fig. 5.3.14: Inhibition p-value vs zscore p-value:** *PPP2CB, protein phosphatase 2 catalytic subunit beta, (yellow) is shown, using negative log of the inhibition and zscore p-values, along with the other most significant upstream regulators. The size of the dot represents the relative number of consistent DE genes, which for selected upstream regulator is 23.*

#### RANGAP1 (Ran GTPase activating protein 1)

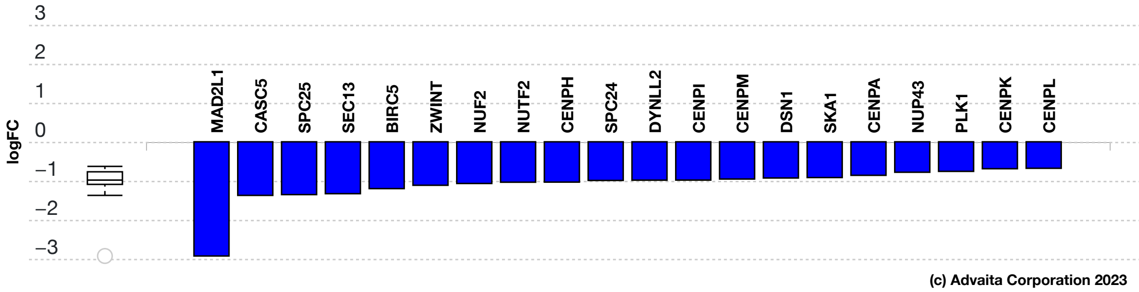


**Fig. 5.3.15: Gene measured expression bar plot:** *All the consistent differentially expressed genes that are targeted by RANGAP1 are ranked based on their absolute value of log fold change. The plot is limited to the top 20 genes out of a total of 23 consistent differentially expressed target genes. Upregulated genes are shown in red, downregulated genes are shown in blue. The box and whisker plot on the left summarizes the distribution of all the consistent differentially expressed genes targeted by this upstream regulator. The box shows the 1st quartile, the median and the 3rd quartile, while the outliers are represented by circles.*

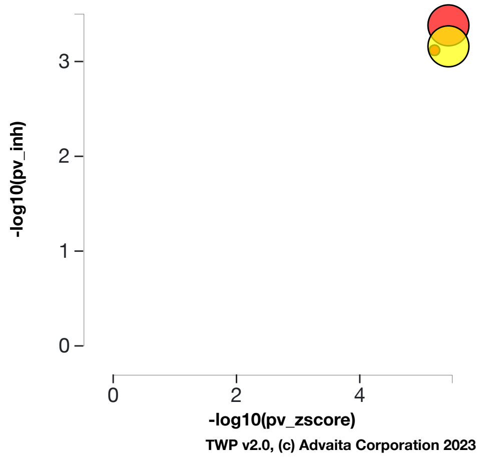


**Fig. 5.3.16: Inhibition p-value vs zscore p-value:** *RANGAP1, Ran GTPase activating protein 1, (yellow) is shown, using negative log of the inhibition and zscore p-values, along with the other most significant upstream regulators. The size of the dot represents the relative number of consistent DE genes, which for selected upstream regulator is 23.*

### BUB1 (BUB1 mitotic checkpoint serine/threonine kinase)

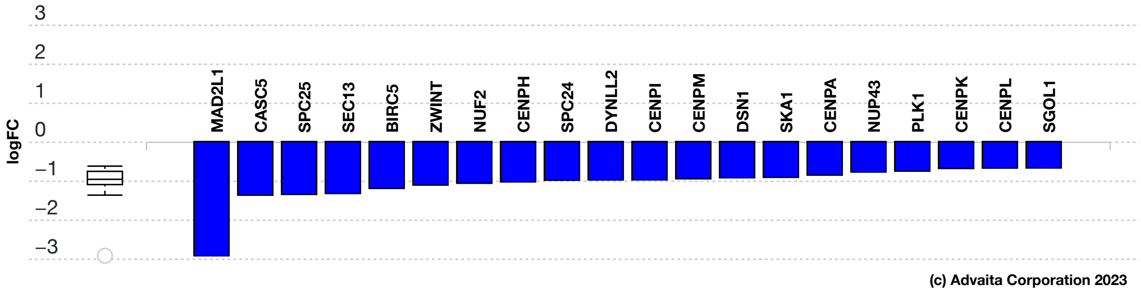


**Fig. 5.3.17: Gene measured expression bar plot:** *All the consistent differentially expressed genes that are targeted by BUB1 are ranked based on their absolute value of log fold change. The plot is limited to the top 20 genes out of a total of 22 consistent differentially expressed target genes. Upregulated genes are shown in red, downregulated genes are shown in blue. The box and whisker plot on the left summarizes the distribution of all the consistent differentially expressed genes targeted by this upstream regulator. The box shows the 1st quartile, the median and the 3rd quartile, while the outliers are represented by circles.*

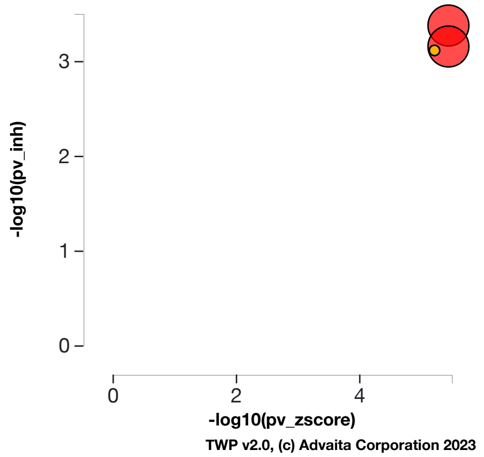


**Fig. 5.3.18: Inhibition p-value vs zscore p-value:** *BUB1, BUB1 mitotic checkpoint serine/threonine kinase, (yellow) is shown, using negative log of the inhibition and zscore p-values, along with the other most significant upstream regulators. The size of the dot represents the relative number of consistent DE genes, which for selected upstream regulator is 22.*

#### CENPC (centromere protein C)

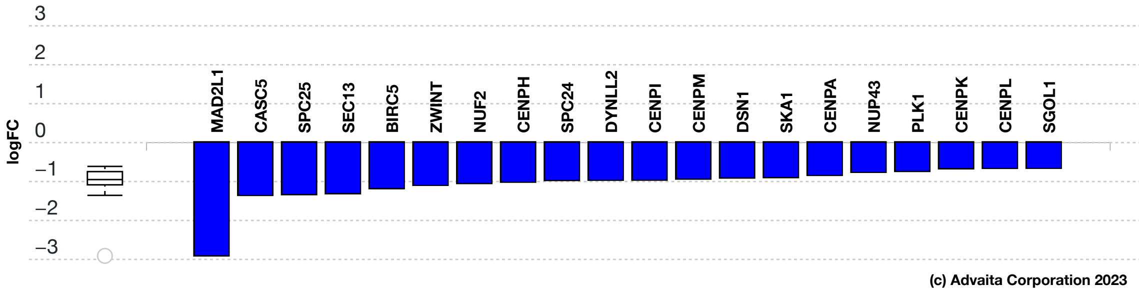


**Fig. 5.3.19: Gene measured expression bar plot:** *All the consistent differentially expressed genes that are targeted by CENPC are ranked based on their absolute value of log fold change. The plot is limited to the top 20 genes out of a total of 22 consistent differentially expressed target genes. Upregulated genes are shown in red, downregulated genes are shown in blue. The box and whisker plot on the left summarizes the distribution of all the consistent differentially expressed genes targeted by this upstream regulator. The box shows the 1st quartile, the median and the 3rd quartile, while the outliers are represented by circles.*

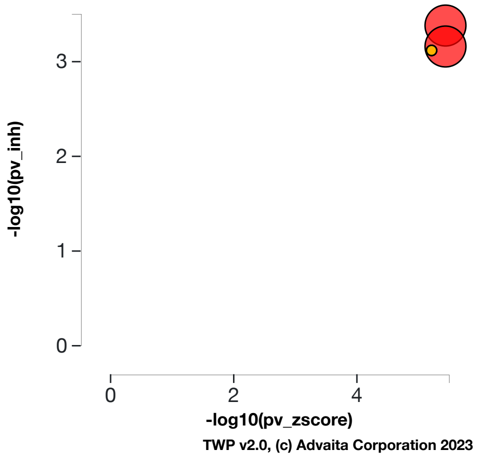


**Fig. 5.3.20: Inhibition p-value vs zscore p-value:** *CENPC, centromere protein C, (yellow) is shown, using negative log of the inhibition and zscore p-values, along with the other most significant upstream regulators. The size of the dot represents the relative number of consistent DE genes, which for selected upstream regulator is 22.*

#### CENPE (centromere protein E)

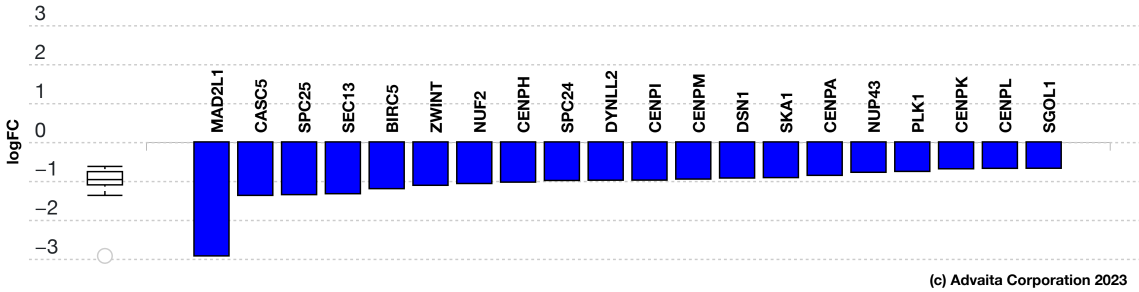


**Fig. 5.3.21: Gene measured expression bar plot:** *All the consistent differentially expressed genes that are targeted by CENPE are ranked based on their absolute value of log fold change. The plot is limited to the top 20 genes out of a total of 22 consistent differentially expressed target genes. Upregulated genes are shown in red, downregulated genes are shown in blue. The box and whisker plot on the left summarizes the distribution of all the consistent differentially expressed genes targeted by this upstream regulator. The box shows the 1st quartile, the median and the 3rd quartile, while the outliers are represented by circles.*

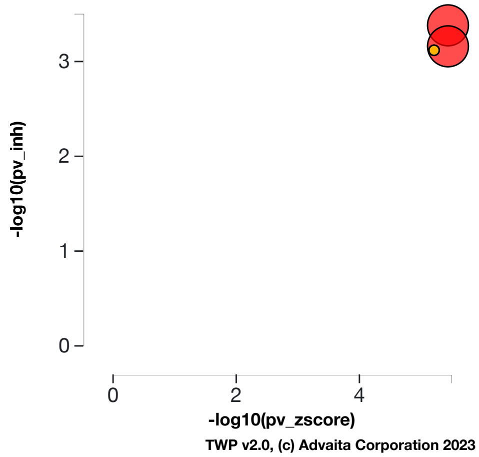


**Fig. 5.3.22: Inhibition p-value vs zscore p-value:** *CENPE, centromere protein E, (yellow) is shown, using negative log of the inhibition and zscore p-values, along with the other most significant upstream regulators. The size of the dot represents the relative number of consistent DE genes, which for selected upstream regulator is 22.*

### 6. Predicted Upstream Regulator Analysis – Chemicals, Drugs, Toxicants (CDTs)

#### 6.1. Methods

The prediction of upstream Chemicals, Drugs, Toxicants (CDTs) is based on two types of information: i) the enrichment of differentially expressed genes from the experiment and ii) a network of interactions from the Advaita Knowledge Base (AKB v17.0). The network is a directed graph in which the source node represents either a chemical substance or compound (e.g. zinc), a drug (e.g. aspirin), or a toxicant (e.g. tobacco smoke). The generic abbreviation CDT will be used henceforth to designate any of these. The edges represent known effects that these CDTs have on various genes. A signed edge in this graph consists of a source CDT, a target gene, and a sign to indicate the type of effect: activation (+) or inhibition (-). The analysis considers two hypotheses:

HP. The upstream chemical, drug or toxicant is **present (or overly abundant)** in the condition studied.
HA. The upstream chemical, drug or toxicant is **absent (or insufficient)** in the condition studied.

The analysis divides the set of all the genes from AKB into several subsets based on the measurements in the experiment and the definitions shown in **Figure 6.1.1** and **Figure 6.1.2**. Let the sign of a measured DE gene be the sign of the log fold change value: (+) for up-regulated genes and (-) for down-regulated genes. A gene is a target gene if it corresponds to a node in the network that has at least one incoming edge. We define a *consistent gene* as a target DE gene such that the sign of the gene is consistent both with the type of the signal **and** with the hypothesis considered. Formally, by definition, a target DE gene *g* is consistent with Hypothesis HP if and only if an incoming edge *e* exists such that *sign(g) = sign(e)*. In other words, this describes the situation when the CDT is predicted as present, the signal is activation and the target DE gene is up-regulated, or the signal is inhibition and the target DE gene is down-regulated (see panel A in **Figure 6.1.1**). A target DE gene *g* is consistent with Hypothesis HA if and only if an incoming edge *e* exists such that *sign(g) ≠ sign(e)*. This second case captures the situation in which the CDT is absent (or insufficient), the signal is inhibition and the target DE gene is up-regulated, or the signal is activation and the target DE gene is down-regulated (see panel B in **Figure 6.1.1**).

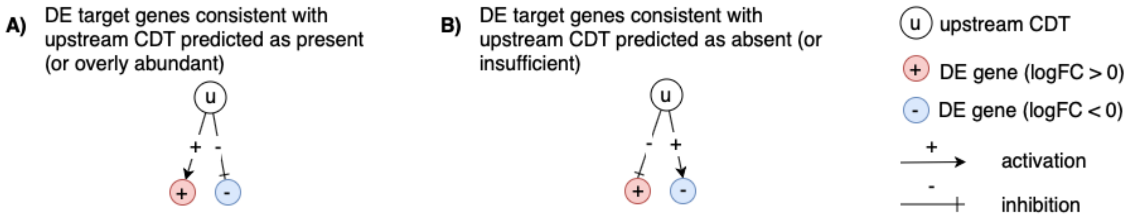


**Fig. 6.1.1: Target genes consistent with the hypothesis considered:** *In panel A, the signs of the DE genes match the signs of their respective incoming edges, increasing the likelihood that the CDT u is present. In panel B, the signs of the DE genes are opposite to the signs of their edges, increasing the likelihood that the CDT u is absent.*

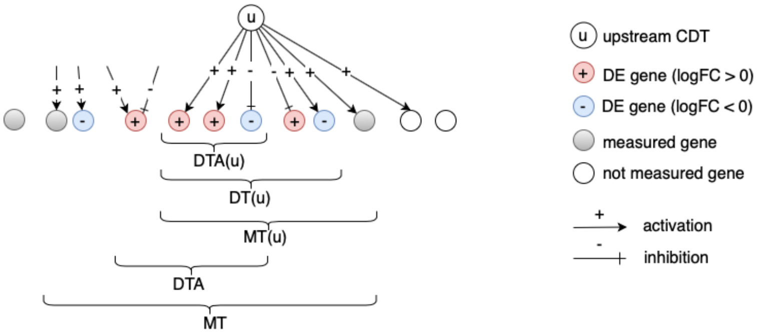


**Fig. 6.1.2:** *The set of all genes includes the set of measured genes that are also targets in the network, or Measured Targets (MT). We define the subset of "DE Targets consistent with the first hypothesis that the CDTs are Present (or overly abundant)", DTA. For a selected upstream CDT u, we have the set of "Measured Targets of u" MT(u), "Differentially expressed Targets downstream of u" DT(u), and the set of "DE targets consistent with the hypothesis HP that u is Present" DTA(u). The equivalent graphic for the hypothesis HA associated with DTI and DTI(u) is not shown.*

#### Z-score

For both research hypotheses, the analysis computes a Z-score for each CDT *z(u)* by iterating over the genes in DT(u) and their incoming edges *in(g)*. We can then compute the p-value corresponding to the z-score $P_z$ as the one-tailed area under the probability density function for a normal distribution, N(0,1).

#### Upstream CDTs predicted as present (or overly abundant)

Here, the research hypothesis considers presence of the CDT. This hypothesis is useful when investigating whether the given phenotype has been impacted by the presence of a given chemical, drug or toxicant (e.g. tobacco smoke, dioxin, etc.). For each CDT *u*, the number of consistent DE genes downstream of *u*, DTA(u) is compared to the number of measured target genes expected to be both consistent and DE just by chance. iPathwayGuide uses an over-representation approach to compute the statistical significance of observing at least the given number of consistent DE genes. The p-value $P_{pres}$ is computed using the hypergeometric distribution (Draghici *et al*., 2003, Draghici 2011).

After computing a p-value for both types of evidence, $P_z$ and $P_{pres}$, we combine these two probabilities into one global probability value, $P_G$ that is used to rank the upstream regulators and test the research hypothesis that the upstream CDTs are predicted as present in the condition studied. The analysis uses the standard Fisher's method to combine p-values into one test statistic (Fisher 1925).

#### Upstream CDTs predicted as absent (or insufficient)

In parallel with upstream CDTs predicted as present, we use $P_{abs}$ and $P_z$ to predict upstream CDTs that are absent. This hypothesis is relevant when investigating whether the given phenotype has been impacted by the lack of a given chemical that is necessary for the well-functioning of the organism or cell (e.g. a vitamin deficiency, iron deficiency, etc.). Here, the research hypothesis states that the upstream CDT are insufficient in the condition studied. For each upstream CDT *u*, the number of consistent DE genes downstream of *u*, DTI(u) is compared to the number of measured target genes expected to be both consistent and DE just by chance. Using the Fisher's method as above, the analysis combines $P_{abs}$ and $P_z$, where $P_z$ is considered only for significant negative z-scores ( $z \leq -2$).

### 6.2. Results: upstream CDTs predicted as present (or overly abundant)

| CDT (u) | DTA(u) | DT(u) | p-value | p-value (FDR) | p-value (Bonferroni) |
|---|---|---|---|---|---|
| Testosterone | 300 | 358 | **5.626e-23** | **3.625e-20** | **1.450e-19** |
| Dasatinib | 146 | 166 | **5.626e-23** | **3.625e-20** | **1.450e-19** |
| Lucanthone | 82 | 85 | **5.626e-23** | **3.625e-20** | **1.450e-19** |
| palbociclib | 66 | 67 | **5.626e-23** | **3.625e-20** | **1.450e-19** |
| incobotulinumtoxinA | 173 | 237 | **7.119e-22** | **3.659e-19** | **1.835e-18** |

**Table 6.2.1: Top upstream CDTs predicted as present (or overly abundant).** *For each upstream CDT u, the table shows the number of DE targets supporting the hypothesis that the CDT is present DTA(u) the total number of DE genes downstream of u DT(u), the combined raw p-value, and the p-value corrected for multiple comparisons.*

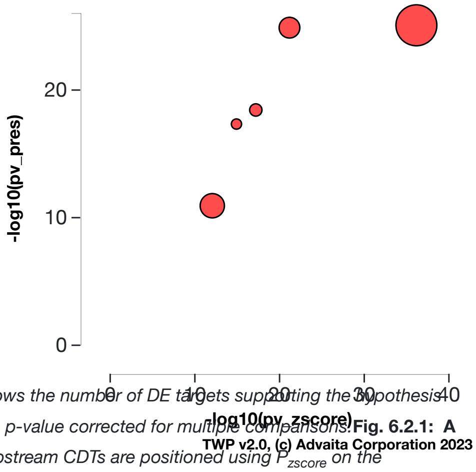


**Fig. 6.2.1: A two-way plot showing the top five upstream CDTs predicted as present (or overly abundant).** *Dots representing upstream CDTs are positioned using $P_{zscore}$ on the horizontal axis, and using $P_{pres}$ on the vertical axis. $P_{pres}$ is the p-value based on the number of DE targets consistent with the typeof the incoming signal and with the selected hypothesis type. Upstream CDTs with a significant combined p-value are shown in red. The size of each dot represents the relative number of consistent DE genes for that CDT.*

#### Testosterone

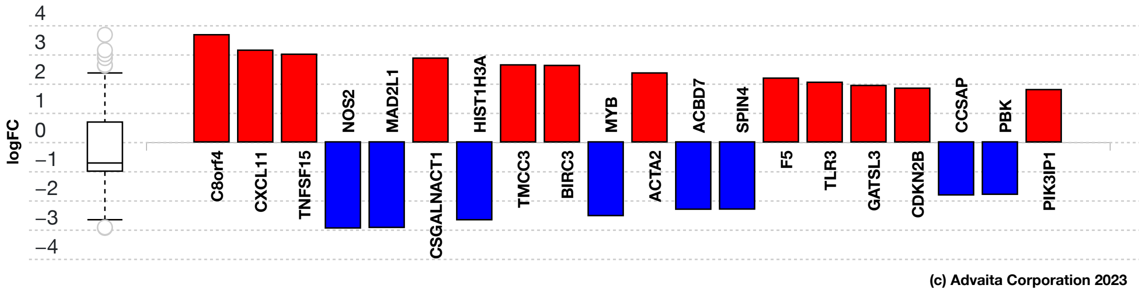


**Fig. 6.2.3: Consistent DE target genes measured expression bar plot:** *All the consistent differentially expressed genes that are targeted by Testosterone are ranked based on their absolute value of log fold change. The plot is limited to the top 20 genes out of a total of 300 consistent differentially expressed target genes. Upregulated genes are shown in red, downregulated genes are shown in blue. The box and whisker plot on the left summarizes the distribution of all the consistent differentially expressed genes targeted by this upstream regulator. The box shows the 1st quartile, the median and the 3rd quartile, while any outliers are represented by circles.*

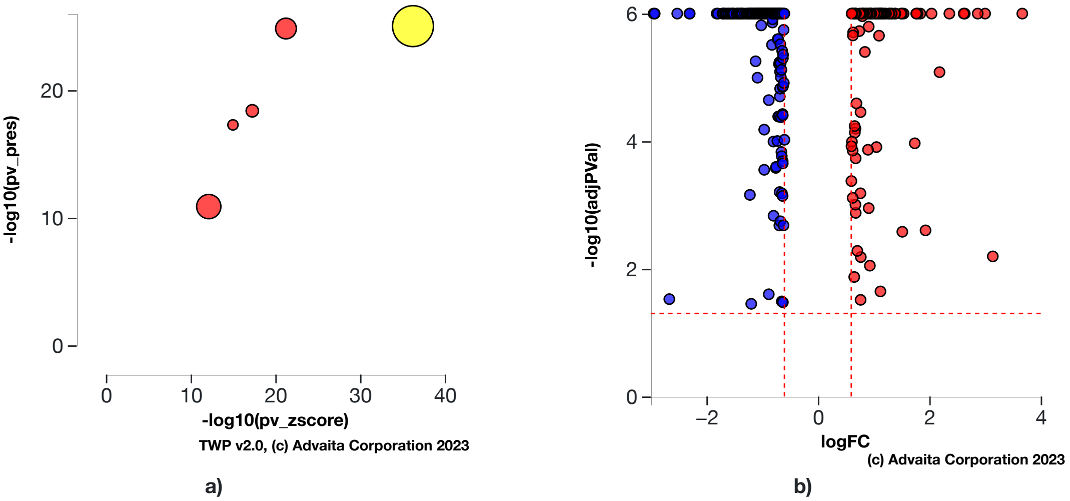


**Fig. 6.2.4: a) Present (overly abundant) p-value vs zscore p-value:** *The significance of Testosterone is plotted on two axes, with negative log of $P_z$ on x-axis and negative log of $P_{pres}$ on y-axis. The size of the dot represents the relative number of consistent DE genes, which for selected upstream regulator is 300.* **b) Volcano plot:** *There are* ***300*** *DE genes that are targets of Testosterone consistent with the hypothesis that Testosterone is present (overly abundant) The target genes are represented in terms of their measured expression change (x-axis) and the significance of the change (y-axis). The significance is represented in terms of the negative log (base 10) of the p-value, so that more significant genes are plotted higher on the y-axis. The dotted lines represent the thresholds used to select the DE genes:* ***0.6*** *for expression change and* ***0.05*** *for significance.*

#### Dasatinib

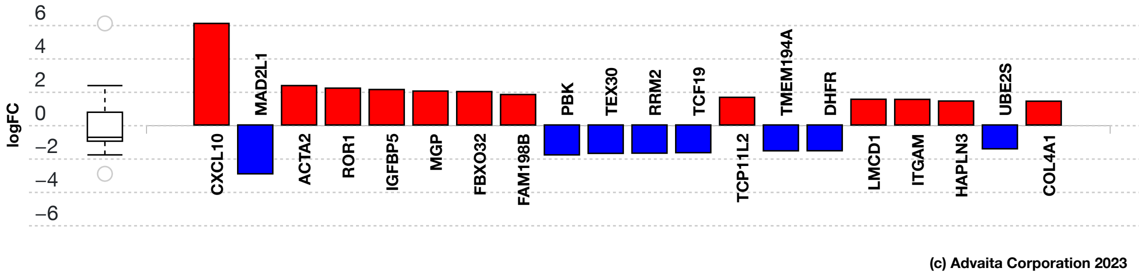


**Fig. 6.2.5: Consistent DE target genes measured expression bar plot:** *All the consistent differentially expressed genes that are targeted by Dasatinib are ranked based on their absolute value of log fold change. The plot is limited to the top 20 genes out of a total of 146 consistent differentially expressed target genes. Upregulated genes are shown in red, downregulated genes are shown in blue. The box and whisker plot on the left summarizes the distribution of all the consistent differentially expressed genes targeted by this upstream regulator. The box shows the 1st quartile, the median and the 3rd quartile, while any outliers are represented by circles.*

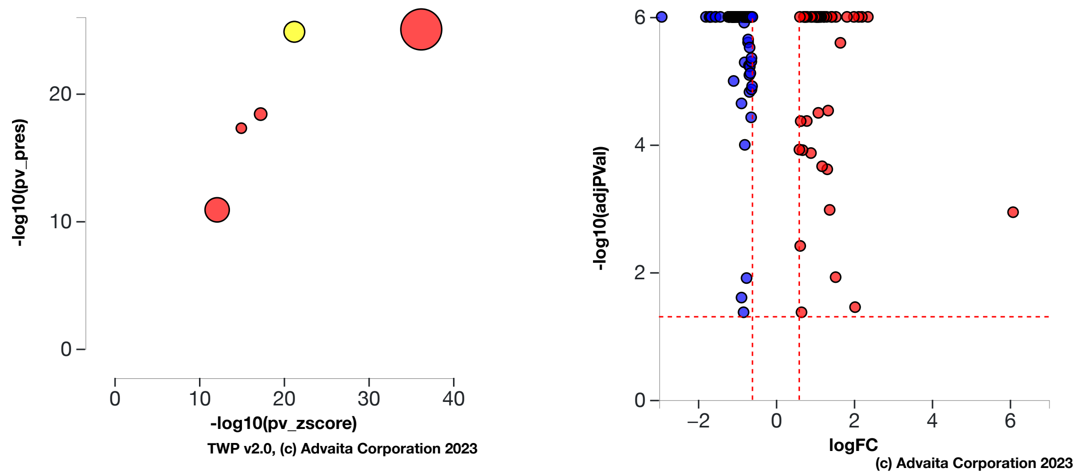


a) b)

**Fig. 6.2.6: a) Present (overly abundant) p-value vs zscore p-value:** *The significance of Dasatinib is plotted on two axes, with negative log of $P_z$ on x-axis and negative log of $P_{pres}$ on y-axis. The size of the dot represents the relative number of consistent DE genes, which for selected upstream regulator is 146.* **b) Volcano plot:** *There are* ***146*** *DE genes that are targets of Dasatinib consistent with the hypothesis that Dasatinib is present (overly abundant) The target genes are represented in terms of their measured expression change (x-axis) and the significance of the change (y-axis). The significance is represented in terms of the negative log (base 10) of the p-value, so that more significant genes are plotted higher on the y-axis. The dotted lines represent the thresholds used to select the DE genes:* ***0.6*** *for expression change and* ***0.05*** *for significance.*

#### Lucanthone

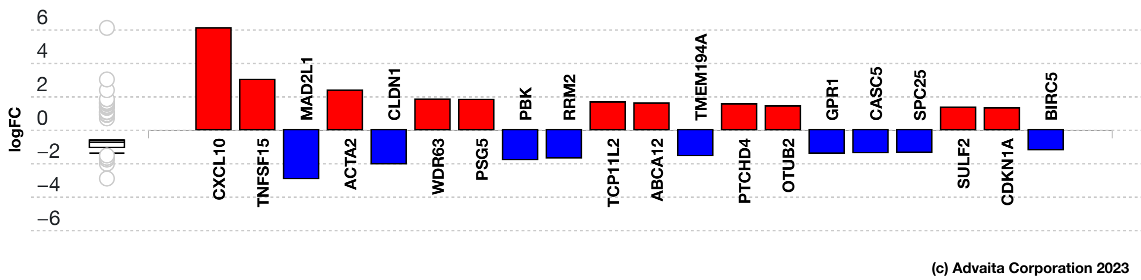


**Fig. 6.2.7: Consistent DE target genes measured expression bar plot:** *All the consistent differentially expressed genes that are targeted by Lucanthone are ranked based on their absolute value of log fold change. The plot is limited to the top 20 genes out of a total of 82 consistent differentially expressed target genes. Upregulated genes are shown in red, downregulated genes are shown in blue. The box and whisker plot on the left summarizes the distribution of all the consistent differentially expressed genes targeted by this upstream regulator. The box shows the 1st quartile, the median and the 3rd quartile, while any outliers are represented by circles.*

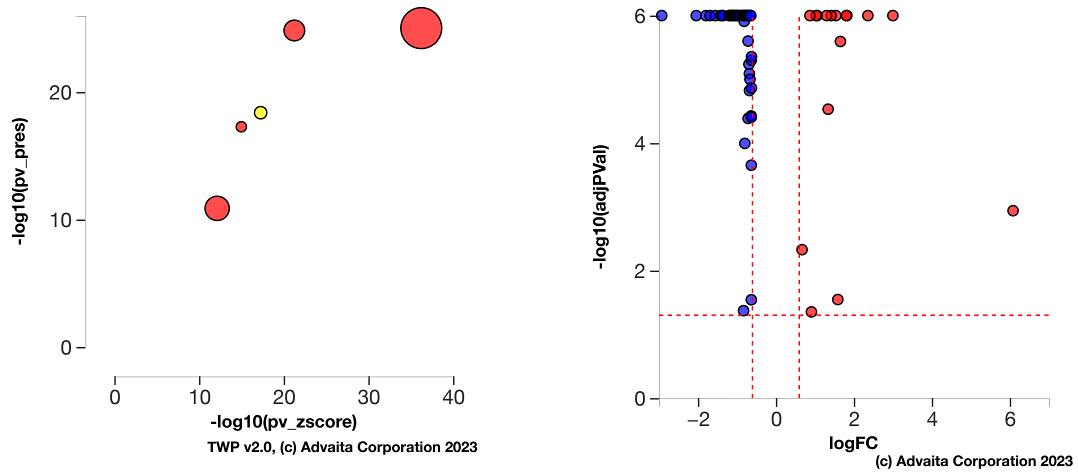


a) b)

**Fig. 6.2.8: a) Present (overly abundant) p-value vs zscore p-value:** *The significance of Lucanthone is plotted on two axes, with negative log of $P_z$ on x-axis and negative log of $P_{pres}$ on y-axis. The size of the dot represents the relative number of consistent DE genes, which for selected upstream regulator is 82.* **b) Volcano plot:** *There are* ***82*** *DE genes that are targets of Lucanthone consistent with the hypothesis that Lucanthone is present (overly abundant) The target genes are represented in terms of their measured expression change (x-axis) and the significance of the change (y-axis). The significance is represented in terms of the negative log (base 10) of the p-value, so that more significant genes are plotted higher on the y-axis. The dotted lines represent the thresholds used to select the DE genes:* ***0.6*** *for expression change and* ***0.05*** *for significance.*

#### palbociclib

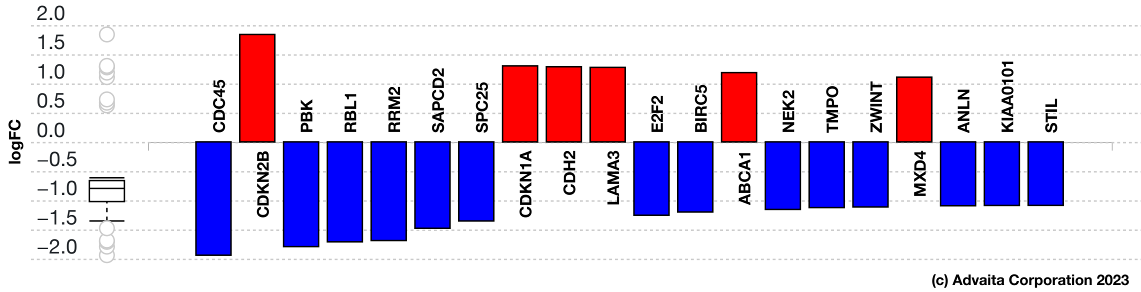


**Fig. 6.2.9: Consistent DE target genes measured expression bar plot:** *All the consistent differentially expressed genes that are targeted by palbociclib are ranked based on their absolute value of log fold change. The plot is limited to the top 20 genes out of a total of 66 consistent differentially expressed target genes. Upregulated genes are shown in red, downregulated genes are shown in blue. The box and whisker plot on the left summarizes the distribution of all the consistent differentially expressed genes targeted by this upstream regulator. The box shows the 1st quartile, the median and the 3rd quartile, while any outliers are represented by circles.*

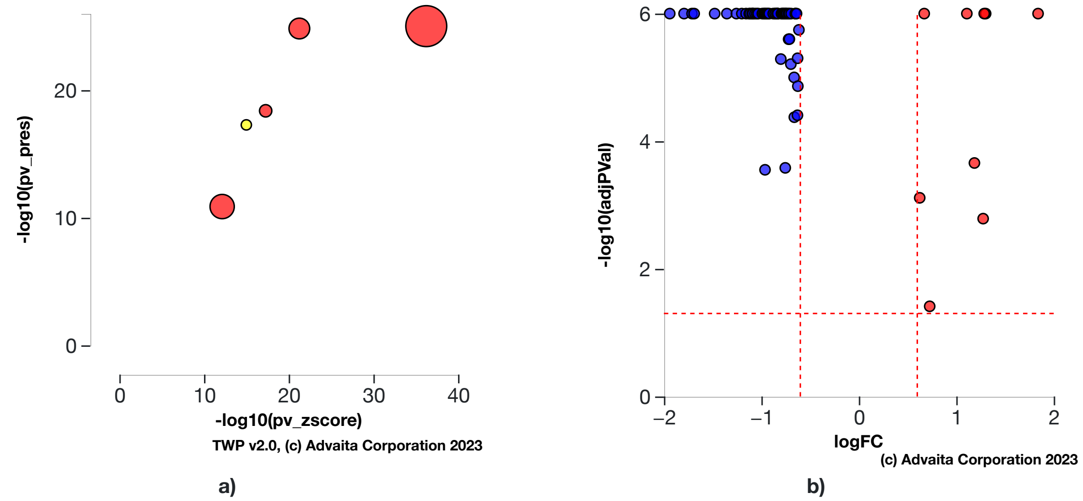


**Fig. 6.2.10: a) Present (overly abundant) p-value vs zscore p-value:** *The significance of palbociclib is plotted on two axes, with negative log of $P_z$ on x-axis and negative log of $P_{pres}$ on y-axis. The size of the dot represents the relative number of consistent DE genes, which for selected upstream regulator is 66.* **b) Volcano plot:** *There are **66** DE genes that are targets of palbociclib consistent with the hypothesis that palbociclib is present (overly abundant) The target genes are represented in terms of their measured expression change (x-axis) and the significance of the change (y-axis). The significance is represented in terms of the negative log (base 10) of the p-value, so that more significant genes are plotted higher on the y-axis. The dotted lines represent the thresholds used to select the DE genes: **0.6** for expression change and **0.05** for significance.*

#### incobotulinumtoxinA

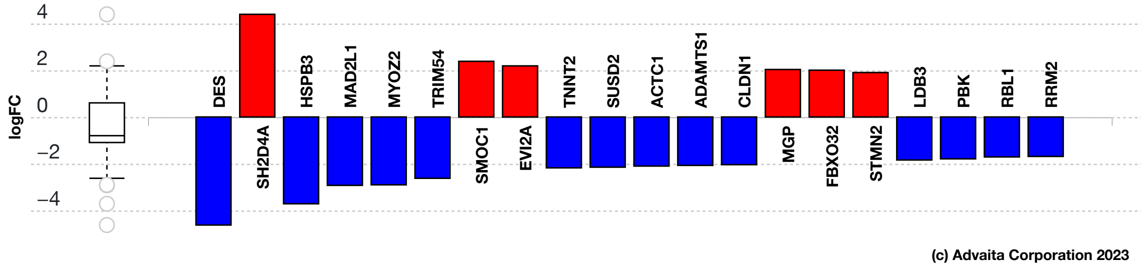


**Fig. 6.2.11: Consistent DE target genes measured expression bar plot:** *All the consistent differentially expressed genes that are targeted by incobotulinumtoxinA are ranked based on their absolute value of log fold change. The plot is limited to the top 20 genes out of a total of 173 consistent differentially expressed target genes. Upregulated genes are shown in red, downregulated genes are shown in blue. The box and whisker plot on the left summarizes the distribution of all the consistent differentially expressed genes targeted by this upstream regulator. The box shows the 1st quartile, the median and the 3rd quartile, while any outliers are represented by circles.*

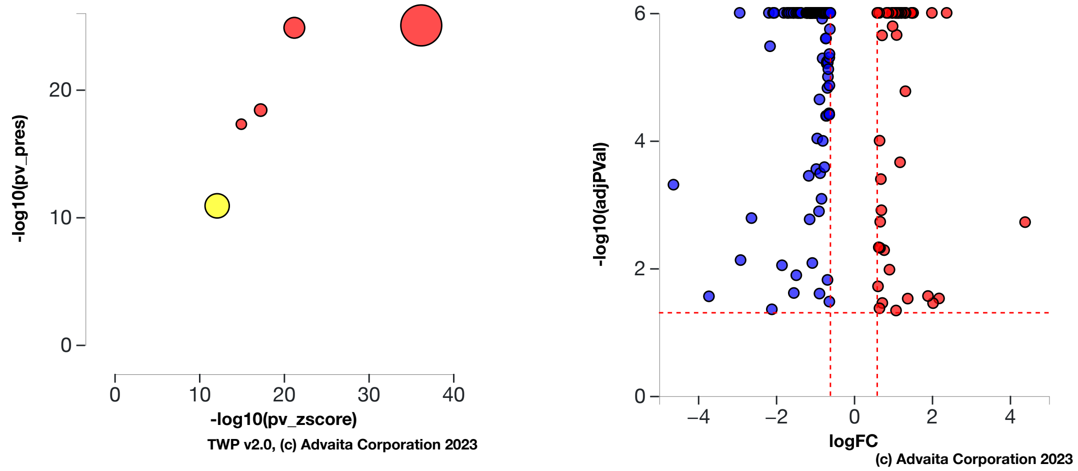


a) b)

**Fig. 6.2.12: a) Present (overly abundant) p-value vs zscore p-value:** *The significance of incobotulinumtoxinA is plotted on two axes, with negative log of $P_z$ on x-axis and negative log of $P_{pres}$ on y-axis. The size of the dot represents the relative number of consistent DE genes, which for selected upstream regulator is 173.* **b) Volcano plot:** *There are **173** DE genes that are targets of incobotulinumtoxinA consistent with the hypothesis that incobotulinumtoxinA is present (overly abundant) The target genes are represented in terms of their measured expression change (x-axis) and the significance of the change (y-axis). The significance is represented in terms of the negative log (base 10) of the p-value, so that more significant genes are plotted higher on the y-axis. The dotted lines represent the thresholds used to select the DE genes: **0.6** for expression change and **0.05** for significance.*

### 6.3. Results: upstream CDTs predicted as absent (or insufficient)

| CDT (u) | DTI(u) | DT(u) | p-value | p-value (FDR) | p-value (Bonferroni) |
|---|---|---|---|---|---|
| Coumestrol | 294 | 365 | **9.438e-22** | **2.432e-18** | **2.432e-18** |
| Polychlorinated Biphenyls | 64 | 74 | **3.842e-19** | **4.951e-16** | **9.902e-16** |
| Phytoestrogens | 30 | 31 | **4.263e-18** | **3.662e-15** | **1.099e-14** |
| Latex | 50 | 59 | **3.046e-13** | **1.962e-10** | **7.850e-10** |
| 2-methoxy-N-(3-methyl-2-oxo-1,2,3,4-tetrahydroquinazolin-6-yl)benzenesulfonamide | 15 | 16 | **7.882e-9** | **4.062e-6** | **2.031e-5** |

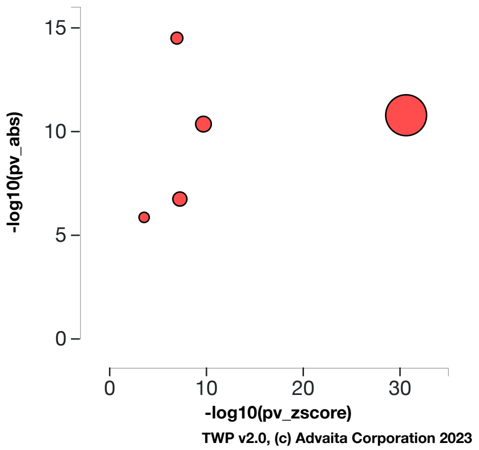


**Table 6.3.1: Top upstream CDTs predicted as absent (or insufficient).** *For each upstream CDT u, the table shows the number of DE targets supporting the hypothesis that the CDT is absent DTI(u) the total number of DE genes downstream of u DT(u), the combined raw p-value, and the p-value corrected for multiple comparisons.* **Fig. 6.3.1: A two-way plot showing the top five upstream CDTs predicted as absent (or insufficient).** *Dots representing upstream CDTs are positioned using $P_{zscore}$ on the horizontal axis, and using $P_{abs}$ on the vertical axis. $P_{abs}$ is the p-value based on the number of DE targets consistent with the typeof the incoming signal and with the selected hypothesis type. Upstream CDTs with a significant combined p-value are shown in red. The size of each dot represents the relative number of consistent DE genes for that CDT.*

#### Coumestrol

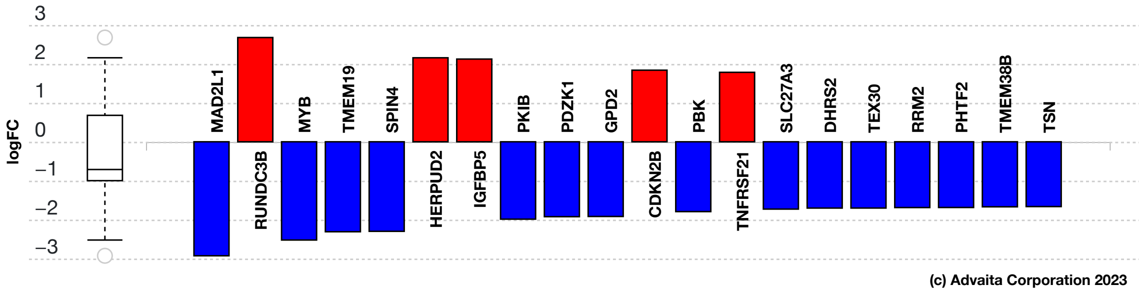


**Fig. 6.3.13: Consistent DE target genes measured expression bar plot:** *All the consistent differentially expressed genes that are targeted by Coumestrol are ranked based on their absolute value of log fold change. The plot is limited to the top 20 genes out of a total of 294 consistent differentially expressed target genes. Upregulated genes are shown in red, downregulated genes are shown in blue. The box and whisker plot on the left summarizes the distribution of all the consistent differentially expressed genes targeted by this upstream regulator. The box shows the 1st quartile, the median and the 3rd quartile, while any outliers are represented by circles.*

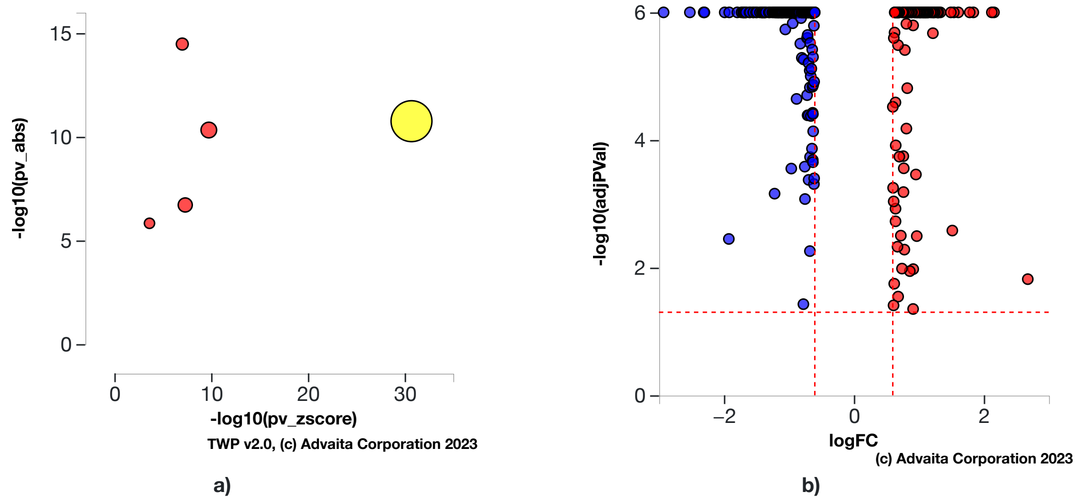


**Fig. 6.3.14: a) Absent (or insufficient) p-value vs zscore p-value:** *The significance of Coumestrol is plotted on two axes, with negative log of $P_z$ on x-axis and negative log of $P_{abs}$ on y-axis. The size of the dot represents the relative number of consistent DE genes, which for selected upstream regulator is 294.* **b) Volcano plot:** *There are **294** DE genes that are targets of Coumestrol consistent with the hypothesis that Coumestrol is absent (or insufficient) The target genes are represented in terms of their measured expression change (x-axis) and the significance of the change (y-axis). The significance is represented in terms of the negative log (base 10) of the p-value, so that more significant genes are plotted higher on the y-axis. The dotted lines represent the thresholds used to select the DE genes: **0.6** for expression change and **0.05** for significance.*

#### Polychlorinated Biphenyls

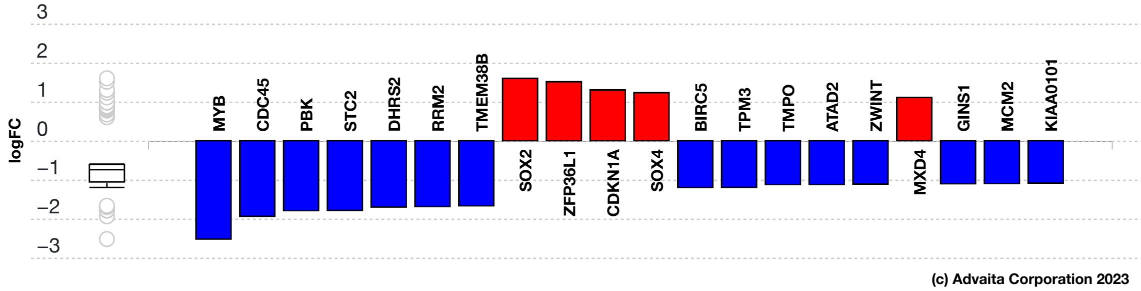


**Fig. 6.3.15: Consistent DE target genes measured expression bar plot:** *All the consistent differentially expressed genes that are targeted by Polychlorinated Biphenyls are ranked based on their absolute value of log fold change. The plot is limited to the top 20 genes out of a total of 64 consistent differentially expressed target genes. Upregulated genes are shown in red, downregulated genes are shown in blue. The box and whisker plot on the left summarizes the distribution of all the consistent differentially expressed genes targeted by this upstream regulator. The box shows the 1st quartile, the median and the 3rd quartile, while any outliers are represented by circles.*

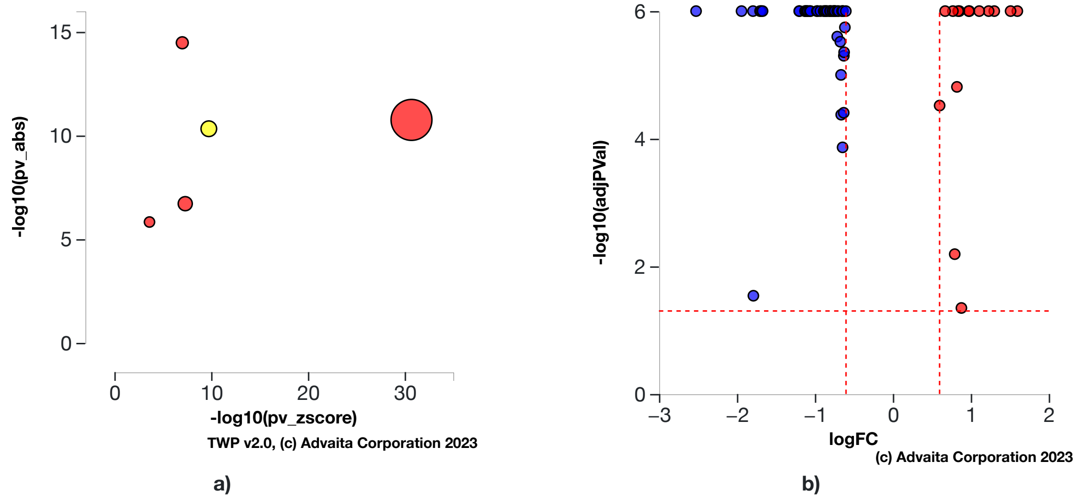


**Fig. 6.3.16: a) Absent (or insufficient) p-value vs zscore p-value:** *The significance of Polychlorinated Biphenyls is plotted on two axes, with negative log of $P_z$ on x-axis and negative log of $P_{abs}$ on y-axis. The size of the dot represents the relative number of consistent DE genes, which for selected upstream regulator is 64.* **b) Volcano plot:** *There are **64** DE genes that are targets of Polychlorinated Biphenyls consistent with the hypothesis that Polychlorinated Biphenyls is absent (or insufficient) The target genes are represented in terms of their measured expression change (x-axis) and the significance of the change (y-axis). The significance is represented in terms of the negative log (base 10) of the p-value, so that more significant genes are plotted higher on the y-axis. The dotted lines represent the thresholds used to select the DE genes: **0.6** for expression change and **0.05** for significance.*

#### Phytoestrogens

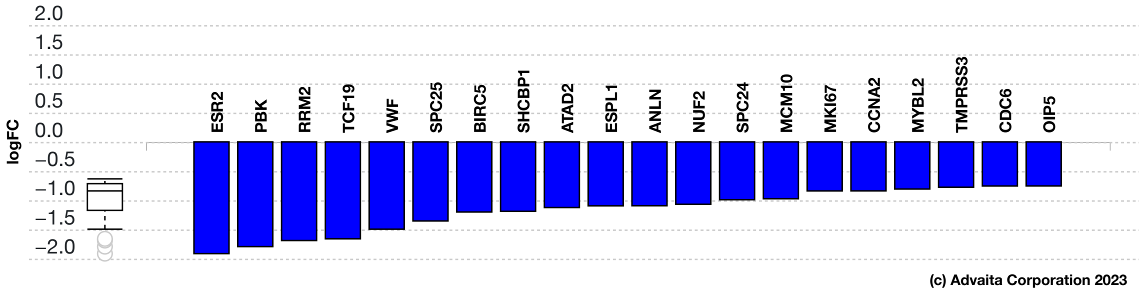


**Fig. 6.3.17: Consistent DE target genes measured expression bar plot:** *All the consistent differentially expressed genes that are targeted by Phytoestrogens are ranked based on their absolute value of log fold change. The plot is limited to the top 20 genes out of a total of 30 consistent differentially expressed target genes. Upregulated genes are shown in red, downregulated genes are shown in blue. The box and whisker plot on the left summarizes the distribution of all the consistent differentially expressed genes targeted by this upstream regulator. The box shows the 1st quartile, the median and the 3rd quartile, while any outliers are represented by circles.*

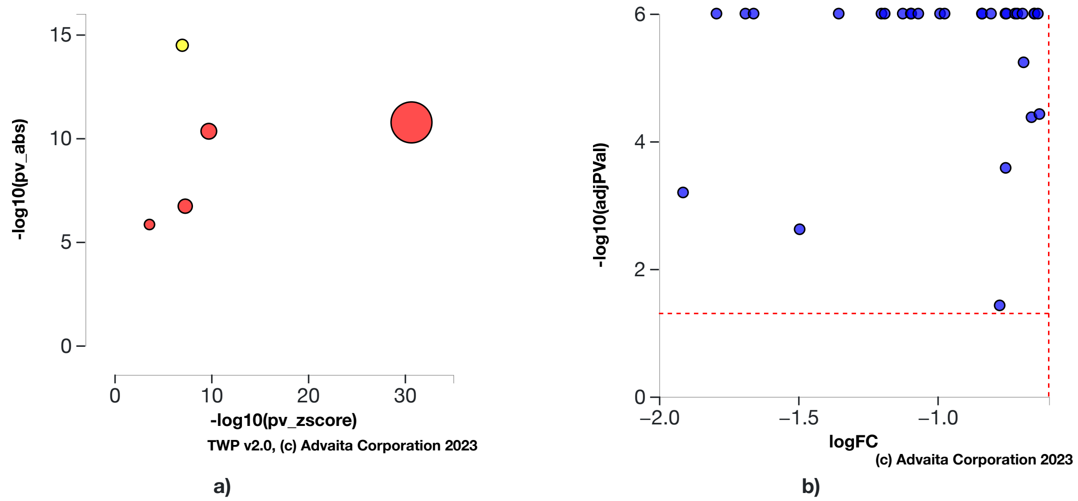


**Fig. 6.3.18: a) Absent (or insufficient) p-value vs zscore p-value:** *The significance of Phytoestrogens is plotted on two axes, with negative log of $P_z$ on x-axis and negative log of $P_{abs}$ on y-axis. The size of the dot represents the relative number of consistent DE genes, which for selected upstream regulator is 30.* **b) Volcano plot:** *There are **30** DE genes that are targets of Phytoestrogens consistent with the hypothesis that Phytoestrogens is absent (or insufficient) The target genes are represented in terms of their measured expression change (x-axis) and the significance of the change (y-axis). The significance is represented in terms of the negative log (base 10) of the p-value, so that more significant genes are plotted higher on the y-axis. The dotted lines represent the thresholds used to select the DE genes: **0.6** for expression change and **0.05** for significance.*

#### Latex

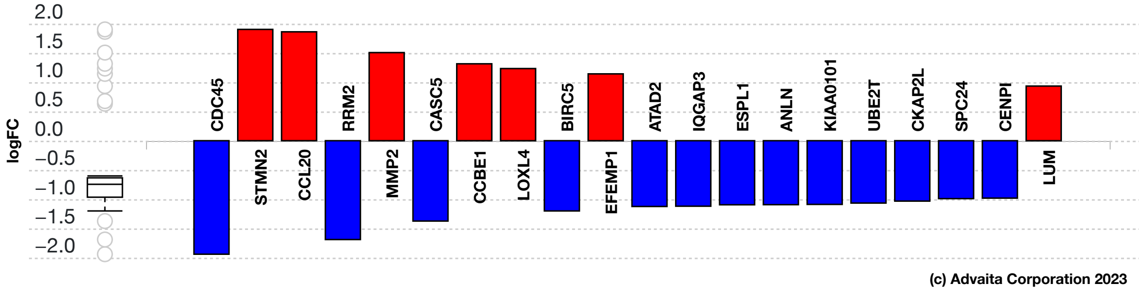


**Fig. 6.3.19: Consistent DE target genes measured expression bar plot:** *All the consistent differentially expressed genes that are targeted by Latex are ranked based on their absolute value of log fold change. The plot is limited to the top 20 genes out of a total of 50 consistent differentially expressed target genes. Upregulated genes are shown in red, downregulated genes are shown in blue. The box and whisker plot on the left summarizes the distribution of all the consistent differentially expressed genes targeted by this upstream regulator. The box shows the 1st quartile, the median and the 3rd quartile, while any outliers are represented by circles.*

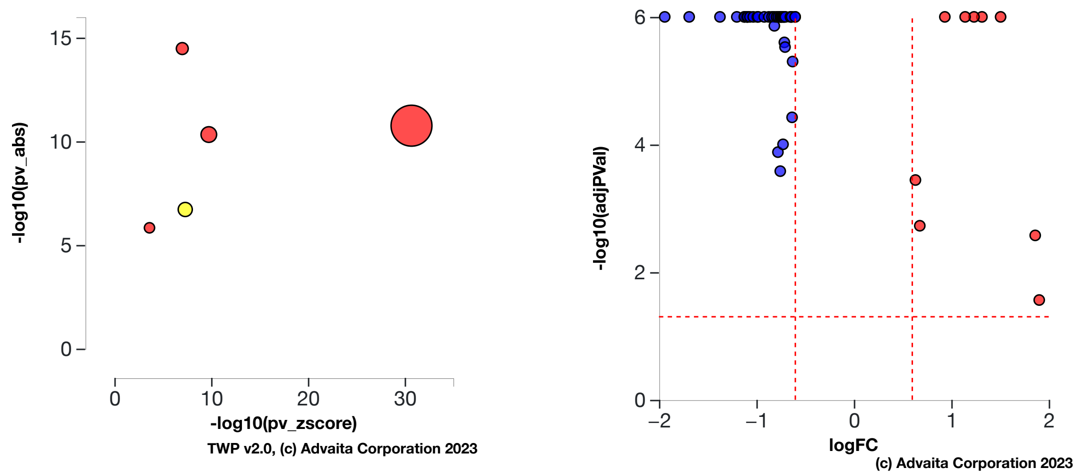


a) b)

**Fig. 6.3.20: a) Absent (or insufficient) p-value vs zscore p-value:** *The significance of Latex is plotted on two axes, with negative log of $P_z$ on x-axis and negative log of $P_{abs}$ on y-axis. The size of the dot represents the relative number of consistent DE genes, which for selected upstream regulator is 50.* **b) Volcano plot:** *There are* ***50*** *DE genes that are targets of Latex consistent with the hypothesis that Latex is absent (or insufficient) The target genes are represented in terms of their measured expression change (x-axis) and the significance of the change (y-axis). The significance is represented in terms of the negative log (base 10) of the p-value, so that more significant genes are plotted higher on the y-axis. The dotted lines represent the thresholds used to select the DE genes:* ***0.6*** *for expression change and* ***0.05*** *for significance.*

#### 2-methoxy-N-(3-methyl-2-oxo-1,2,3,4-tetrahydroquinazolin-6-yl)benzenesulfonamide

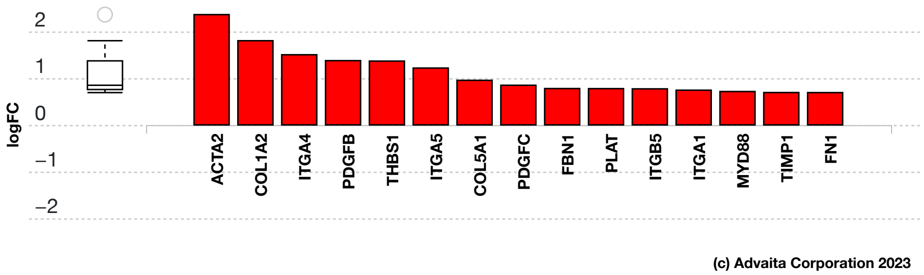


**Fig. 6.3.21: Consistent DE target genes measured expression bar plot:** *All the consistent differentially expressed genes that are targeted by 2-methoxy-N-(3-methyl-2-oxo-1,2,3,4-tetrahydroquinazolin-6-yl)benzenesulfonamide are ranked based on their absolute value of log fold change. The box and whisker plot on the left summarizes the distribution of all the consistent differentially expressed genes targeted by this upstream regulator. The box shows the 1st quartile, the median and the 3rd quartile, while any outliers are represented by circles.*

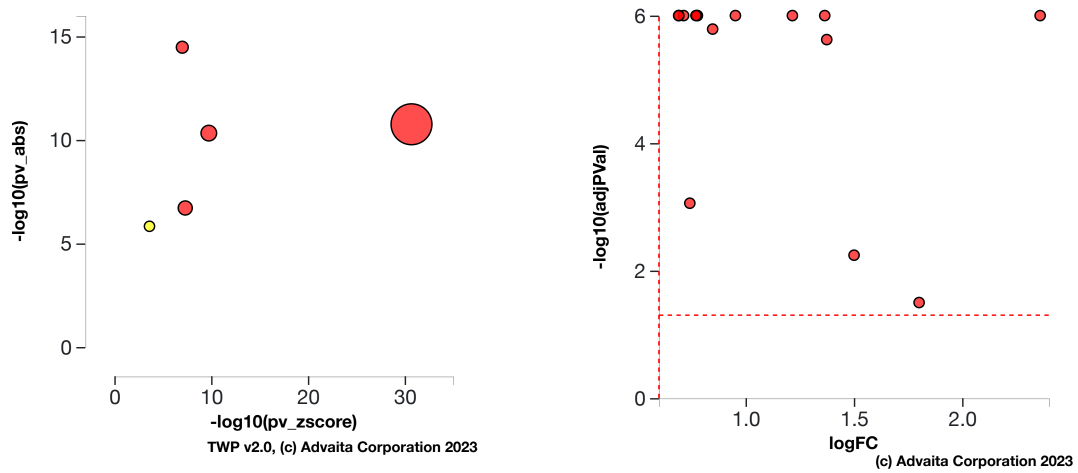


a) b)

**Fig. 6.3.22: a) Absent (or insufficient) p-value vs zscore p-value:** *The significance of 2-methoxy-N-(3-methyl-2-oxo-1,2,3,4-tetrahydroquinazolin-6-yl)benzenesulfonamide is plotted on two axes, with negative log of $P_z$ on x-axis and negative log of $P_{abs}$ on y-axis. The size of the dot represents the relative number of consistent DE genes, which for selected upstream regulator is 15.* **b) Volcano plot:** *There are* ***15*** *DE genes that are targets of 2-methoxy-N-(3-methyl-2-oxo-1,2,3,4-tetrahydroquinazolin-6-yl)benzenesulfonamide consistent with the hypothesis that 2-methoxy-N-(3-methyl-2-oxo-1,2,3,4-tetrahydroquinazolin-6-yl)benzenesulfonamide is absent (or insufficient) The target genes are represented in terms of their measured expression change (x-axis) and the significance of the change (y-axis). The significance is represented in terms of the negative log (base 10) of the p-value, so that more significant genes are plotted higher on the y-axis. The dotted lines represent the thresholds used to select the DE genes:* ***0.6*** *for expression change and* ***0.05*** *for significance.*

### 7. Disease Analysis

#### 7.1. Methods

For each disease, the number of differentially expressed (DE) genes annotated to a disease term is compared to the number of DE genes expected just by chance. iPathwayGuide uses an over-representation approach to compute the statistical significance of observing at least the given number of DE genes. The p-value is computed using the hypergeometric distribution as described for pORA in the Pathway Analysis section. This p-value is corrected for multiple comparisons using FDR and Bonferroni.

#### 7.2. Results

**Table 7.2.1: Top identified diseases**

| Disease Name | p-value | p-value (FDR) | p-value (Bonferroni) |
|---|---|---|---|
| Sphingolipidosis | **3.567e-4** | 0.208 | 0.234 |
| Osteoporosis | **7.892e-4** | 0.208 | 0.517 |
| Fanconi anemia | **9.524e-4** | 0.208 | 0.624 |
| Arrhythmogenic right ventricular cardiomyopathy | **0.002** | 0.255 | 1.000 |
| Benign familial infantile seizure | **0.002** | 0.255 | 1.000 |

##### Sphingolipidosis (H00423)

The sphingolipidoses are a group of monogenic inherited diseases caused by defects in the system of lysosomal sphingolipid degradation, with subsequent accumulation of non-degradable storage material in one or more organs. In this experiment, the algorithm identified **6** differentially expressed genes out of **9** genes associated with the disease.

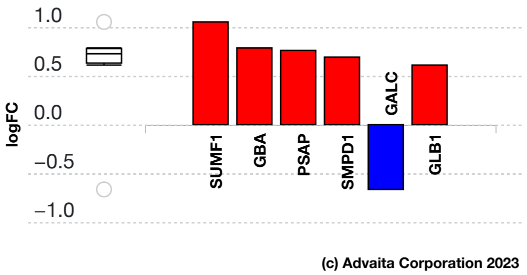


**Fig. 7.2.1: Gene measured expression bar plot:** *All the differentially expressed genes that are annotated to Sphingolipidosis are ranked based on their absolute value of log fold change. Upregulated genes are shown in red, downregulated genes are shown in blue. The box plot on the left summarizes the distribution of all the differentially expressed genes that are annotated to this disease. The box represents the 1st quartile, the median and the 3rd quartile, while the outliers are represented by circles.*

#### Osteoporosis (H01593)

Osteoporosis is a common disease characterised by a generalised reduction in bone mineral density (BMD), microarchitectural deterioration of bone tissue and an increased risk of fracture. Since BMD values fall progressively with age, the prevalence of osteoporosis increases with age. It has been estimated that approximately 50% of all women will have osteoporosis by the age of 80. Studies in twins and families indicate that genetic factors play an important role in the regulation of BMD and other determinants of osteoporotic fracture risk. Osteoporosis is a polygenic disorder, determined by the effects of several genes, each with relatively modest effects. Population-based studies and case-control studies have similarly identified polymorphisms in several candidate genes that have been associated with bone mass or osteoporotic fracture, including the vitamin D receptor, oestrogen receptor and collagen gene. Bisphosphonates, and in some patients denosumab, are first-line drugs for osteoporosis. In this experiment, the algorithm identified **6** differentially expressed genes out of **10** genes associated with the disease.

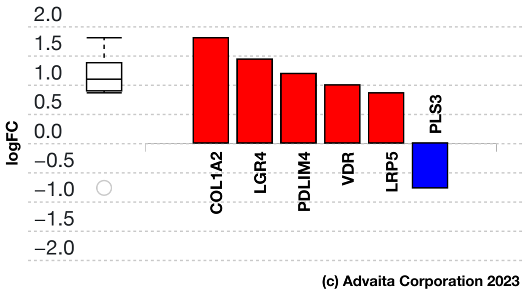


**Fig. 7.2.2: Gene measured expression bar plot:** *All the differentially expressed genes that are annotated to Osteoporosis are ranked based on their absolute value of log fold change. Upregulated genes are shown in red, downregulated genes are shown in blue. The box plot on the left summarizes the distribution of all the differentially expressed genes that are annotated to this disease. The box represents the 1st quartile, the median and the 3rd quartile, while the outliers are represented by circles.*

#### Fanconi anemia (H00238)

Fanconi anemia (FA), a recessive syndrome with both autosomal and X-linked inheritance, features diverse clinical symptoms, such as progressive bone marrow failures, chromosomal instability and susceptibility to cancer. To date, 13 FA gene products have been identified, which cooperate in a common DNA damage-activated signaling pathway regulating DNA repair (the FA pathway). In this experiment, the algorithm identified **9** differentially expressed genes out of **21** genes associated with the disease.

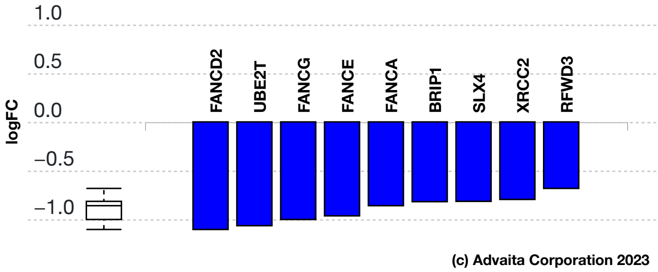


**Fig. 7.2.3: Gene measured expression bar plot:** *All the differentially expressed genes that are annotated to Fanconi anemia are ranked based on their absolute value of log fold change. Upregulated genes are shown in red, downregulated genes are shown in blue. The box plot on the left summarizes the distribution of all the differentially expressed genes that are annotated to this disease. The box represents the 1st quartile, the median and the 3rd quartile, while the outliers are represented by circles.*

#### Arrhythmogenic right ventricular cardiomyopathy (H00293)

Arrhythmogenic right ventricular cardiomyopathy (ARVC) is an inherited heart muscle disease that may result in arrhythmia, heart failure, and sudden death. The hallmark pathological findings are progressive myocyte loss and fibrofatty replacement, with a predilection for the right ventricle. A number of genetic studies have identified mutations in various components of the cardiac desmosome that have important roles in the pathogenesis of ARVC. Disruption of desmosomal function by defective proteins might lead to death of myocytes under mechanical stress. The myocardial injury may be accompanied by inflammation. Since regeneration of cardiac myocytes is limited, repair by fibrofatty replacement occurs. Several studies have implicated that desmosome dysfunction results in the delocalization and nuclear translocation of plakoglobin. As a result, competition between plakoglobin and beta-catenin will lead to the inhibition of Wnt/beta-catenin signaling, resulting in a shift from a myocyte fate towards an adipocyte fate of cells. The ryanodine receptor plays a crucial part in electromechanical coupling by control of release of calcium from the sarcoplasmic reticulum into the cytosol. Therefore, defects in this receptor could result in an imbalance of calcium homeostasis that might trigger cell death. In this experiment, the algorithm identified **5** differentially expressed genes out of **8** genes associated with the disease.

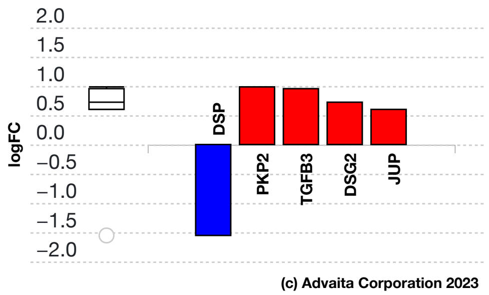


**Fig. 7.2.4: Gene measured expression bar plot:** *All the differentially expressed genes that are annotated to Arrhythmogenic right ventricular cardiomyopathy are ranked based on their absolute value of log fold change. Upregulated genes are shown in red, downregulated genes are shown in blue. The box plot on the left summarizes the distribution of all the differentially expressed genes that are annotated to this disease. The box represents the 1st quartile, the median and the 3rd quartile, while the outliers are represented by circles.*

#### Benign familial infantile seizure (H02362)

Benign familial infantile seizure (BFIS) is an autosomal dominant disease characterized by focal seizures, occurring mostly in clusters, and usually first seen between 4 and 8 months of life. Psychomotor development is normal, and seizures usually resolve within the first year of life. PRRT2 has been identified as the major gene found to be mutated in 80 to 90% of cases. Recently, mutations in the genes coding for the voltage-gated sodium channel subunits has been reported. In this experiment, the algorithm identified **3** differentially expressed genes out of **3** genes associated with the disease.

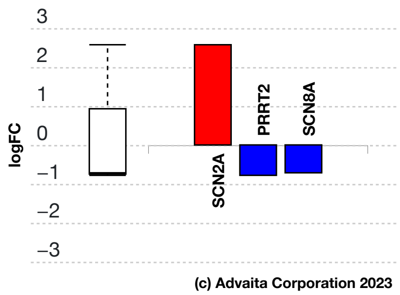


**Fig. 7.2.5: Gene measured expression bar plot:** *All the differentially expressed genes that are annotated to Benign familial infantile seizure are ranked based on their absolute value of log fold change. Upregulated genes are shown in red, downregulated genes are shown in blue. The box plot on the left summarizes the distribution of all the differentially expressed genes that are annotated to this disease. The box represents the 1st quartile, the median and the 3rd quartile, while the outliers are represented by circles.*

### 8. References


- Agarwal V, Bell GW, Nam J, Bartel DP. Predicting effective microRNA target sites in mammalian mRNAs. eLife, 4:e05005 (2015).
- Alexa, A., Rahnenfuehrer, J., Lengauer,T.: Improved scoring of functional groups from gene expression data by decorrelating GO graph structure. Bioinformatics 22(13): 1600-1607 (2006).
- Ashburner, M., Ball, C.A., Blake, J.A., Botstein, D., Butler, H., Cherry, J.M., Davis, A.P., Dolinski, K., Dwight, S.S., Eppig, J.T., Harris, M.A., Hill, D.P., Issel-Tarver, L., Kasarskis, A., Lewis, S., Matese, J.C., Richardson, J.E., Ringwald, M., Rubin, G.M., Sherlock, G.: The Gene Ontology Consortium. Gene ontology: Tool for the unification of biology. Nature Genetics 25(1): 25-9 (2000).
- Ashburner, M., Lewis, S.: On Ontologies for Biologists: The Gene Ontology - Untangling the web: 'In Silico' simulation of biological processes: Novartis Found Symp, 247:66-80; discussion 80-3, 84-90: 244-52 (2002).
- Benjamini, Y. and Hochberg, Y.: Controlling the false discovery rate: A practical and powerful approach to multiple testing. Journal of the Royal Statistical Society B, 57(1):289-300, (1995).
- Benjamini, Y. and Yekutieli, D.: The control of the false discovery rate in multiple testing under dependency. Annals of Statistics, 29(4):1165-1188, (2001).

- Bonferroni, C. E.: Il calcolo delle assicurazioni su gruppi di teste, chapter "Studi in Onore del Professore Salvatore Ortu Carboni", pages 13-60, Rome, (1935).
- Bonferroni, C. E.: Teoria statistica delle classi e calcolo delle probabilita. Pubblicazioni del Istituto Superiore di Scienze Economiche e Commerciali di Firenze, 8:3-62, (1936).
- Camon, E., Magrane, M., Barrell, D., Lee, V., Dimmer, E., Maslen, J., Binns, D., Harte, N., Lopez, R., Apweiler, R.: The Gene Ontology Annotation (GOA) database: sharing knowledge in Uniprot with Gene Ontology. Nucleic Acids Research, 32(Database issue), D262-D266 (2004).
- Davis AP, Grondin CJ, Johnson RJ, Sciaky D, McMorran R, Wiegers J, Wiegers TC, Mattingly CJ, The Comparative Toxicogenomics Database: update 2019, Nucleic Acids Research, 47(D1): D948-D954 (2019).
- Draghici, S., Khatri, P., Martins, R.P., Ostermeier, G.C. and Krawetz, S.A.: Global functional profiling of gene expression. Genomics, 81(2), pp.98-104 (2003).
- Draghici, S., Khatri, P., Bhavsar, P., Shah, A., Krawetz, S., Tainsky, M.A.: Onto-Tools, The toolkit of the modern biologist: Onto-Express, Onto-Compare, Onto-Design and Onto-Translate. Nucleic Acids Research, 31(13): 3775-81 (2003).
- Draghici, S., Khatri, P., Tarca, A.L., Amin, K., Done, A., Voichita, C., Georgescu, C., Romero, R.: A systems biology approach for pathway level analysis. Genome Research, 17(10): 1537-45 (2007).
- Draghici, S.: Statistics and Data Analysis for Microarrays Using R and Bioconductor, second edition. Chapman and Hall/CRC (2011).
- Friedman, R.C., Farh, K.K., Burge, C.B., Bartel, D.P.: Most mammalian mRNAs are conserved targets of microRNAs. Genome Research, 19: 92-105 (2009).
- Garcia, D.M., Baek, D., Shin, C., Bell, G.W., Grimson, A., Bartel, D.P.: Weak seed-pairing stability and high target-site abundance decrease the proficiency of lsy-6 and other miRNAs. Nature Structural & Molecular Biology, 18: 1139-1146 (2011).
- Gene Ontology Consortium. Creating the Gene Ontology Resource: Design and Implementation. Genome Research 11: 1425-1433 (2001).
- Gene Ontology Consortium. The Gene Ontology (GO) database and informatics resource. Nucleic Acids Research 32 (suppl 1): D258-D261 (2004).
- Griffiths-Jones S.: The microRNA Registry. Nucleic Acids Research 32:D109-D111 (2004).
- Griffiths-Jones S., Grocock R.J., van Dongen S., Bateman A., Enright A.J.: miRBase: microRNA sequences, targets and gene nomenclature. Nucleic Acids Research 34:D140-D144 (2006).
- Griffiths-Jones S., Saini H.K., van Dongen S., Enright A.J.: miRBase: tools for microRNA genomics. Nucleic Acids Research 36:D154-D158 (2008).
- Grimson, A., Farh, K.K., Johnston, W.K., Garrett-Engele, P., Lim, L.P., Bartel, D.P.: MicroRNA targeting specificity in mammals: Determinants beyond seed pairing. Molecular Cell, 27: 91-105 (2007).
- Fisher R. A.: Statistical methods for research workers. Oliver & Boyd, Edinburgh, (1925).
- Kanehisa, M., Goto, S.: KEGG: Kyoto Encyclopedia of Genes and Genomes. Nucleic Acids Research 28: 27-30 (2000).
- Kanehisa, M., Goto, S., Kawashima, S., and Nakaya, A.: The KEGG databases at GenomeNet. Nucleic Acids Research 30: 42-46 (2002).
- Kanehisa, M., Goto, S., Kawashima, S., Okuno, Y., and Hattori, M.; The KEGG resources for deciphering the genome. Nucleic Acids Research 32: D277-D280 (2004).
- Kanehisa, M., Araki, M., Goto, S., Hattori, M., Hirakawa, M., Itoh, M., Katayama, T., Kawashima, S., Okuda, S., Tokimatsu, T., and Yamanishi, Y.: KEGG for linking genomes to life and the environment. Nucleic Acids Research 36: D480-D484 (2008).
- Kanehisa, M., Goto, S., Furumichi, M., Tanabe, M., Hirakawa, M.: KEGG for representation and analysis of molecular networks involving diseases and drugs. Nucleic Acids Research 38: D355-D360 (2010).
- Kanehisa, M., Goto, S., Sato, Y., Furumichi, M., Tanabe, M.: KEGG for integration and interpretation of large-scale molecular datasets. Nucleic Acids Research 40: D109-D114 (2012).
- Kanehisa, M., Goto, S., Sato, Y., Kawashima, M., Furumichi, M., and Tanabe, M.; Data, information, knowledge and principle: back to metabolism in KEGG. Nucleic Acids Research 42: D199-D205 (2014).
- Khatri, P., Draghici, S., Tarca, A.D., Hassan, S.S., Romero, R.: A system biology approach for the steady-state analysis of gene signaling networks. Lecture Notes in Computer Science (LNCS) 4756, pp 32-41 (2007).
- Kozomara A., Griffiths-Jones S.: miRBase: integrating microRNA annotation and deep-sequencing data. Nucleic Acids Research 39:D152-D157 (2011).

- Kozomara A., Griffiths-Jones S.: miRBase: annotating high confidence microRNAs using deep sequencing data. Nucleic Acids Research 42:D68-D73 (2014).
- Lewis, B.P., Burge, C.B., Bartel, D.P.: Conserved seed pairing, often flanked by adenosines, indicates that thousands of human genes are microRNA targets. Cell, 120(1):15-20 (2005).
- Nam J, Rissland OS, Koppstein D, Abreu-Goodger C, Jan CH, Agarwal V, Yildirim MA, Rodriguez A, Bartel DP. Global analyses of the effect of different cellular contexts on microRNA targeting.Molecular Cell, 53:1031-43 (2014).
- Rhee, S.Y., Wood, V., Dolinski, K., Draghici, S.: Use and misuse of the gene ontology annotations. Nature Reviews Genetics 9(4):509-515 (2008).
- Szklarczyk, D., Morris, J.H., Cook, H., *et al*. The STRING database in 2017: quality-controlled protein-protein association networks, made broadly accessible. Nucleic Acids Research 45(D1):D362-D368 (2017).
- Tarca, A.L., Draghici, S., Khatri, P., Hassan, S., Mittal, P., Kim, J.S., Kim, C.J., Kusanovic, J.P., Romero, R.: A novel Signaling Pathway Impact Analysis (SPIA). Bioinformatics 25(1), 75-82 (2009).
